# Multiple latent clustering model for the inference of RNA life-cycle kinetic rates from sequencing data

**DOI:** 10.1101/2020.11.20.391573

**Authors:** Gianluca Mastrantonio, Enrico Bibbona, Mattia Furlan

## Abstract

We here propose a hierarchical Bayesian model to infer RNA synthesis, processing, and degradation rates from sequencing data, based on an ordinary differential equation system that models the RNA life cycle. We parametrize the latent kinetic rates, that rule the system, with a novel functional form, and estimate their parameters through 6 Dirichlet process mixture models. Owing to the complexity of this approach, we are able to simultaneously perform inference, clustering and model selection. We apply our method to investigate transcriptional and post-transcriptional responses of murine fibroblasts to the activation of proto-oncogene Myc. Our approach uncovers simultaneous regulations of the rates, which had not previously been observed in this biological system.

## 1. Introduction

RNA is one of the most important actors in the cellular biology context and it is involved, directly or indirectly, in any process that occurs inside a cell. This molecule is a cornerstone of the information-flow that subsists from DNA to proteins, due to both its role as a template for protein assembly and because of the involvement of non-coding RNAs in the regulation of gene expression levels (GELs) (e.g. modulation of transcript stability, protein synthesis and protein localisation) (Marchese <u>et al.</u>, 2017; Slack and Chinnaiyan, 2019; Vandevenne <u>et al.</u>, 2019). A cell constantly regulates the expression levels of thousands of genes, i.e. the number of the associated transcripts, in order to preserve its homoeostasis and adapt to the environment.

The RNA life-cycle in eukaryotic cells can be summarized in the following three steps: the synthesis of premature RNA molecules in the nucleus, their processing into mature transcripts (which includes exporting it to the cytosol), and mature RNA cytoplasmic degradation. The characterization of these mechanisms, which the cell exploits to modulate the amount of specific transcripts, according to internal and external stimuli, can provide exceptional insights into the biology of these responses. The investigation of the RNA life-cycle requires the experimental quantification of the GELs. The state-of-the-art approach used to perform this task is Next Generation RNA sequencing (RNA-Seq) (Goodwin <u>et al.</u>, 2016). Owing to the relevance of the topic, a remarkable number of public RNA-Seq datasets are currently available and easily accessible (e.g., the *Gene Expression Omnibus project* (Edgar <u>et al.</u>, 2002)), and a large amount of literature has been produced on the analysis of RNA-Seq data. In some papers, mixture models are used on the observed data to identify differences in GELs, see, for example, “RNA-Seq by expectation-maximization” (Li and Dewey, 2011), “Cufflinks” (Trapnell <u>et al.</u>, 2013), “Casper” (Rossell <u>et al.</u>, 2014), or the new approach developed by Papastamoulis and Rattray (2018). Some of these tools have also been used for the identification of genes that are differentially expressed under multiple experimental conditions, and this is one of the most common practices in the field (Trapnell <u>et al.</u>, 2013; Papastamoulis and Rattray, 2018). However, the mere quantification of expression levels is not enough to acquire a complete picture of the RNA life cycle, and the study of this datum alone could lead to misleading conclusions. Indeed, a cell can regulate gene expression through different fundamental processes. For example, an increase in the expression level of a gene between two conditions is usually interpreted as an intensification of its transcription, although the same observation could be due to a decreased degradation rate. The recent literature has shown that modeling the fundamental processes of the RNA life cycle and estimating the rates at which they occur can contribute greatly to a better understanding of the regulation mechanisms (Rabani <u>et al.</u>, 2011, 2014; Furlan <u>et al.</u>, 2019; Tesi <u>et al.</u>, 2019).

The RNA life cycle can be modeled as a system of linear ordinary differential equations (ODEs) that arises from a network of chemical reactions (see, for example, de Pretis <u>et al.</u>, 2015; Anderson and Kurtz, 2015). The time-dependent coefficients of such ODEs, the so-called kinetic rates (KRs), can be interpreted as the instantaneous rates at which the fundamental synthesis, processing and degradation mechanisms occur. In the last few years, several tools have been proposed to infer KRs from RNA-seq data, and of these, DRiLL (Rabani <u>et al.</u>, 2014) and INSPEcT (de Pretis <u>et al.</u>, 2015; Furlan <u>et al.</u>, 2020) provide a characterization of the full RNA life cycle. Motivated by a real data application, focused on the study of the activation of the proto-oncogene Myc in murine fibroblasts (de Pretis <u>et al.</u>, 2017), we propose a novel approach to the inference of the RNA life-cycle, cast in a Bayesian framework, that includes several clusterization steps.

To highlight the elements of novelty of our methodology, we here give a short summary of our contribution. We start by remarking that the experimental data are GELs, while our goal is to infer the three KRs, which are latent time and gene dependent functions. In our experimental setting, the cell regulates the expression of groups of genes to respond to a stimulus. In this common biological scenario, (Allocco <u>et al.</u>, 2004), the KRs are expected to take on several different but typical shapes. To infer such shapes, we define a single parametric family of functions that is sufficiently flexible to cover all of them. In this way, we recast our problem into the framework of parametric inference, and therefore do not need any a-posteriori model selection step. As a further layer of complexity, which has not yet been added to similar models, we define two distinct mixture models for each KR, that group genes with similar baselines or temporal responses to Myc activation into clusters. We decided to maintain these clusterizations independent, because genes with different/similar initial rates may respond to the treatment in similar/different ways.

The complexity of the model is such that certain devices are needed to make the implementation reasonably simple and, at the same time, the computational cost acceptable.

The most relevant source of complexity is that the six mixture models are defined over latent quantities; this is rare in the literature, since mixture models are generally defined over the observed data. Moreover, the emission distribution of the mixture models should be defined on the space where the KR parameters are identifiable. Unfortunately, due to the complexity of this space, such a distribution would be difficult to define and handle. Our solution is to define an approximated likelihood that allows us to use a Gaussian emission distribution over a suitable modified set of parameters, and to mimic the spike and slab mechanism for two of the parameters which, from an interpretative point of view, must be allowed to assume the value 0 with positive posterior probability.

Unlike other proposals, we estimate likelihood parameters, variance of the measurement error, the scaling factor, KR functions, and clusterings in a single Bayesian model. Moreover our proposal is able to extract and exploit information shared by the elements of the same cluster, thus resulting in better estimates of the parameters. We demonstrate the inferential gain provided by our approach, by detecting small but significant modulations of post-transcriptional rates (i.e. processing and degradation) which had not been appreciated in previous analyses of the same dataset (de Pretis <u>et al.</u>, 2015). Since Myc is a transcription factor, the synthesis rate is the most informative layer of regulation, and clusterization of its parameters provides groups of genes that are similarly involved in basic cellular processes, cancer-related processes, and in the RNA metabolism. Moreover, we manage to identify pervasive modulations of post-transcriptional rates, most likely due to either secondary regulations or the adaptation of processing and degradation to the transcriptional induction following Myc activation.

The paper is organised as follows. We start by describing the experiment used to study Myc activation, and the resulting dataset (Section 2). We then present the mathematical model we use to describe the RNA life cycle (Section 3) and the function we developed to parametrize the KRs (Section 3.3); we also discuss the solutions to some identifiability issues. We proceed by formalising the latent clustering models and their practical application to study Myc activation (Section 3.4). The final section of the paper (Section 4) regards a comparison of our novel Bayesian approach with other methods and a discussion of our results. We conclude with a critical summary of our work and some perspectives (Section 5). The online supplemental material (SM), available on the web page of the journal, contains additional figures.

## 2. The Data

Our dataset, taken from de Pretis <u>et al.</u> (2017), is organised as illustrated in Figure 1 A. It provides GELs (in Reads Per Kilobase Million, RPKM) of premature, mature, and nascent RNA for more than 10000 genes, at 11 time-points, for three replicas of the experiment. The experiment is designed to follow the activation of the transcription factor Myc in a murine fibroblast cell-line ( 3T9) over time. Myc plays a crucial role in the genesis and progression of tumors, and it is involved in the regulation of such basal cellular processes as differentiation, growth and proliferation (Dang, 2012; Chen <u>et al.</u>, 2018).

**Fig. 1.**
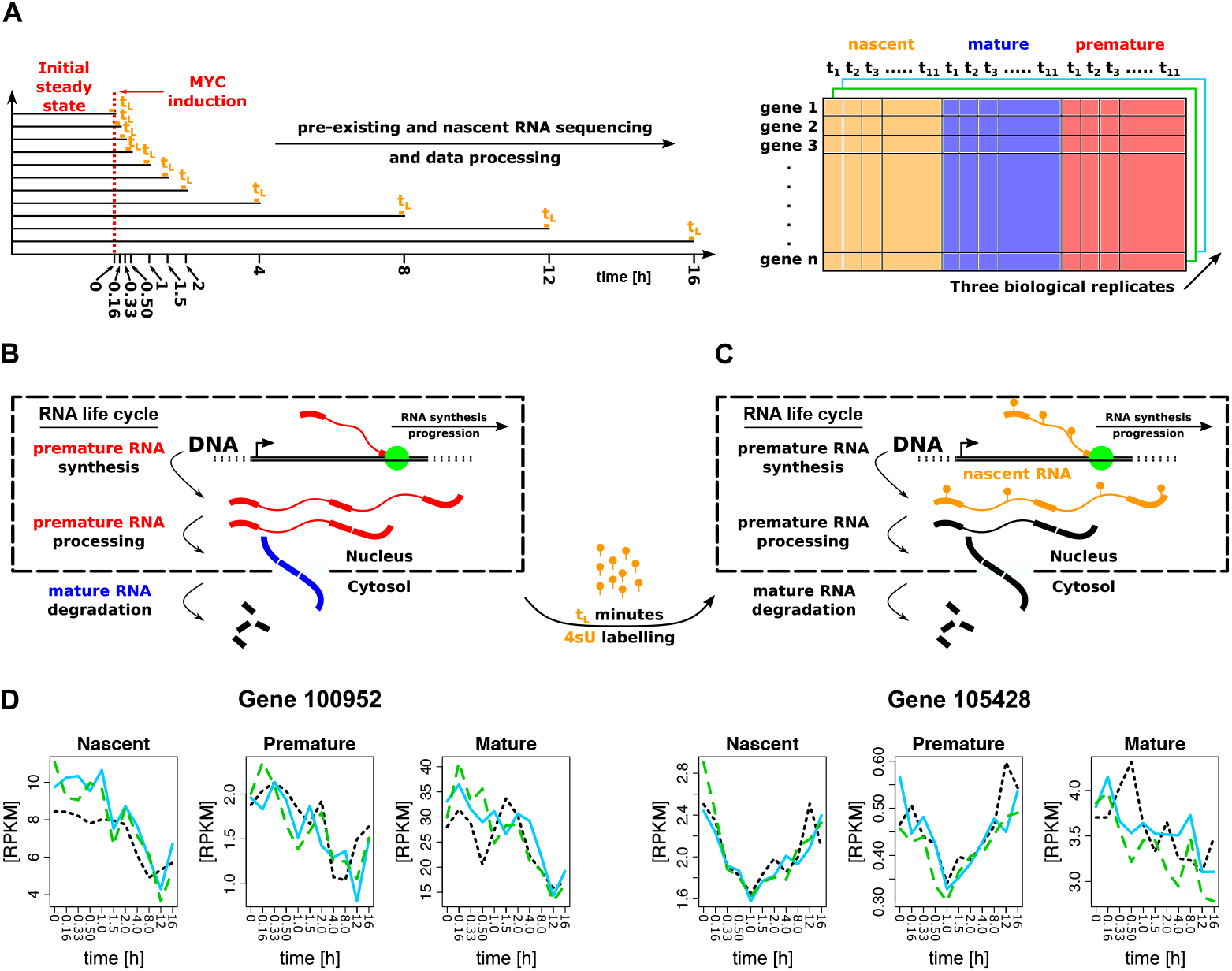
(A) Experimental design used for the study of Myc activation in 3T9 cells.(B) RNA life cycle in eukaryotic cells and definition of premature and mature RNA. (C) RNA metabolic labeling for nascent RNA quantification. (D) Gene expression profiles for two genes from the dataset; each replicate is represented by a different type of line.

The experiment starts with a population of cells, which is divided into multiple samples, in a stationary biological environment. Each sample is treated to induce Myc activation and, after a different time span, it is sequenced to quantify RNA expression levels. Myc activation is achieved through the expression of an artificial chimera (Littlewood <u>et al.</u>, 1995). This protein is natively inactive, and unable to perform any function, but it can rapidly be activated by adding the 4-hydroxytamoxifen (OHT) hormone to the cell culture medium. The authors performed standard (ribo-depleted) RNA-Seq, following Myc activation, through 11 time-points from an OHT treatment: 0h, 1/6h, 2/6h, 1/2h, 1h, 3/2h, 2h, 4h, 8h, 12h, 16h (Figure 1 A). Each experiment, performed on independent samples, was replicated three times, and resulted in expression levels of premature and mature RNA (Figure 1 B). The same experimental design was used to quantify nascent RNA through 4sU-Seq (Figure 1 C). In this case, an exogenous nucleotide (4-thiouridine or 4sU) is provided to the cells before sequencing for a fixed span of time (labeling time). 4sU is incorporated in the transcripts produced during the entire labeling time (nascent RNA) and is later exploited to physically separate them from the pre-existing RNA molecules. This portion of the transcriptome can be sequenced through standard RNA-Seq (Dölken <u>et al.</u>, 2008).

We focus on a set of 4909 transcriptional units, classified as Myc targets through a chromatin immunoprecipitation sequencing experiment, and altered in their kinetics.

However, it was not possible to analyze 12 transcriptional units, because they had negative expression levels. In the end, we retrieved a dataset of premature, mature and nascent RNA expression levels for 4897 genes in 3 replicates and for 11 time-points. Figure 1 D reports two examples from the dataset: the first one represents a typical transcriptional regulation, as can be seen from the adherence of the three profiles, while the second one may be associated with a more complex post-transcriptional scenario.

## 3. Methods

We provide a schematic representation of our Bayesian hierarchical model in Figure 2, in the form of a tree diagram whose elements are described in detail in the next sections and summarized hereafter.

**Fig. 2.**
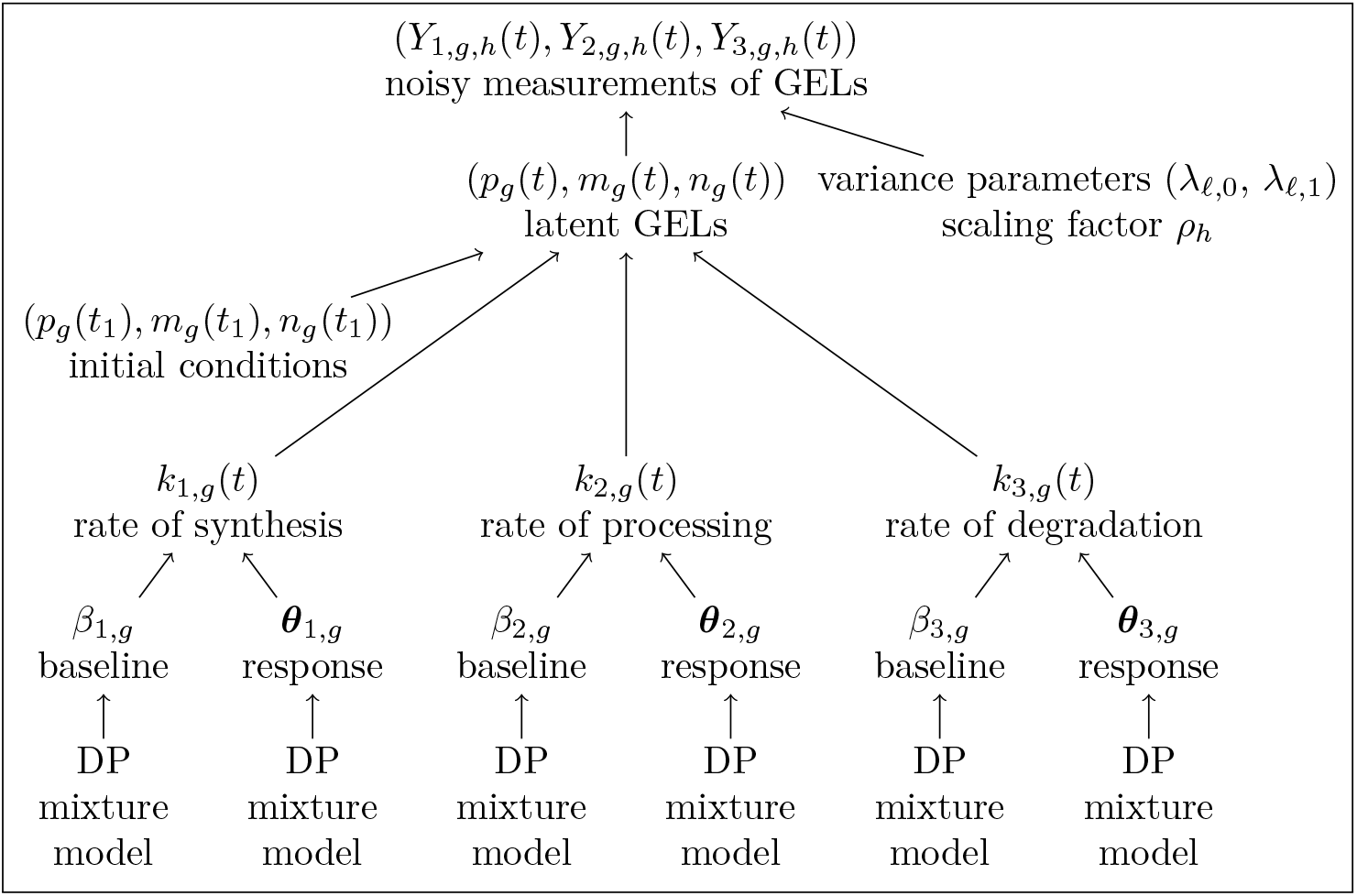
Graphical representation of the model structure.

For each gene *g* ∈ {1, … , *G*}, time-point *t* ∈ {*t*_1_, … , *t_T_*} and replica *h* ∈ {1, … , *H*}, *Y_ℓ,g,h_*(*t*) denote the *measured* GEL of premature RNA if *ℓ* = 1, mature RNA if *ℓ* = 2, and nascent RNA if *ℓ* = 3. These are noisy and scaled versions of the true unobserved values, which we indicate with *p_g_*(*t*), *m_g_*(*t*) and *n_g_*(*t*). In our dataset, *G* = 4897, *T* = 11, and *H* = 3. The distribution of the observed GELs depends on the parameters *ρ_h_*(*t*), *λ_ℓ,_*_0_, and *λ_ℓ,_*_1_, and is described in Section 3.1. The three *latent* GELs, (*p_g_*(*t*), *m_g_*(*t*), *n_g_*(*t*)), can be modeled as the solution of a system of ordinary differential equations that is specified by three time-dependent KRs *k_j,g_*(*t*), with *j* = 1, 2, 3. These KRs can be interpreted as the rates at which the fundamental mechanisms underlying the RNA life cycle, namely synthesis, processing and degradation, occur; see Section 3.2. A parametric family is introduced to define these KRs, see Section 3.3, and we show how the parameters can conveniently be subdivided into two sets, *β_j,g_* and ***θ**_j,g_*. The former is a scalar variable related to the baseline value of the KR before Myc activation, while the latter is composed of four parameters which characterize the shape of the response to the stimulus. We define for each of these parameters (*β_j,g_* and ***θ**_j,g_*), in each rate (*k_j,g_*(*t*)), a mixture model that groups genes with similar baselines (*β_j,g_*) or temporal responses to Myc activation (***θ**_j,g_*), into clusters. We decided to keep the clusterization on the three *β_j,g_* distinct from that of the three ***θ**_j,g_* because the relation between the basal value of the rates and their response to stimuli is not trivial, and, genes with different (or similar) initial rates may therefore respond to the treatment in similar (or different) ways.

### 3.1. The noise model

Since each *Y_ℓ,g,h_*(*t*) needs to be positive, for the noisy observed GELs we assume

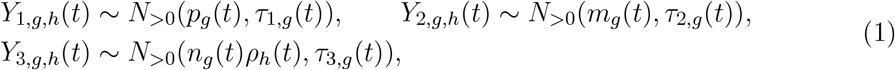

where *N*_>0_(·, ·) indicates a truncated normal distribution defined over ℝ^+^, and *ρ_h_*(*t*) is a scaling factor, that is required to normalize the nascent RNA libraries to the pre-existing RNA counterparts (Rabani <u>et al.</u>, 2011, 2014; de Pretis <u>et al.</u>, 2015). As shown in previous works, see, for example, Subramaniam and Hsiao (2012), the GELs affect both the mean and the variance of the measurement and we therefore assume the following:

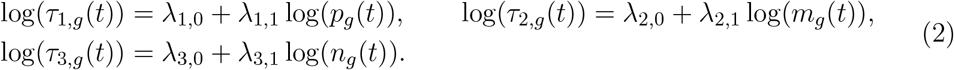

It should be noted that all the replicates of the expression levels of a gene share the same parameters, and, by introducing replicas we therefore increase the information on the latent GELs.

### 3.2. A mathematical model of the RNA life cycle

The life cycle of RNA molecules in eukaryotic cells, is divided into three sub-processes (Figure 1 B). The first one is the synthesis of premature RNA from DNA. This portion of the transcriptome is located inside the nucleus and it is not ready to perform its task (e.g. protein translation). Premature RNA requires structural modifications and/or exporting to the cytosol. This second stage of the RNA life cycle is named processing and its product is the mature RNA, which is eventually degraded by the cell (third step). The process may be described by the following network of chemical reactions

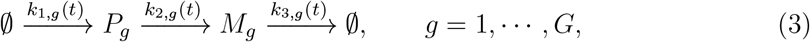

where *P_g_* and *M_g_* denote premature and mature RNA for gene *g*, respectively. The empty-set symbols are used to emphasize that premature RNA is synthesized from DNA without consuming any reactant, and mature RNA is subjected to degradation without forming any product. A system of ODEs that translates the reaction network (3) into mathematical terms is

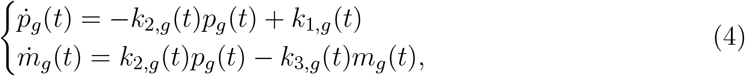

where the dots denote time derivatives. The effect of the processing is to decrease *p_g_*(*t*) and correspondingly increase *m_g_*(*t*) at rate *k*_2,*g*_(*t*). The degradation decreases *m_g_*(*t*) at rate *k*_3,*g*_(*t*), while the synthesis increases *p_g_*(*t*) at rate *k*_1,*g*_(*t*).

It is well known that, for the model described so far, it is difficult to identify all three KRs. Measurements of another variable, the so-called nascent RNA (Dölken <u>et al.</u>, 2008; Rabani <u>et al.</u>, 2011, 2014; de Pretis <u>et al.</u>, 2015), are usually included to ameliorate the identifiability, cf. Figure 1 C. Nascent RNA is defined as the amount of total RNA, premature plus mature, which is produced by the cell in a short span of *labeling* time *t_L_*, cf. Figure 1. By definition, the nascent RNA is absent at the beginning of the labeling time, and it is produced according to the same dynamics as the pre-existing counterpart during *t_L_*. However, the effect of degradation can be neglected in such a short span. The expression level of the premature 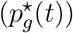 and mature 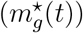 nascent RNA is therefore ruled by the following equations

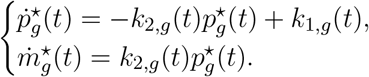

The sum 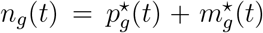 is the *nascent* RNA level. By summing the previous equations, one has that *n_g_*(*t*) only varies as a result of the effect of the synthesis:

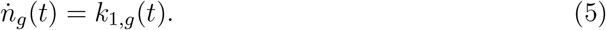

Since the time window for which nascent RNA evolves is short (*t_L_*), this rate can be considered approximately constant, and equation (5) can be integrated to obtain *n_g_*(*t*) = *k*_1,*g*_(*t*)*t_L_*, which is a third equation that has to be added to model (4) to facilitate the estimation of *k*_1,*g*_(*t*). It should be noted that the initial conditions (*p_g_*(*t*_1_), *m_g_*(*t*_1_)) need to be known to solve the ODE (4), and are here considered as further model parameters.

### 3.3. KR parametrization

In several biological experiments, a cell culture is perturbed and gene expression levels are repeatedly measured before and after the perturbation in order to understand which genes are involved in the response. In this case, the KRs typically vary over time and the typical shapes that we expect them to take on have the following characteristics: steady at the beginning of the experiment, steady after a long time from the perturbation, and varying (with some regularity) in the transient region between the two steady-states. Some typical shapes are: constant (some rates are not altered at all), monotonic (both increasing and decreasing), and peak-like functions. They have already been successfully applied to describe transcriptional and post-transcriptional responses in several biological systems (see, for example, Chechik and Koller, 2009; Rabani <u>et al.</u>, 2011, 2014; de Pretis <u>et al.</u>, 2015). More complex patterns would be difficult to identify with the number of time-points and replicates that are usually available.

We introduce a unique parametric family of functions which, for different values of the parameters, can cover all such characteristic shapes. Let *φ*(·|*μ*, | *σ*^2^) be a Gaussian density with mean *μ* ∈ ℝ and variance *σ*^2^ ∈ ℝ^+^. We define the family of functions *f* (*t* | *μ, σ*^2^, *κ*_−∞_, *κ_μ_*, *κ*_+∞_) to which all the KRs belong in the following way:

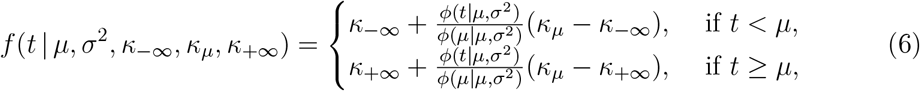

where *κ*_−∞_, *κ_μ_* and *κ*_+∞_ all belong to ℝ^+^. Function *f* in equation (6) is obtained by applying different scalings and vertical translations of a Gaussian density to its right and left halves, with respect to the mean value *μ*, taking care to preserve continuity at time-point *t* = *μ* (Figure 3). It is easy to see that:

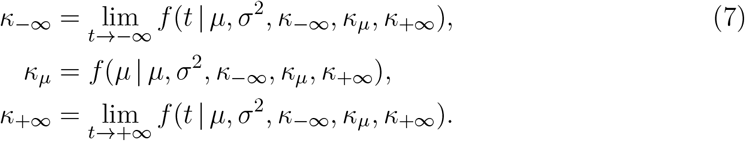

For easiness of interpretation, we split and rename the parameters as follows. First, we single out *κ*_−∞_ and we rename it *β* to simplify the notation. Unlike the other parameters, which are related to the response, *β* is the baseline level, i.e. the initial steady-state, and it is analyzed separately. Secondly, we introduce the logarithmic ratios

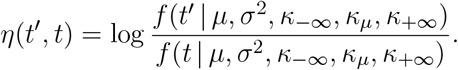

**Fig. 3.**
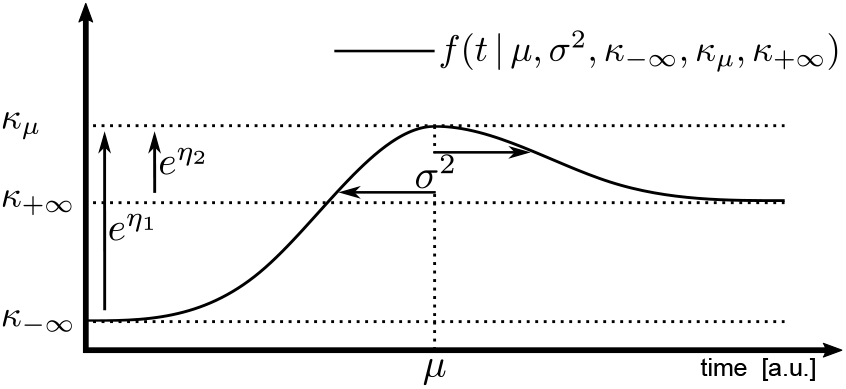
Graphical representations of the KR parametrization. It should be noted that 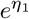 and 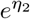 are shown to indicate the section of the function they determine, but they are not equal to the length of the arrows, see equation (8).

These quantities are called *log-fold-change*s in computational biology and are usually used to measure modulations with respect to the baseline level *f* (*t* | *μ, σ*^2^, *κ*_−∞_, *κ_μ_*, *κ*_+∞_), and we therefore define

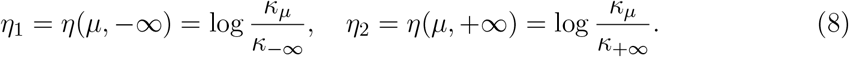

Parameters *μ*, *σ*^2^, *η*_1_, *η*_2_ are all related to the characterization of the response to perturbations. In particular, *μ* and *σ*^2^ characterize the *temporal* location and duration of the response, while *η*_1_ and *η*_2_ determine the typical shape, as highlighted in Table 1. We collect these four parameters in a single vector that we denote ***θ*** to obtain a more compact notation. Examples of the forms that can be obtained with (6), by changing its parameters, are shown in Table 1, where we can see that all the standard shapes (constant, increasing/decreasing, peak-like) are possible. The family of functions (7) can now be re-parametrized as *f* (*t* | *β*, ***θ***), with *β* = *κ*_−∞_ and ***θ*** = (*μ, σ*^2^, *η*_1_, *η*_2_).

**Table 1.**
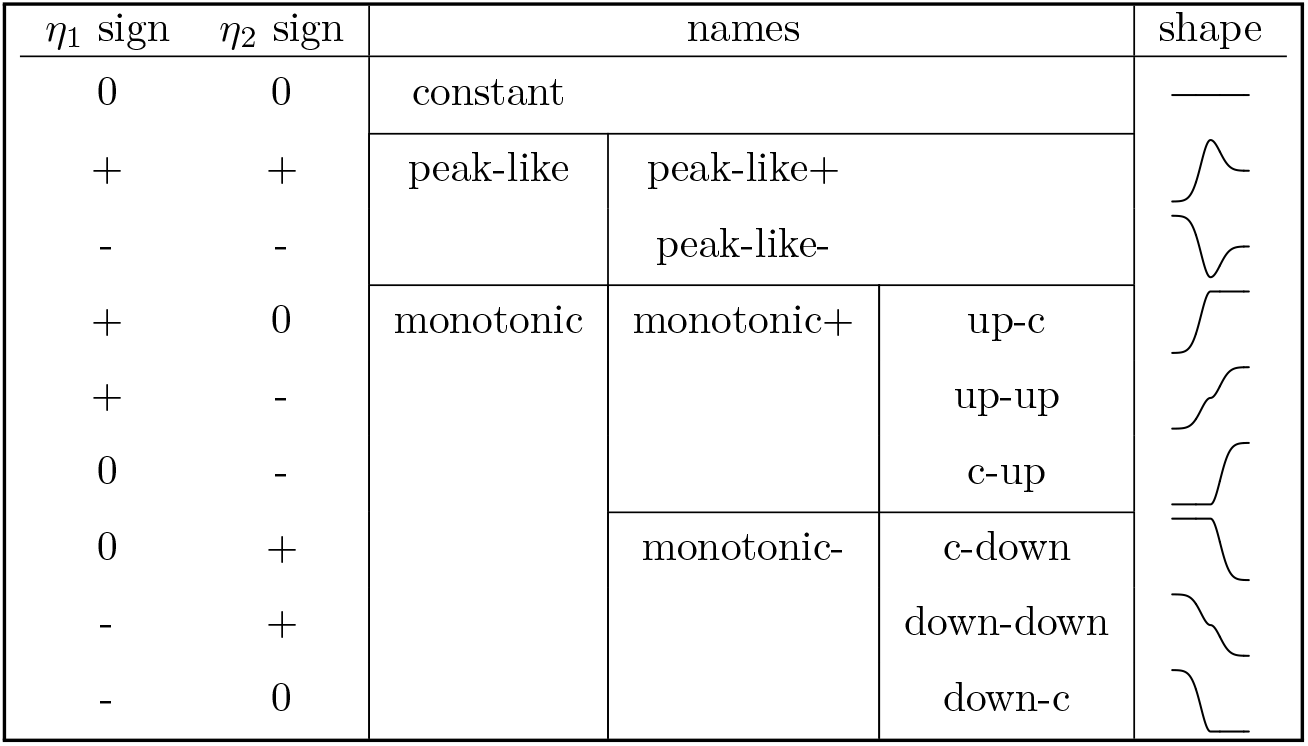
KR shapes as functions of the log-fold-changes and the names we use to describe the shapes.

| $\eta_1$ sign | $\eta_2$ sign | names | | | shape |
| --- | --- | --- | --- | --- | --- |
| 0 | 0 | constant |  |  | — |
| + | + | peak-like | peak-like+ |  |  |
| - | - |  | peak-like- |  |  |
| + | 0 | monotonic | monotonic+ | up-c |  |
| + | - |  |  | up-up |  |
| 0 | - |  |  | c-up |  |
| 0 | + |  | monotonic- | c-down |  |
| - | + |  |  | down-down |  |
| - | 0 |  |  | down-c |  |

Although the family of functions (7) is well defined for all real values of *μ*, *η*_1_, and *η*_2_, and for positive values of *σ*^2^, certain identifiability and interpretability issues may arise if some conditions are not met. For example, if *μ* is smaller than the first observed time *t*_1_, and *σ*^2^ is small (compared to |*μ* − *t*_1_|), function *f* in the interval [*t*_1_, *t_T_*], is indistinguishable for any arbitrary choice of *η*_1_ and *η*_2_ from a constant one, which should instead be given by *η*_1_ = *η*_2_ = 0 (Table 1). For this reason, identifiability constraints are needed. One main requirement is that the value of the function (6) at time-points *t*_1_ and *t_T_* should be “close” to *β* and *κ*_∞_, respectively, which means that the most relevant part of the function graph lies within the observed time window. Hence, for peak-like shapes (*η*_1_*η*_2_ > 0, cf. Table 1), we require

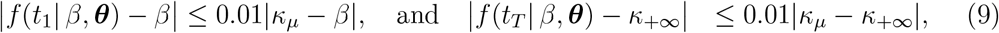

which implies two conditions:

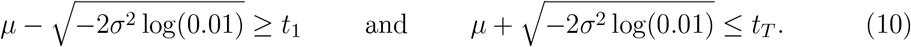

The value 0.01, or any other value with comparable magnitude, ensures that the derivative of our function is almost zero at time *t*_1_ and *t_T_* (it is ≈ 0.01 in (10)) and then *f* (*t*_1_ | *β*, ***θ***) ≈ *β*, which lets us interpret *β* as the steady-state at time *t*_1_, and *f* (*t_T_* | *β*, ***θ***) ≈ *κ*_∞_, which ensures a steady-state at the end of the observed window. Considering a value that us too small would restrict the possible values of (*μ, σ*^2^), without any significant improvement in the steady-state approximation, while such an approximation may no longer be acceptable for a value that is too large. It should be noted that one of the log-fold-changes vanishes for monotonic shapes; however, we can still derive an identifiability condition, provided that (9) holds, e.g. when *η*_1_ = 0, we have a c-up or c-down shape, and *κ_μ_* = *β*. Since the function is constant from *t*_1_ to *μ*, the first equation of (9) holds if, and only if, *t*_1_ ≤ *μ*. The conditions we obtain for c-up and c-down shapes (*η*_1_ = 0) are therefore

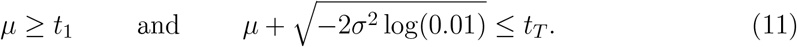

Similarly, for up-c and down-c shapes (*η*_2_ = 0), we obtain

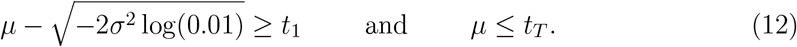

The monotonic up-up (down-down) is an intermediate shapes between c-up and up-c (c-down and down-c). Hence, the limits on *μ* should be close to (11), if *η*_1_ < *η*_2_, and close to (12), if the opposite is true. Therefore, we define the constraints on *μ* as:

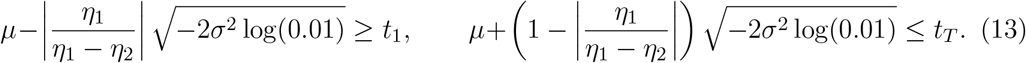

The subset of the parameter space where identifiability holds is denoted as 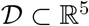, and is defined by the condition *β* > 0 and one of inequalities (10), (11), (12) or (13), depending on the signs of *η*_1_ and *η*_2_, which are instead unconstrained real numbers. Although identifiability is only granted in 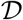, we prefer not to force the parameter to belong to this set for easiness of implementation, and, as we explain in the next section, we introduce an approximated likelihood that gives very little support to values outside 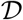.

### 3.4. Latent clustering models

It is biologically reasonable to allow groups of genes that have a similar baseline *β_j,g_* level to respond differently to a perturbation (therefore with different ***θ**_j,g_*). Moreover, even though two genes have a similar rate of synthesis, it is not necessarily true that the other rates are similar. For this reason, we separately introduce mixture models, based on the DP, to the bottom level of the model hierarchy, for the parameters *β_j,g_* and ***θ**_j,g_* for each *j* ∈ {1, 2, 3}, see Figure 2. Making inference on such hidden mixtures is particularly hard and makes the implementation of the MCMC more involved. For this reason, it is necessary to ensure that the sampling of the DP is as straightforward as possible, and the use of Gibbs updates for the mixture parameters is preferable. Moreover, we also want to ensure that the model can discriminate between the possible shapes of *k_j,g_*(*t*), and that our MCMC algorithm does not become trapped in a region of the parameter space where such shapes are not identifiable. To this aim, the most natural choice would be to adopt a distribution with support in 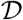 for (*β_j,g_, **θ**_j,g_*); however, this choice is impractical, since is not possible to find an easy-to-handle distribution over this complicated domain. The solution we propose here is to let (*β_j,g_, **θ**_j,g_*) be defined over the whole ℝ^5^ and to modify the data likelihood (equation (1)) with an alternative one that gives an almost null posterior support to all the parameter values outside 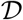. Our proposal is:

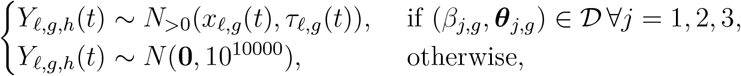

where *x*_1,*g*_(*t*) = *p_g_*(*t*), *x_ℓ,g_*(*t*) = *m_g_*(*t*) and *x*_3,*g*_(*t*) = *n_g_*(*t*)*ρ_h_*(*t*). By doing so, we enforce the posterior distribution to have a very small mass outside 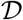, which means that the parameters in the unidentifiable region are almost never sampled in the MCMC algorithm. A second issue is that if the marginal distribution of the log-fold-change *η*_1,*j,g*_ is continuous, the posterior probability to be equal to 0 vanishes, which means that we are not able to estimate a constant *k_j,g_*(*t*) (or c-up, c-down, up-c, and down-c shape). One possible solution is to use a distribution that is continuous over (−∞, 0) ∪ (0, ∞) and has a point mass at 0. We can do this by introducing the new variables 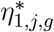 and 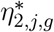, related to *η*_1,*j,g*_ and *η*_2,*j,g*_ through the following:

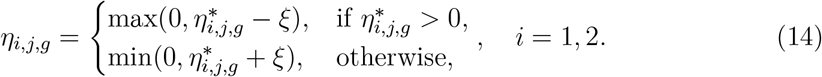

If we assume a continuous distribution for 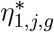, as a result of (14), the distribution over *η*_1,*j,g*_ is continuous over (−∞, 0) ∪ (0, ∞) and has a point mass at 0 equal to the cumulative distribution of 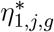 between −*ξ* and *ξ*. A similar result holds for *η*_2,*j,g*_. Equation (14), therefore defines a spike-and-slab distribution for *η*_1,*j,g*_ and *η*_2,*j,g*_ (Ishwaran and Rao, 2005), which has a spike (point mass) at 0 and a slab (continuous distribution) over ℝ. However, our approach allows us to work with the continuous “latent” variables 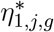 and 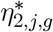, which makes the implementation of the MCMC easier. It should also be noted that the posterior probability mass on *η_i,j,g_* = 0 and *η*_2,*j,g*_ = 0 depends on their marginal distributions, e.g., if the mean is zero and the variance is small/large, the mass is close to 1/negligible. As shown in equation (15), the means and variances of these distributions are parameters that are inferred from the model fitting and, more importantly, they are cluster dependent. Therefore, also the posterior probability mass on *η*_1,*j,g*_ = 0 and *η*_2,*j,g*_ = 0 is cluster dependent.

We can now work with the parameters *β_j,g_* and 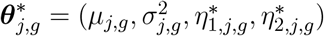, which are all defined over ℝ. We then define 6 mixture models, based on Gaussian densities, 3 of them over *β*_1,*g*_, *β*_2,*g*_ and *β*_3,*g*_ and the others over 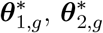 and 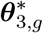. In other words, the models are DP Gaussian mixtures (Neal, 2000) formalized as:

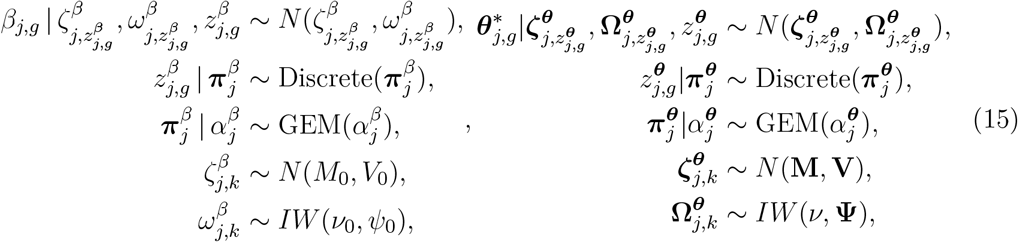

where *k* ∈ ℕ. In models (15), 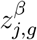 and 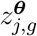 are the discrete random variables which represent the labels which identify the component of the mixture to which the parameters belong. These variables are assumed to come from a discrete distribution, whose probabilities follow a Dirichlet Processes defined by the GEM (or stick-breaking) distribution (Gnedin <u>et al.</u>, 2001). Given the allocation variables 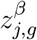 and 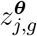, parameters *β_j,g_* and 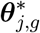 are normally distributed. All the random quantities in model (15) can easily be updated in the MCMC algorithm using only Gibbs steps, thereby facilitating the implementation of the model.

## 4. Real data application

The results obtained by means of our model on the real data application are presented in this section. Before discussing these results, we offer some implementation details to explain how we process the model output to obtain the statistics used to interpret the results.

### 4.1. Implementation details

#### Priors

We choose weakly informative priors for all the model parameters. We set **M** = **0**, **V** = 100**I**, *M*_0_ = 0, *V*_0_ = 100, **Ψ** = **I**, *ν* = 5, *ν*_0_ = 2, and 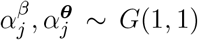 for the DP components of our proposal (equation (15)). We choose a normal distribution *N* (0, 100) for the variance parameters *λ_ℓ,_*_0_ and *λ_ℓ,_*_1_ (equation (2)), and we select a Gamma *G*(1, 1) for the scaling factor *ρ_h_*(*t*) (equation (1)). A uniform *U* (0, 10000) is used as a prior for the initial conditions *n_g_*(*t*) and *m_g_*(*t*) (equation (4)) of the ODE system, where 10000 is the maximum level we can expect to see. We also assume *ξ* = 10 but, through preliminary simulation experiments, we verified that if *ξ* < 20, and considering the given priors, its value has almost no influence on the posterior inference.

#### Implementation

The model is implemented in R/C++ (R Core Team, 2020) and uses OpenMP (OpenMP Architecture Review Board, 2008) for parallel computing. We use the sampling scheme proposed by Escobar and West (1995) for the DP hyper-parameters 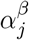 and 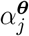. We marginalize with respect to the state probabilities 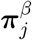 and 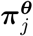, and we use the algorithm number 8 of Neal (2000) (with parameter *m* = 1) at each MCMC iteration and for each mixture model, to perform a global update of the latent allocation variables 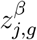 and 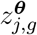, and a single step of the “restricted Gibbs sampling split-merge” of Jain and Neal (2007) (with parameters (0,1,0,0)), to split and merge existing states. Both updates are easily implemented since, given the chosen priors, the means and variances of the normal densities can easily be sampled. Parameters 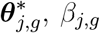, and sets 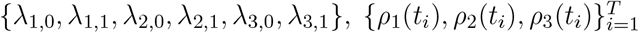, and {*p_g_*(*t*_1_), *m_g_*(*t*_1_)} are all sampled separately using Metropolis steps with the adaptive proposals of Andrieu and Thoms (2008); algorithm number 4. The ODE solutions are computed with the *odeint* C++ library (Ahnert and Mulansky, 2011). The code that implements the methodology is available at https://github.com/GianlucaMastrantonio/Multiple_latent_clusterisation_model_for_the_inference_of_RNA_life_cycle_kinetic_rates.

#### Posterior samples

We estimate the model on a computer cluster with 32 cores and using a stop-and-go strategy to check whether and when convergence has been achieved. In detail, we run the model using batches of 250000 iterations, burnin 175000, and thin 30. We check convergence after each batch using split 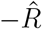 statistics (Gelman <u>et al.</u>, 2013) computed on parameters 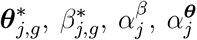, *λ*_ℓ,0_, *λ*_ℓ,1_, *n_g_*(*t*), *m_g_*(*t*), and *ρ_h_*(*t*) and, if not all of them have a lower split 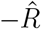 than 1.05 (our selected threshold), we run another batch with the last simulated parameters of the previous MCMC batch as the starting values. We use random values for the first batch. After 9 batches of simulations, all the parameters were within the split 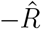 threshold and only then, did we use a 10-th batch of simulations to obtain the posterior samples we use to describe the results. Hence, the model required 250000 × 10 = 2500000 iterations, with a total burnin of 250000 × 9+175000 = 2425000, thin 30, having than 2500 posterior samples; the computations took 20 days.

#### Output processing

Using the algorithm of Wade and Ghahramani (2018), we find a representative point estimate of the cluster membership variables 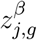 and 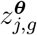, which we indicate with 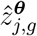 and 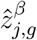, while we use the posterior means 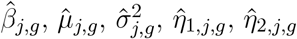 as representative values of the KR parameters. Since the posterior probability that a KR has one of the 5 shapes defined in Table 1 depends on whether *η*_1,*j,g*_, *η*_2,*j,g*_ are positive, negative or null, we can associate the representative shape as the one with the highest posterior probability to each KR.

### 4.2. Comparison of the multiple inference method

We compare the performance of our model (M1) with a simplified version (M2), which assumes 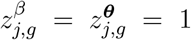 for all *j* and *g* (i.e. the mixture has a single component). We use the same priors as M1 for the estimation of M2, and the same strategy to check for the MCMC convergence. We use the Deviance Information Criterion 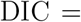 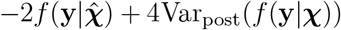 (Gelman <u>et al.</u>, 2014) to compare the models, where *f* (**y** | ·) is the data likelihood given by the distributions in (1), ***χ*** is the vector of the likelihood parameters computed using the posterior means of our model, and Var_post_(·) is the variance with respect to the posterior distribution. The value of DIC is 209400 for M1 and 257268 for M2, thus showing that our proposal is preferable. Since our approach can be though of as a Bayesian extension of the frequentist framework implemented in INSPEcT (M3), a comparison between these two methods would also be interesting, despite the absence of obvious ways of comparing Bayesian and frequentist approaches in the literature. One possible solution is to use cross-validation indices, but this would require fitting the models multiple times which, given the significant computational cost of our proposal, would be prohibitive. We approximate the cross-validation predictive distribution for a given gene/replica/time-point with the standard predictive distribution, and we use the continuous ranked probability score (CRPS) (Gneiting and Raftery, 2007) for the comparison. CPRS is often used to assess the accuracy of probabilistic models, and it is defined as the mean square error between the predicted and empirical cumulative distribution functions:

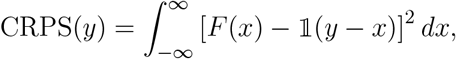

where *F* (·) is the predictive distribution and 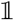 is the Heaviside step function. We estimate the CRPS for each of our observations and, for comparison purposes, we compute the fraction of data points with a lower index for one model than another one; we decided not to average across all the data points because they are not identically distributed. These comparisons are recapitulated in Table 4.2. Further insight can be gained by computing pairwise Pearson’s correlations between the CRPS indexes estimated for premature, mature and nascent RNA for each model. The correlations are between 0.34 and 0.62, and interestingly, they are always higher for M1 and M2, than for M3 (0.52, 0.52 and 0.38 on average, respectively, Figure 37-SM). It follows that the Bayesian approaches tend to fit all the RNA species profiles with a similar goodness, while the frequentist one generally recapitulates the profile of one experimental quantity better than the others. This is not desirable in the current application scenario, since any RNA species is potentially equally informative about the underlying regulations of the RNA metabolism.

As a result of the way we compute the CRPS, i.e. without using holdout observations, its value may be influenced by overfitting. Hence, we also decided to compare M1 and M3 focusing on the different features and interpretability of the estimated KRs, showing that our proposal (M1) gives a better and clearer description of the data. Therefore we report the proportions of constant and variable rates in Table 3, both increasing (monotonic+ and peak+) and decreasing (monotonic− and peak−) for each model. Since Myc is a transcription factor, we expect a primary response at the synthesis level. This is in particular the case for M1 and M3, which have less than 2% of the genes constant in synthesis. The percentage is higher for M2 (10%). Nevertheless, *k*_1_ is still by far the most variable rate. We can also observe an agreement about the direction of the modulations for all the inference methods which predict a higher percentage of repressed genes than induced ones (0.49 – 0.63 versus 0.36 – 0.42). The inferred regulations are more different for the post-transcriptional rates; indeed, M3 predicts a higher percentages of constant *k*_2_ and *k*_3_ (71% and 80%, respectively) than the Bayesian approaches (between 46% and 70%). This means that M1 and M2 tend to explain the expression more as a simultaneous regulation of all the three kinetic rates than M3 (Figure 4 and 38-SM). This choral regulation, which was not observed to the same extent in past analyses (M3), is biologically reasonable. Indeed, the role of synthesis remains prevalent in shaping the Myc response, but indirect and less intense post-transcriptional regulations can also be expected (see, for example, McManus <u>et al.</u>, 2015). We conclude that INSPEcT fails to capture the more complex regulatory scenarios, which are instead inferred by means of our novel Bayesian method.

**Table 2.** Fraction of the CRPS values which satisfy the condition reported at the top of the grid.

| | CRPS M1 ≤ CRPS M2 | CRPS M1 ≤ CRPS M3 | CRPS M2 ≤ CRPS M3 |
| --- | --- | --- | --- |
| premature | **0.650** | 0.517 | 0.489 |
| mature | **0.653** | 0.517 | 0.501 |
| nascent | **0.643** | 0.528 | 0.508 |

**Table 3.** For each model and function *k*, we indicate the fraction of constant functions, decreasing monotonic or peak-like functions (monotonic− and peak−), and increasing monotonic or peak-like functions (monotonic+ and peak+)

|  |  | constant | peak+ or monotonic+ | peak- or monotonic- |
| --- | --- | --- | --- | --- |
| M1 | $k_{1,\cdot}(\cdot)$ | 0.019 | 0.415 | 0.566 |
| | $k_{2,\cdot}(\cdot)$ | 0.515 | 0.342 | 0.143 |
| | $k_{3,\cdot}(\cdot)$ | 0.460 | 0.063 | 0.476 |
| M2 | $k_{1,\cdot}(\cdot)$ | 0.100 | 0.408 | 0.492 |
| | $k_{2,\cdot}(\cdot)$ | 0.641 | 0.198 | 0.161 |
| | $k_{3,\cdot}(\cdot)$ | 0.703 | 0.150 | 0.147 |
| M3 | $k_{1,\cdot}(\cdot)$ | 0.013 | 0.357 | 0.630 |
| | $k_{2,\cdot}(\cdot)$ | 0.712 | 0.081 | 0.207 |
| | $k_{3,\cdot}(\cdot)$ | 0.801 | 0.118 | 0.081 |

**Fig. 4.**
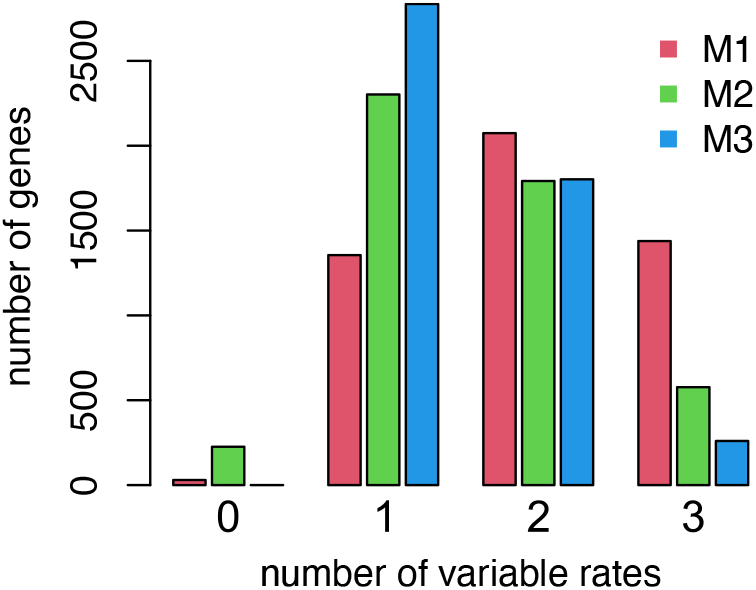
Number of genes (y-axis) with a given number of non-constant rates (x axis) for M1, M2 and M3.

### 4.3. Myc response analysis

In this section, we analyze the results inferred by means of our model. Each gene and KR is associated with a cluster, and we comment on all the sets of genes returned by our procedure for the processing and degradation rates. We instead focus on the clusters with at least 50 elements for the synthesis rate. This constraint is introduced to filter small gene sets (less than 200 genes in total) with modest responses and/or heterogeneity, for the sake of clarity and space, even though the number of elements is not necessarily a proxy of the biological relevance of a cluster. The missing clusters are depicted in Figure 3-SM and Figure 4-SM. In the first column of Figures 5 and 6, we show heatmaps of the temporal behavior of the KR log-fold-changes that belong to each cluster, computed using the posterior mean values 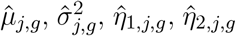, of the KR parameters. In the second column, for each time-point, we show the boxplots that represent the distributions of the KR log-fold-changes computed in the first column. The third and fourth columns present the 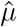 versus 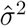 and 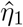 versus 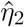 smoothed color density representations of the scatterplots. Both of these graphs provide valuable information about the shape of the responses in the cluster of interest. Figure 6 contains analogues graphs for the processing and the degradation rates.

**Fig. 5.**
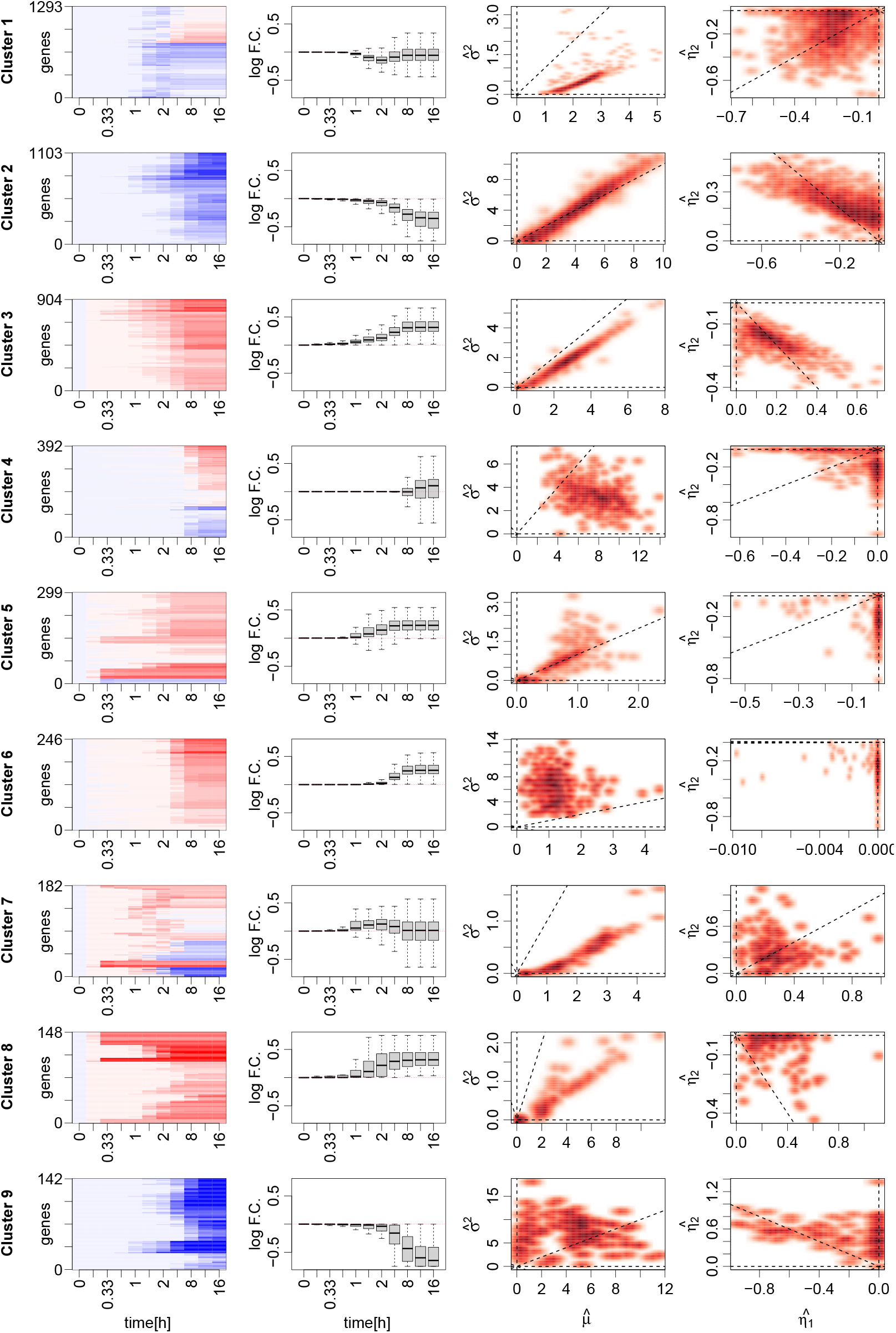
Synthesis rate modulations in response to Myc activation for 7 clusters that are composed of at least 50 genes. (First column) Heatmaps showing the log-fold-changes of the rate 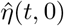, compared to the first time-point, for each gene in the cluster (values saturated at ±0.75). (Second column) Boxplots showing the distribution of the synthesis rate log-fold-changes, compared to the first timepoint (values saturated at ±0.75). (Third column) 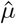 versus 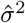 smooth-scatter plot. (Fourth column) 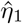 versus 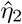 smooth-scatter plot. The dashed lines represent the horizontal and vertical axes and the bisector of the first and third quadrants.

**Fig. 6.**
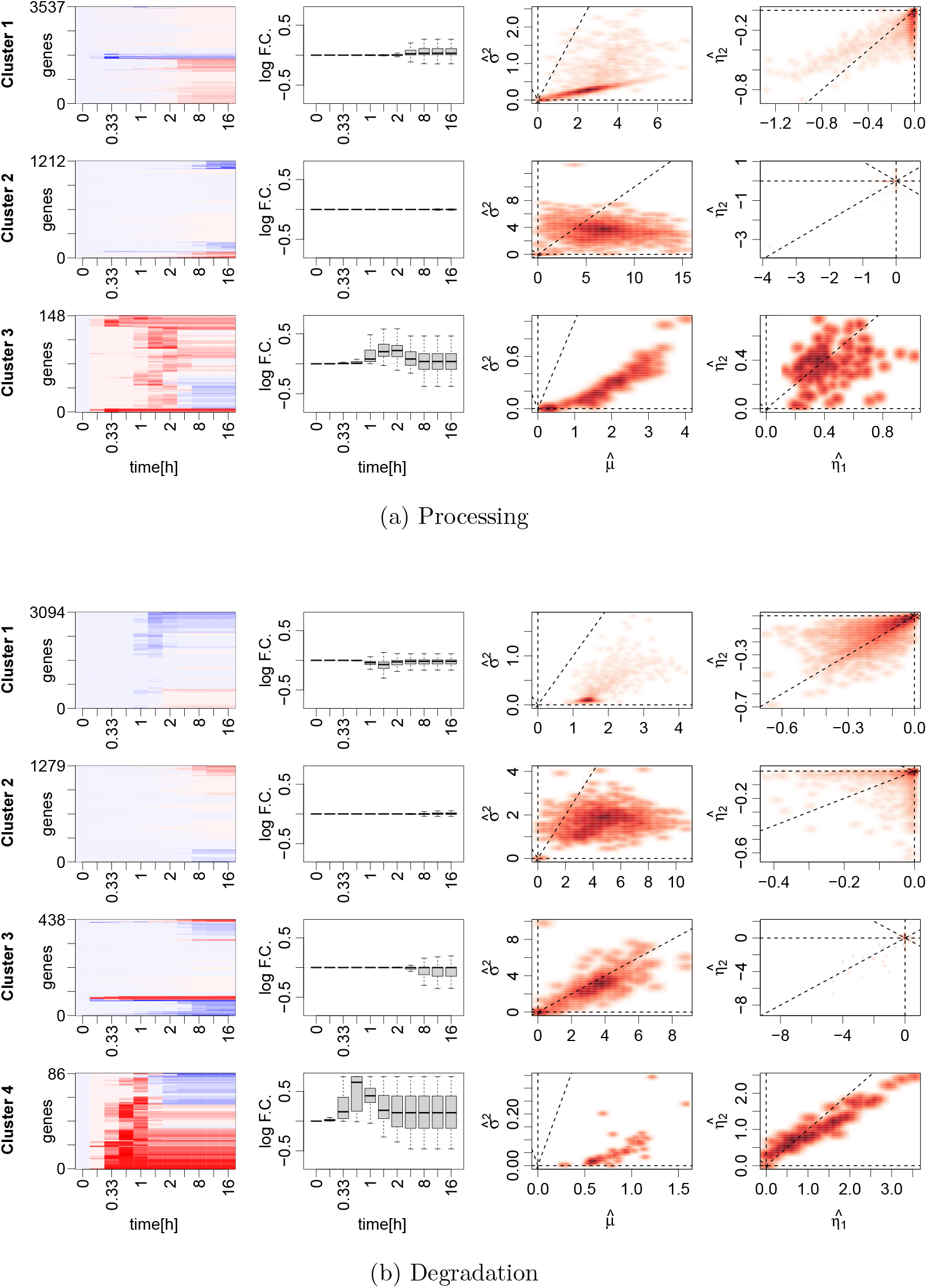
Processing (a) and Degradation (b) rate modulations in response to Myc activation. (First column) Heatmaps showing the log-fold-changes of the rate 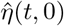, compared to the first timepoint, for each gene in the cluster (values saturated at ±0.75). (Second column) Boxplots showing the distribution of the rate log-fold-changes, compared to the first time-point (values saturated at ±0.75). (Third column) 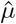 versus 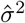 smooth-scatter plot. (Fourth column) 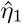 versus 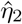 smoothscatter plot. The dashed lines represent the horizontal and vertical axes and the bisector of the first and third quadrants.

In order to support the discussion, we perform enrichment analyses, based on GO annotations (Dessimoz and Škunca, 2017; The Gene Ontology Consortium, 2019), for each rate. The graphical results of these analyses are shown in the supplemental material, with the exception of Figure 7, which here is reported as an example of a full output and refers to a specific case which has been selected both because of its interest and its graphical clarity. The GO enrichment analysis exploits a set of terms which, in our case, are pertinent to biological processes (e.g. “regulation of mRNA stability” or “cellular response to stress”), associated by means of relations (the edges in Figure 7, such as “is a” or “regulates”). Each term is also matched with a set of relevant genes by means of a curated annotation, which is constantly updated according to the literature, in order to reflect the knowledge of the scientific community on the biological domain. Given a set of genes, it is possible to search for those terms that are over-represented (enriched) in the group of interest, compared to a background (a number of associated genes that is proportional to the node size in Figure 7), which is a larger set that potentially accounts for all the annotated transcriptional units. A hypergeometric test is usually performed for any possible term to statistically test the enrichments, and a threshold is then imposed on the corrected p-values or q-values, selecting the most significant results. These terms (labels in Figure 7) provide a broad overview of the biological processes involved with the selected genes. We perform these analyses using the Bioconductor R-package *clusterProfiler* (Yu <u>et al.</u>, 2012).

**Fig. 7.**
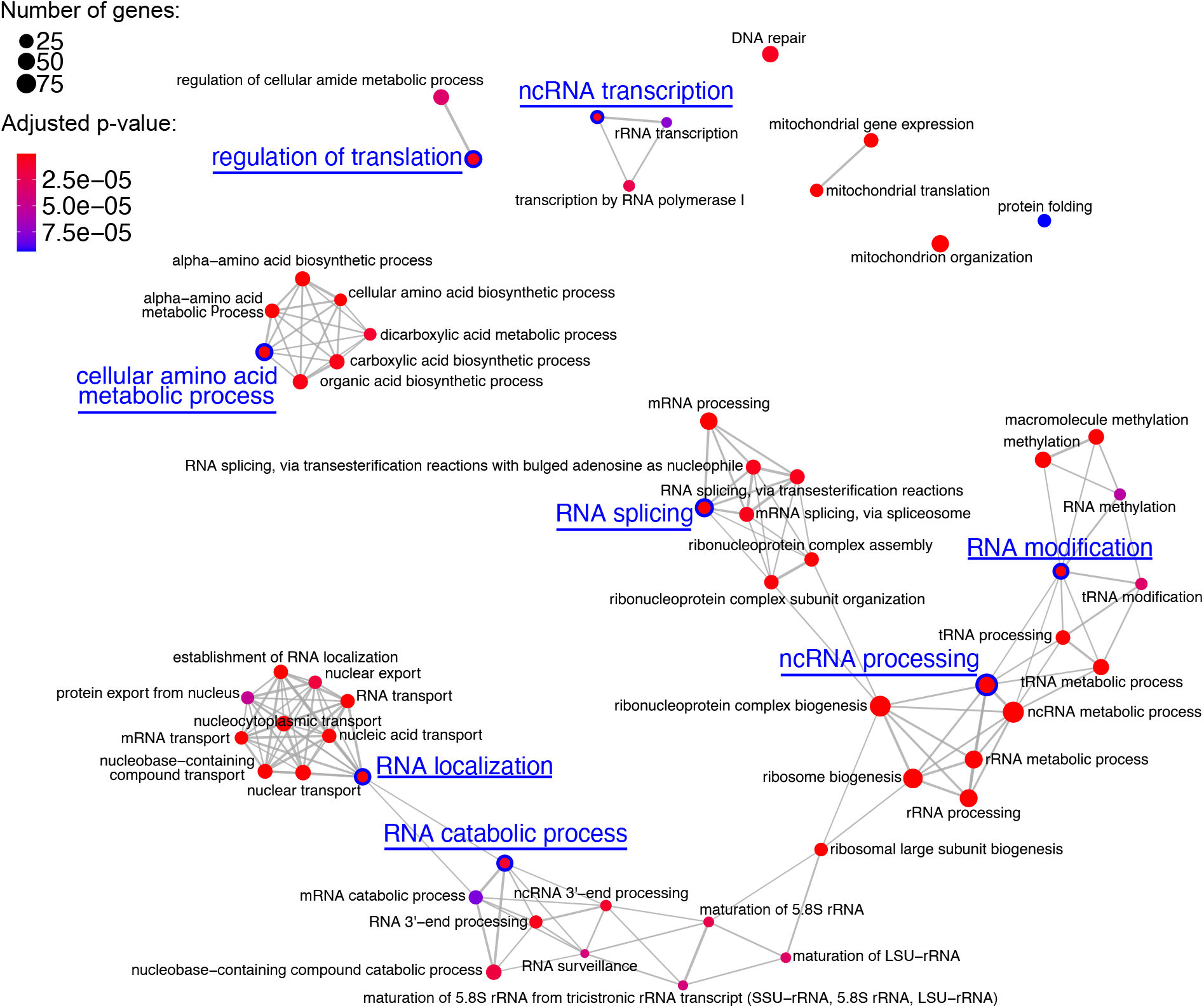
Synthesis rate Cluster 3 Gene Ontology analysis. Biological processes (nodes of the network) associated with the genes belonging to the third cluster identified from the analysis of the synthesis rate modulations. The network structure is indicative of the semantic similarity of the terms; i.e., linked and adjacent terms are close to each other in the reference ontology. The size of each dot is proportional to the number of genes identified in the cluster, while the color is a proxy of the significance of the enrichment that takes into account the total number of genes associated with the specific term in the background. The nodes that are easier to interpret and which are characteristic of different communities of the network are highlighted.

Our analyses, detailed in the next sections, return a picture of complex gene expression programs involving all three KRs. The clusters obtained for the rate of synthesis show a plethora of diverse up- and down-regulations, which differ according to their magnitude, timing and shape. Several clusters are supported by a significant enrichment, associated with basic biological processes as well as interesting terms in the context of cancer biology and RNA metabolism. As expected, the clustering derived from the temporal response of the synthesis rate only partially recapitulates the steady-state counterpart which, however, reveals that the sets of genes involved in the regulation of the RNA metabolism are also among the most rapidly transcribed. As expected, the signals emerging from our analyses are less defined for the post-transcriptional rates, since the regulation of RNA synthesis is predominant in the Myc induction context. Nevertheless, our analyses provide interesting insights into the biology of this system and show that large sets of genes are potentially involve indirectly in the cellular response.

#### Synthesis rate

The graphical description of the clusters associated with the synthesis rate is shown in Figure 5. *Cluster 1* accounts for 1293 genes characterized by peak-type functions (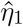 and 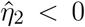), with a minimum between 1 and 3 hours and a small variance, which indicates quick responses. This behavior can be explained as a side effect of Myc activation, which causes the polarization of the resources required for the proficient transcription of the induced target genes (discussed in the following clusters). This response is globally compensated for in the last fraction of the time-course, thus indicating that these transcriptional units are required by the cell for its homoeostasis; this is confirmed by the enrichment analysis and pointed out in the literature (Dang, 2012; Chen <u>et al.</u>, 2018) (see Figure 7-SM). *Cluster 2* accounts for 1103 genes, characterized by a monotonic decrease of the synthesis rate (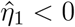 and 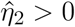). The temporal response is more heterogeneous, with 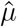 and 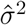 spanning large domains. This cluster contains genes that are involved in cell growth and development, cell adhesion, migration, and apoptotic signaling regulation (Figure 8-SM); these are interesting clues that point to the role played by Myc in cancer biology (Dang, 2012; Chen <u>et al.</u>, 2018). *Cluster 3*, which accounts for 904 genes, is composed of monotonic increasing responses (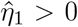 and 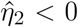) concentrated in the first 5 hours of the time-course. This behavior indicates a potential direct Myc regulation. These genes are involved in coding and non-coding RNA metabolisms (Figure 7). They affect *RNA localization*, e.g. exporting from the nucleus, *RNA splicing* and *non-coding RNA processing*, and also RNA stability, e.g. the *RNA catabolic process*. Moreover, these genes are related to *RNA modification*, an emerging dynamic regulatory layer of the transcript metabolism (Roundtree <u>et al.</u>, 2017). For example, *N*^6^ –*methyladenosine* is a methylated nucleotide (*methylation*, *RNA methylation*, *macromolecule methylation*) that is pervasive in the transcriptome of various species, e.g. mice, with a well established role in the control of transcript stability (Wang <u>et al.</u>, 2014) and translation (Wang <u>et al.</u>, 2015). Increasing evidence also links this RNA modification to synthesis a nd splicing ( see, for example, Louloupi <u>et al.</u>, 2018; Furlan <u>et al.</u>, 2019). The analysis of these terms provides a coherent picture that relates Myc activation to several regulatory layers of the RNA metabolism and translation (e.g., the *cellular amino acid metabolic process*, *regulation of translation*). This evidence supports the post-transcriptional rate modulations predicted by our approach and, partially, by INSPEcT. *Cluster 4* accounts for 392 monotonic + and − (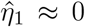 and 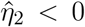 or 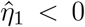 and 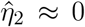, respectively) late responding genes (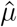 generally larger than 4 and up to 14). These transcriptional regulations are probably secondary responses and are clustered together due to their temporal features. *Clusters 5, 6 and 8* are composed of 299, 246 and 148 monotonically induced genes, respectively, in response to Myc activation. These clusters were split due to their temporal responses which are slower and more heterogeneous in *Cluster 6*, and less sharp in *Cluster 8* (up-c shape: 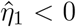 and 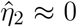). *Cluster 6* is partially related to non-coding RNA processing (see Figure 10-SM). *Cluster 7* is composed of 182 genes characterized by weak peak-like induction (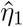 and 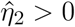), which occurs earlier in the time-course and quicker than those belonging to Cluster 1, as can be seen from the values of 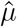 and 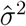. Finally, *Cluster 9* is composed of 142 genes characterized by a monotonic decrease of the synthesis rate (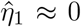 and 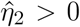). The response has a similar timing compared to *Cluster 2*, but is sharper and stronger; noticeably, these genes are involved in epithelial cell proliferation and migration, in agreement with the terms related to the second cluster. Our model also provides a clustering of genes according to their steady-state values of the synthesis rate (*β*_1_), specifically 6 clusters enumerated with a progression of letters. *Cluster A* is related to cell-cycle regulation, cell growth, apoptosis and RNA processing. The latter process is also enriched in *Clusters C and D* (Figures 20-SM, 22-SM and 23-SM), while, the genes in *Cluster F* are involved in epithelium cell migration and translation (Figure 24-SM). Interestingly, a remarkable percentage (≈ 30%) of the genes in *Clusters C and D* is also part of *Cluster 3* (Figure 35-SM). Moreover, these are clusters that are characterized by fast kinetics, which is a required condition, even though not sufficient, to quickly regulate the expression level of a gene, and is a clue of the fundamental regulatory role played by these transcriptional units (Figures 36-SM).

#### Processing rate

A graphical description of the clusters associated with the processing rate is given in Figure 6 (a). *Cluster 1* is composed of 3537 elements, which respond, to a great extent, within 3 hours from the stimulus with a moderate and sharp monotonic increase of *k*_2_. Because of the timing of the response and the extension of this cluster, this behavior could be due to the general feedback mechanisms which link RNA synthesis and processing. On the other hand, *Cluster 2* is composed of 1212 genes characterized by small and late, positive and negative monotonic responses. These are probably secondary responses that are mediated by the remarkable number of genes involved in the RNA processing regulation perturbed by Myc activation. The weakness of these modulations is puzzling but, since they take place in the most coarse-grained part of the time-course, our approach may only detect a reflex of the real biological regulation that could occur between the experimental observations. Finally, *Cluster 3* is composed of 148 genes characterized by a fast and transitory induction of the processing rate taking place in the first 2 hours of the time-course, and resulting in a transitory increase in the amount of mature RNAs. The Gene Ontology enrichment analysis of *Cluster 2* and *Cluster 3* do not provide any relevant terms, while several biological processes, already mentioned while discussing the synthesis rate response, can be found for *Cluster 1*. However, the enrichments are less significant and they disappear when the 4897 differentially expressed genes are used as the background instead of all the annotated ones (Figure 12-SM and Figure 13-SM). The same is true for all of the 5 clusters which our method returns for the processing of the steady-state rate (Figure 25-SM, Figure 26-SM and Figure 27-SM).

#### Degradation rate

The modulations of the degradation rate are divided into four clusters, see Figure 6 (b). *Cluster 1* is composed of 3094 elements characterized by very quick peak-type responses and with a minimum between 1 and 2 hours. As we have seen for *k*_2_, this behavior may be due to the coupling with the synthesis rate. *Cluster 2* and *Cluster 3* account for 1279 and 438 genes, respectively, characterized by late, positive and negative, monotonic responses, which are probably secondary. Like the processing rate, this could point to indirect regulations that were under-sampled due to the temporal design of the experiment. *Cluster 4* is composed of 86 genes characterized by a quick peak-like induction. A similar increase in the rate of degradation would result in a more rapid turnover of mature RNA molecules, while a similar behavior could be compatible with either deleterious transcripts for the cellular response, or those whose fast modulation is needed in the Myc induction context. The Gene Ontology enrichment analysis results are analogues with those discussed for the processing rate (Figure 14-SM and Figure 15-SM) for the 5 steady-state clusters (Figure 28-SM, Figure 29-SM and Figure 34-SM).

## 5. Final remarks

Motivated by a real data application, we here proposed a Bayesian approach to the analysis of RNA expression levels. In our proposal, the experimental data are hypothesized to be noisy observations of a true process, which is a solution to a system of ODEs. We assume that the ODEs depend on KRs, that are time- and gene-dependent. The KRs are the main object of inference, since they characterize the RNA life cycle and provide important insights into the analysis of gene expression levels. The temporal evolution of KRs is encoded with a new single family of functions, defined by only 5 parameters that can easily be interpreted from the biological perspective (i.e. initial value, relative log-fold-changes, and temporal location and duration of the response). The parameters are divided into two groups, according to their role in defining either the initial value of the KR or its temporal modulation. A mixture model, based on the DP, is defined for both of them, and for each KR. This allows us to find sets of genes with similar KR shapes or steady-state values to guide the inference. This approach is conceptually based on the well-established co-regulation of genes, which a cell often exploits to coordinate the expression level of the multiple transcripts required to operate a specific task. Therefore, the idea of including a clustering step in the inference process is not only biologically robust, but also provides a valuable information.

The results obtained with the new method are biologically relevant. The enrichment analysis of the clusters results in sets of terms that are meaningful in the Myc biology context, and which are in conceptual agreement with the shape of the responses. This is particularly true for the synthesis rate, which is the most informative regulatory layer in this system. Moreover, our method manages to identify a remarkable fraction of genes as post-transcriptionally regulated, thus pointing to weak indirect secondary responses.

A limitation of our method is the independence of the synthesis, processing and degradation rate clusters. In principle, this could be overcome by defining a mixture model on the parameters that shape the response of multiple rates. However, this inference would take place over a much larger space, which would be difficult to handle.

## Acknowledgements

The work of the first two authors has partially been developed under the MIUR grant Dipartimenti di Eccellenza 2018 – 2022 (E11G18000350001), conferred to the Dipartimento di Scienze Matematiche – DISMA, Politecnico di Torino. The computational resources were provided by the Dipartimento di Scienze Matematiche – DISMA, Politecnico di Torino. The authors would like to thank Mattia Pelizzola (CGS@SEMM – IIT), the Editor-in-Chief, the Associate Editor and the two anonymous reviewers for their comments that have greatly improved the manuscript.

