## Supplemental figures for "Multiple latent clustering model for the inference of RNA life-cycle kinetic rates from sequencing data"

### Thematic table of content


- **Supplemental Figure 1 (p.4) - Supplemental Figure 6 (p.9)):** Kinetic rates modulations in response to Myc activation, extended and magnified versions of Main Figures 5 and 6.
- **Supplemental Figure 7 (p.10) - Supplemental Figure 11 (p.14):** Gene Ontology enrichment analyses based on synthesis rate responses, and a universe compose off all the annotated transcriptional units.
- **Supplemental Figure 12 (p.15) - Supplemental Figure 13 (p.16):** Gene Ontology enrichment analyses based on processing rate responses, and a universe compose off all the annotated transcriptional units.
- **Supplemental Figure 14 (p.17) - Supplemental Figure 15 (p.18):** Gene Ontology enrichment analyses based on degradation rate responses, and a universe compose off all the annotated transcriptional units.
- **Supplemental Figure 16 (p.19) - Supplemental Figure 19 (p.22):** Gene Ontology enrichment analyses based on synthesis rate responses, and a universe compose off the genes involved in the analysis.
- **Supplemental Figure 20 (p.23) - Supplemental Figure 24 (p.27):** Gene Ontology enrichment analyses based on synthesis rate steady state values, and a universe compose off all the annotated transcriptional units.
- **Supplemental Figure 25 (p.28) - Supplemental Figure 27 (p.30):** Gene Ontology enrichment analyses based on processing rate steady state values, and a universe compose off all the annotated transcriptional units.
- **Supplemental Figure 28 (p.31) - Supplemental Figure 29 (p.32):** Gene Ontology enrichment analyses based on degradation rate steady state values, and a universe compose off all the annotated transcriptional units.
- **Supplemental Figure 30 (p.33) - Supplemental Figure 33 (p.36):** Gene Ontology enrichment analyses based on synthesis rate steady state values, and a universe compose off the genes involved in the analysis.
- **Supplemental Figure 34 (p.37):** Gene Ontology enrichment analyses based on degradation rate steady

state values, and a universe compose off the genes involved in the analysis.

- **Supplemental Figure 35 (p.38):**
Alluvial plots showing, for each rate, the flux of elements from the clusters defined on rates modulations to clusters defined on rates steady state values.

- **Supplemental Figure 36 (p.39):**
Distributions of the rates steady state values clustered according to the corresponding mixture models.

- **Supplemental Figure 37 (p.40):**
Correlation between the CRPS coefficients across RNA species.

- **Supplemental Figure 38 (p.41):**
Comparison between the models inferred through our proposal and INSPEcT.

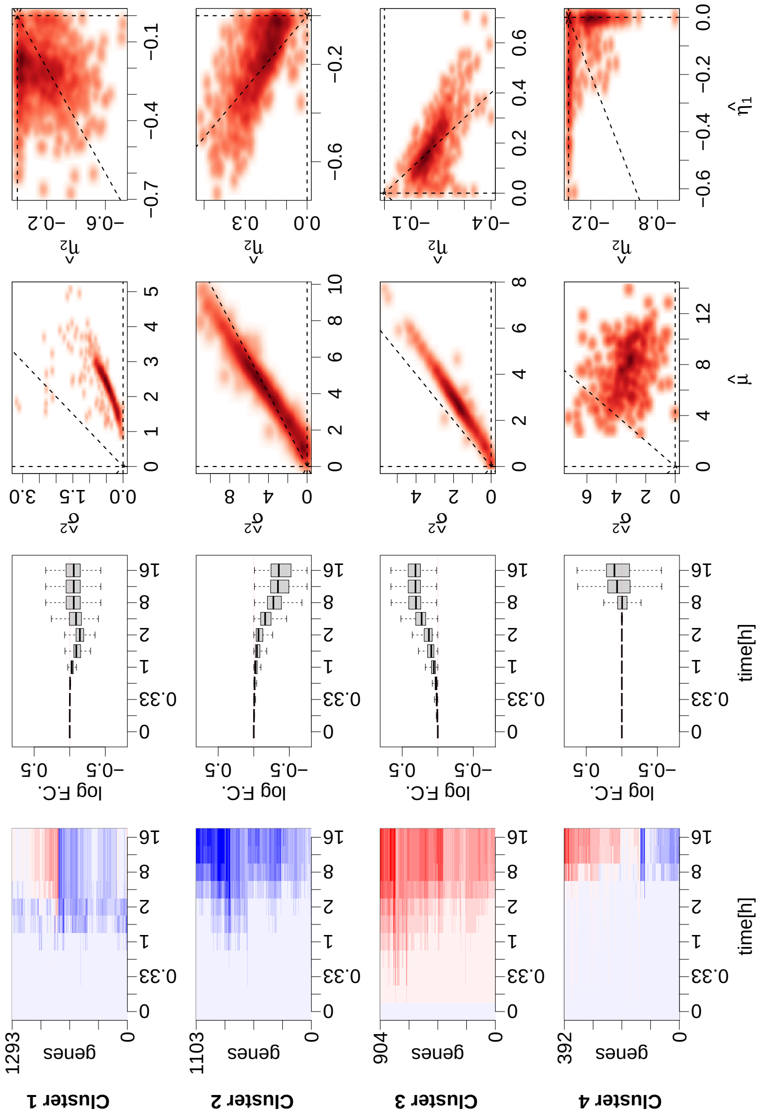


Supplemental Figure 1: **Synthesis rate modulations in response to Myc activation for Clusters 1-4.** (First column) Heatmaps showing the log-fold-changes of the rate $\hat{\eta}(t,0)$, compared to the first time-point, for each gene in the cluster (values saturated at $\pm 0.75$). (Second column) Boxplots showing the distribution of the synthesis rate log-fold-changes, compared to the first time-point (values saturated at $\pm 0.75$). (Third column) $\hat{\mu}$ versus $\hat{\sigma}^2$ smooth-scatter plot. (Fourth column) $\hat{\eta}_1$ versus $\hat{\eta}_2$ smooth-scatter plot. The dashed lines represent the horizontal and vertical axes and the bisector of the first and third quadrants.

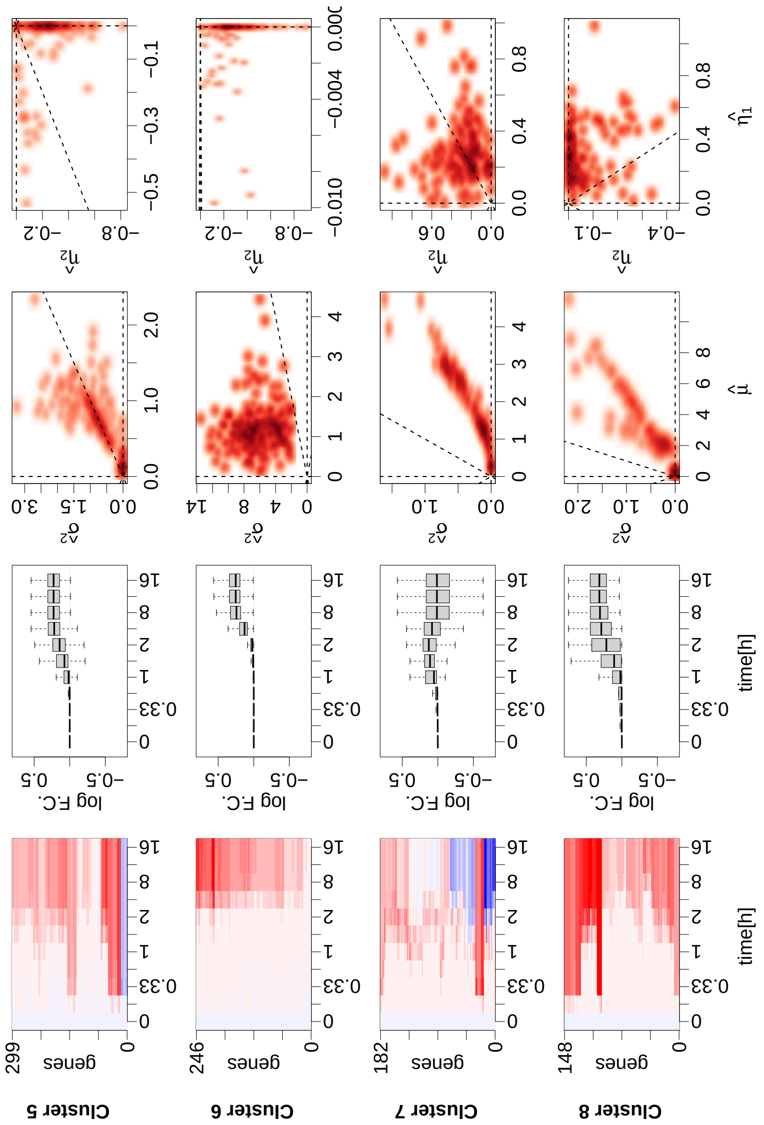


Supplemental Figure 2: **Synthesis rate modulations in response to Myc activation for Clusters 5-8.** (First column) Heatmaps showing the log-fold-changes of the rate $\hat{\eta}(t, 0)$, compared to the first time-point, for each gene in the cluster (values saturated at $\pm 0.75$). (Second column) Boxplots showing the distribution of the synthesis rate log-fold-changes, compared to the first time-point (values saturated at $\pm 0.75$). (Third column) $\hat{\mu}$ versus $\hat{\sigma}^2$ smooth-scatter plot. (Fourth column) $\hat{\eta}_1$ versus $\hat{\eta}_2$ smooth-scatter plot. The dashed lines represent the horizontal and vertical axes and the bisector of the first and third quadrants.

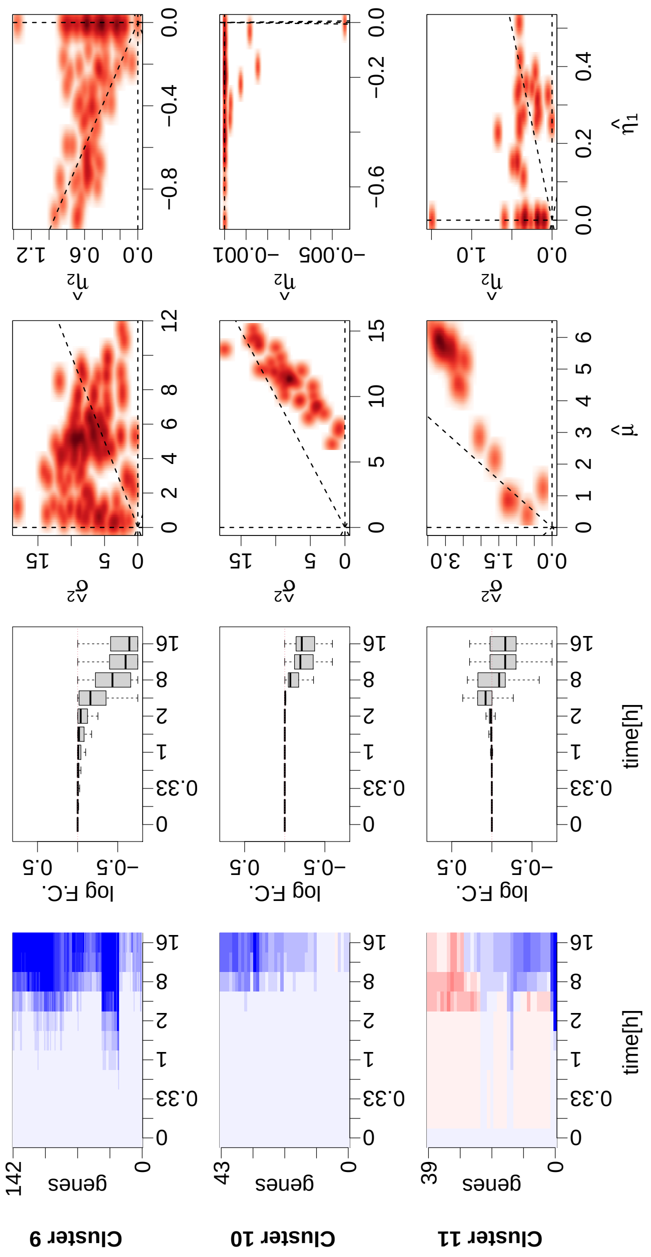


Supplemental Figure 3: **Synthesis rate modulations in response to Myc activation for Clusters 9-11.** (First column) Heatmaps showing the log-fold-changes of the rate $\hat{\eta}(t, 0)$, compared to the first time-point, for each gene in the cluster (values saturated at $\pm 0.75$). (Second column) Boxplots showing the distribution of the synthesis rate log-fold-changes, compared to the first time-point (values saturated at $\pm 0.75$). (Third column) $\hat{\mu}$ versus $\hat{\sigma}^2$ smooth-scatter plot. (Fourth column) $\hat{\eta}_1$ versus $\hat{\eta}_2$ smooth-scatter plot. The dashed lines represent the horizontal and vertical axes and the bisector of the first and third quadrants.

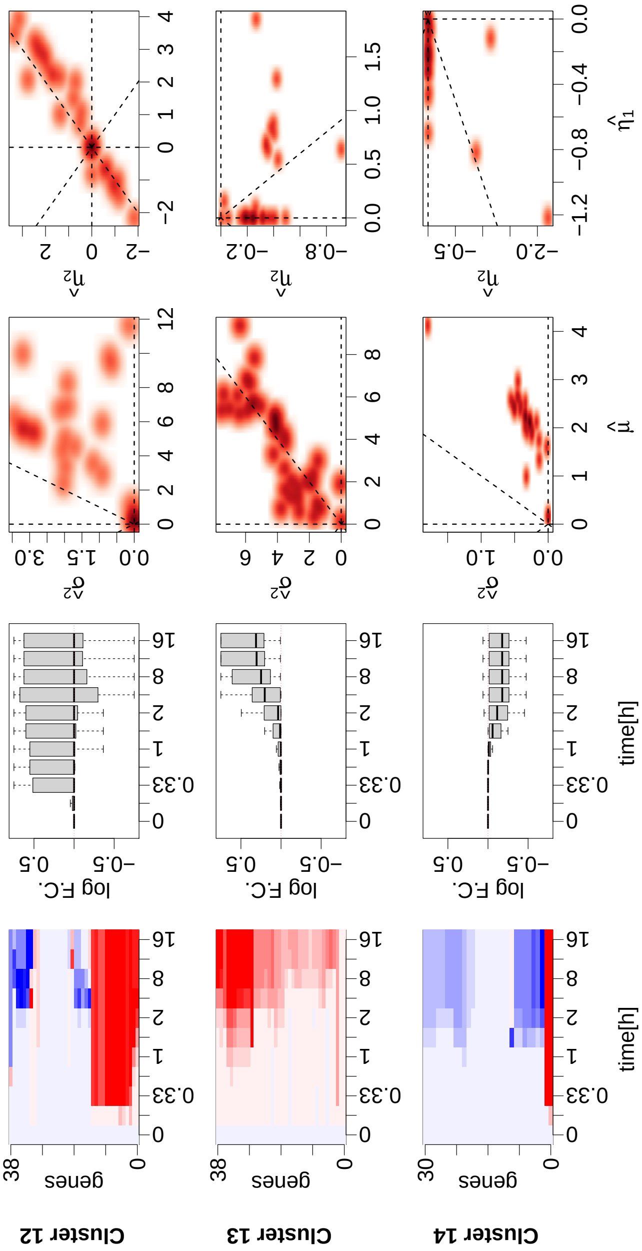


Supplemental Figure 4: **Synthesis rate modulations in response to Myc activation for Clusters 12-14.** (First column) Heatmaps showing the log-fold-changes of the rate $\hat{\eta}(t, 0)$, compared to the first time-point, for each gene in the cluster (values saturated at $\pm 0.75$). (Second column) Box-plots showing the distribution of the synthesis rate log-fold-changes, compared to the first time-point (values saturated at $\pm 0.75$). (Third column) $\hat{\mu}$ versus $\hat{\sigma}^2$ smooth-scatter plot. (Fourth column) $\hat{\eta}_1$ versus $\hat{\eta}_2$ smooth-scatter plot. The dashed lines represent the horizontal and vertical axes and the bisector of the first and third quadrants.

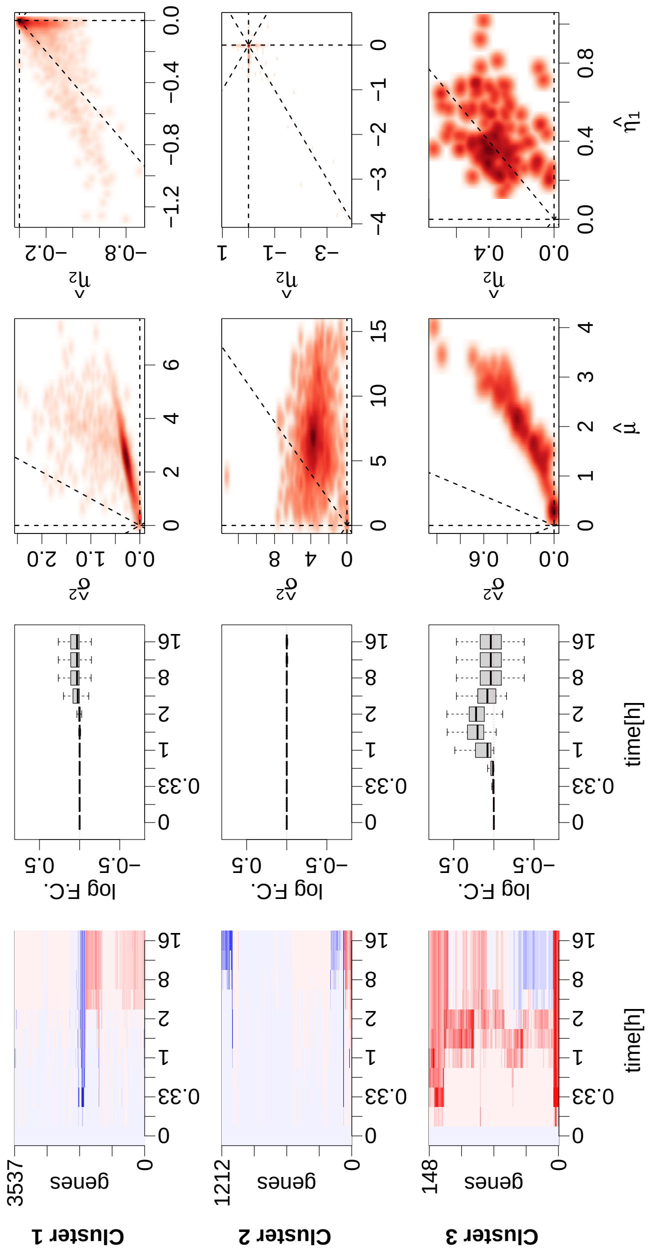


Supplemental Figure 5: **Processing rate modulations in response to Myc activation.** (First column) Heatmaps showing the log-fold-changes of the rate $\hat{\eta}(t,0)$, compared to the first time-point, for each gene in the cluster (values saturated at $\pm 0.75$). (Second column) Boxplots showing the distribution of the processing rate log-fold-changes, compared to the first time-point (values saturated at $\pm 0.75$). (Third column) $\hat{\mu}$ versus $\hat{\sigma}^2$ smooth-scatter plot. (Fourth column) $\hat{\eta}_1$ versus $\hat{\eta}_2$ smooth-scatter plot. The dashed lines represent the horizontal and vertical axes and the bisector of the first and third quadrants.

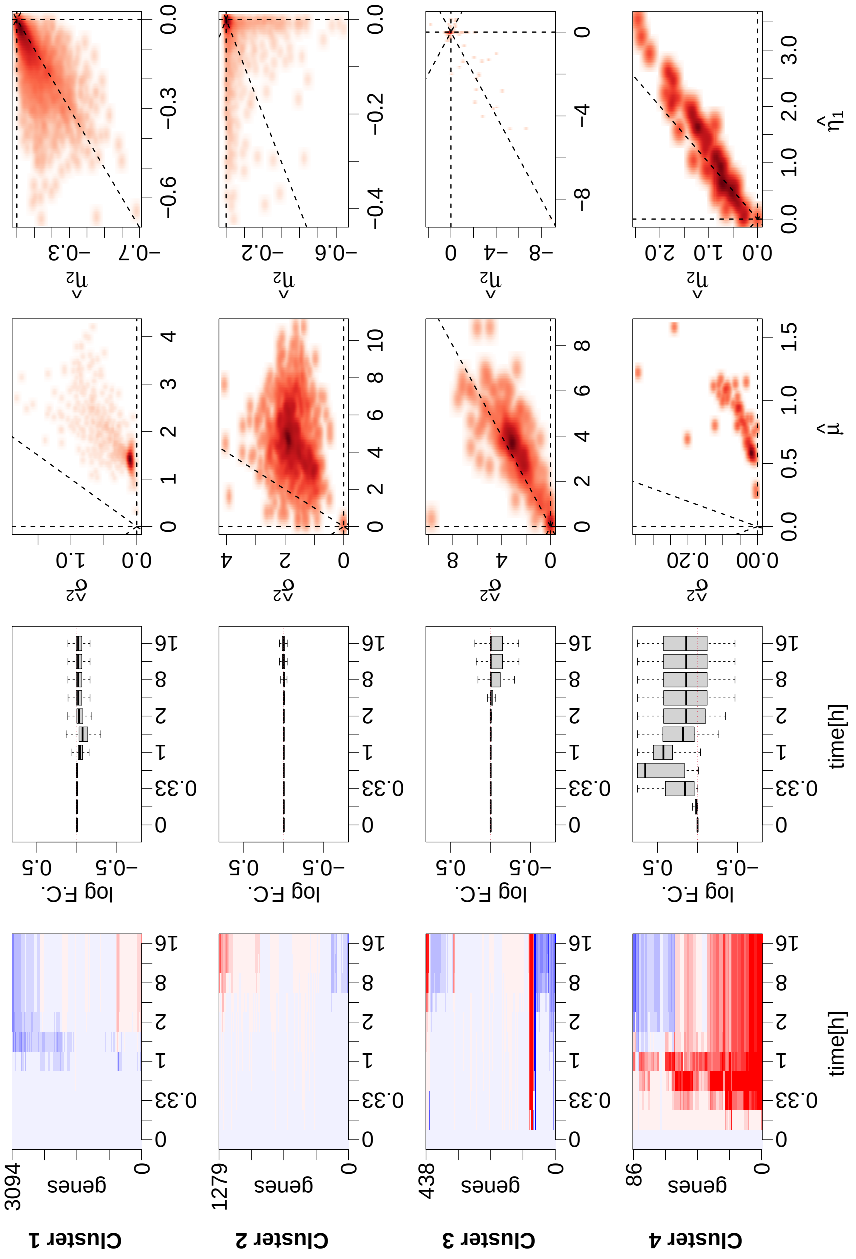


Supplemental Figure 6: **Degradation rate modulations in response to Myc activation for Clusters.** (First column) Heatmaps showing the log-fold-changes of the rate $\hat{\eta}(t, 0)$, compared to the first time-point, for each gene in the cluster (values saturated at $\pm 0.75$). (Second column) Boxplots showing the distribution of the degradation rate log-fold-changes, compared to the first time-point (values saturated at $\pm 0.75$). (Third column) $\hat{\mu}$ versus $\hat{\sigma}^2$ smooth-scatter plot. (Fourth column) $\hat{\eta}_1$ versus $\hat{\eta}_2$ smooth-scatter plot. The dashed lines represent the horizontal and vertical axes and the bisector of the first and third quadrants.

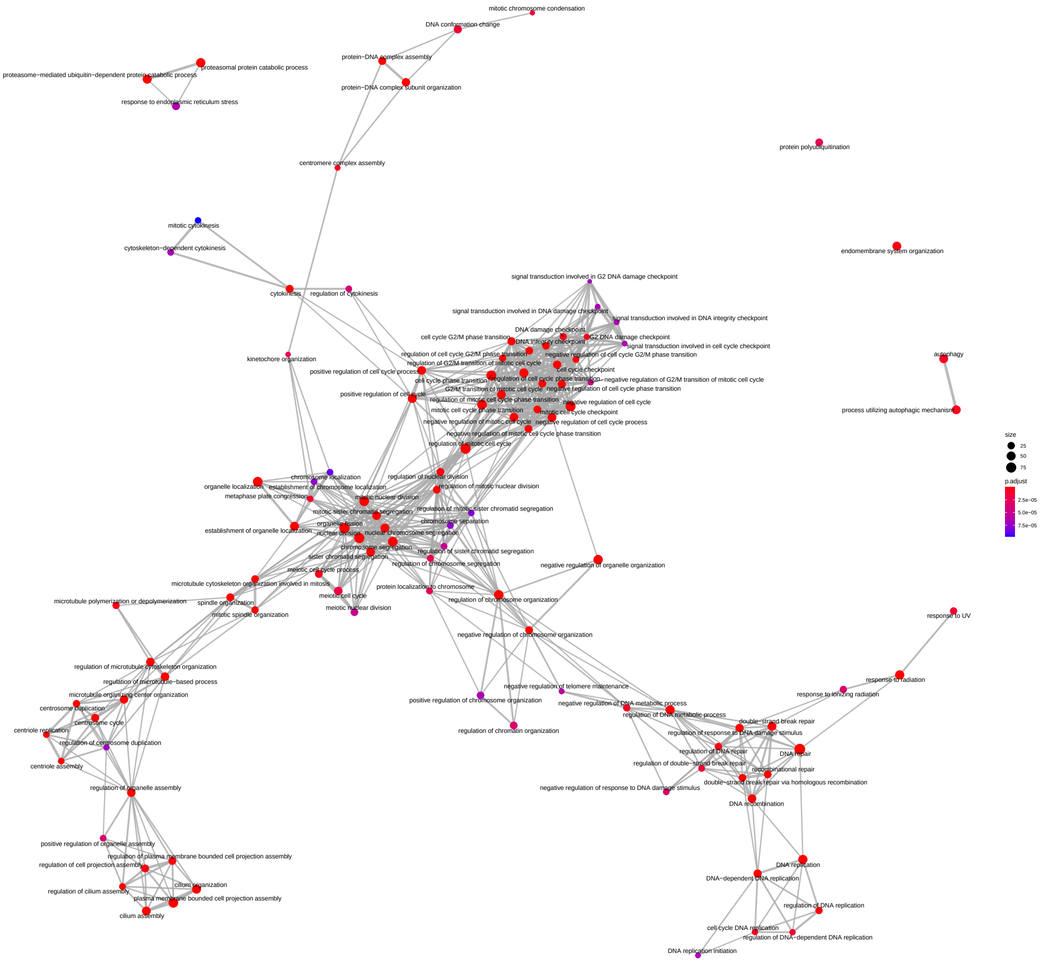


Supplemental Figure 7: **Synthesis rate modulation of the Cluster 1 Gene Ontology analysis.** Biological processes (nodes of the network) associated with the genes belonging to the first cluster identified from the analysis of the synthesis rate modulations. The network structure is indicative of the semantic similarity of the terms; i.e. linked and adjacent terms are close to each other in the reference ontology. The size of each dot is proportional to the number of genes identified in the cluster, while the color is a proxy of the significance of the enrichment which takes into account the total number of genes associated with the specific term.

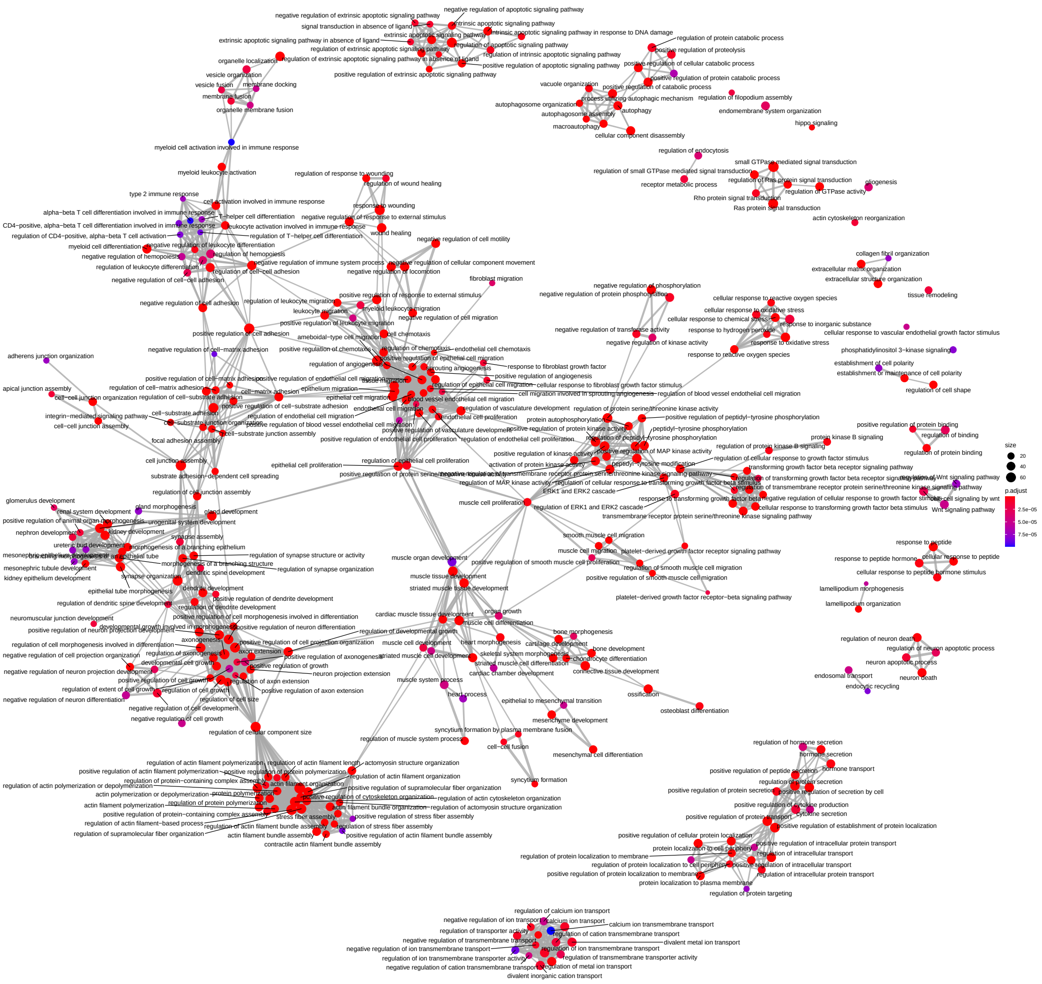

Supplemental Figure 8: **Synthesis rate modulation of the Cluster 2 Gene Ontology analysis.** Biological processes (nodes of the network) associated with the genes belonging to the second cluster identified from the analysis of the synthesis rate modulations. The network structure is indicative of the semantic similarity of the terms; i.e. linked and adjacent terms are close to each other in the reference ontology. The size of each dot is proportional to the number of genes identified in the cluster, while the color is a proxy of the significance of the enrichment which takes into account the total number of genes associated with the specific term.

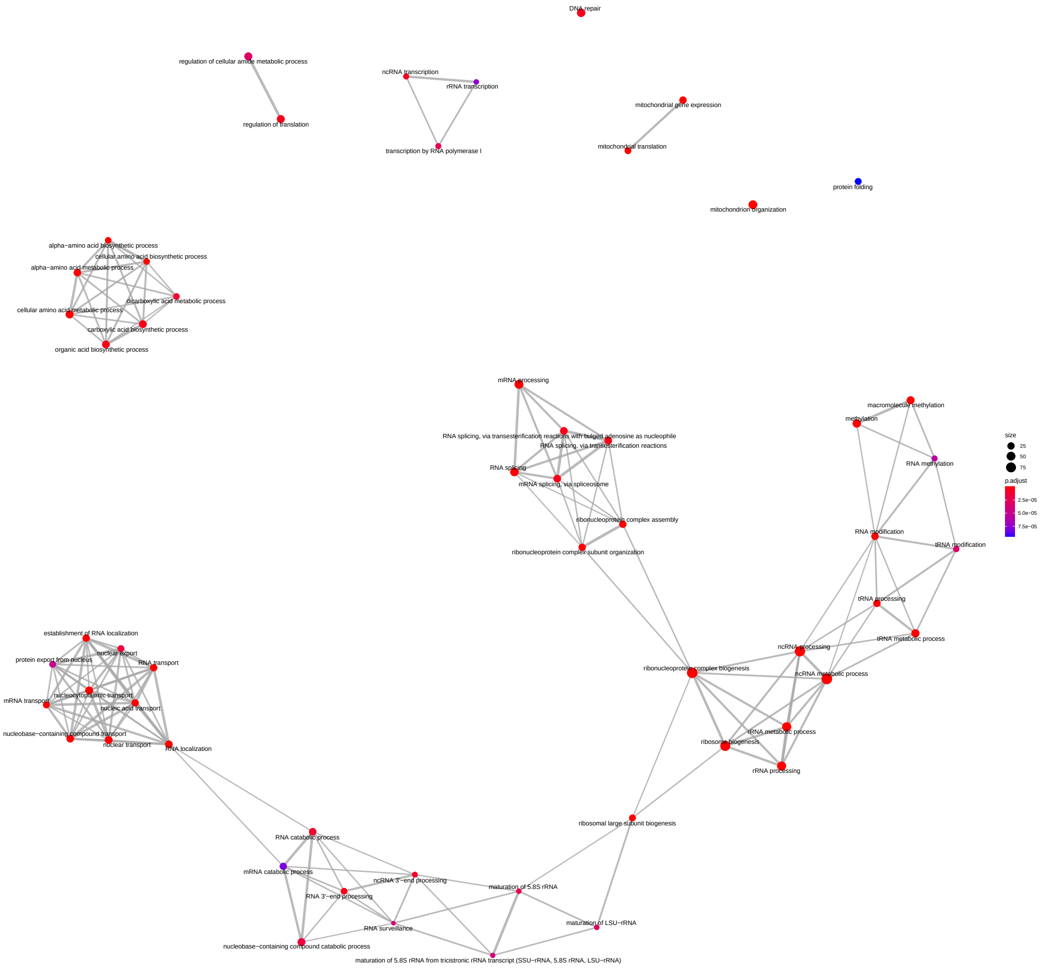


Supplemental Figure 9: **Synthesis rate modulation of the Cluster 3 Gene Ontology analysis.** Biological processes (nodes of the network) associated with the genes belonging to the third cluster identified from the analysis of the synthesis rate modulations. The network structure is indicative of the semantic similarity of the terms; i.e. linked and adjacent terms are close to each other in the reference ontology. The size of each dot is proportional to the number of genes identified in the cluster, while the color is a proxy of the significance of the enrichment which takes into account the total number of genes associated with the specific term.

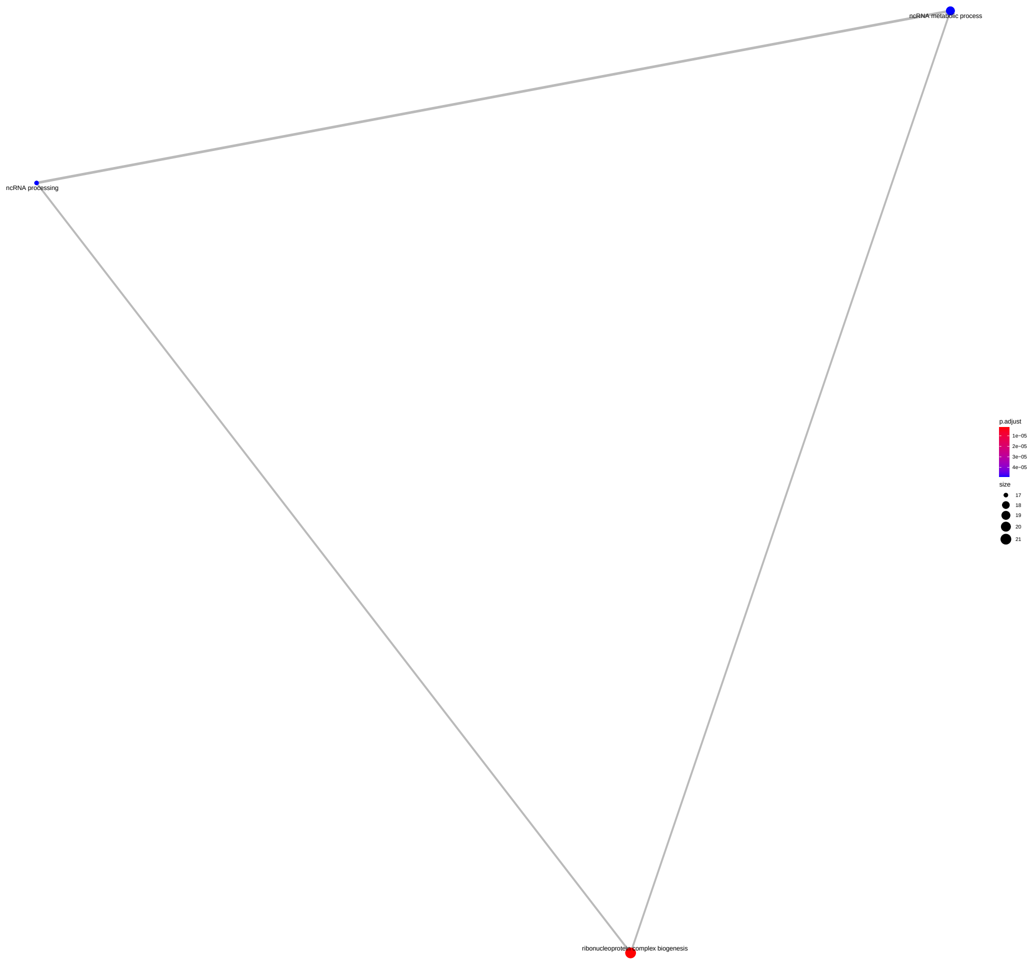


Supplemental Figure 10: **Synthesis rate modulation of the Cluster 6 Gene Ontology analysis.** Biological processes (nodes of the network) associated with the genes belonging to the sixth cluster identified from the analysis of the synthesis rate modulations. The network structure is indicative of the semantic similarity of the terms; i.e. linked and adjacent terms are close to each other in the reference ontology. The size of each dot is proportional to the number of genes identified in the cluster, while the color is a proxy of the significance of the enrichment which takes into account the total number of genes associated with the specific term.

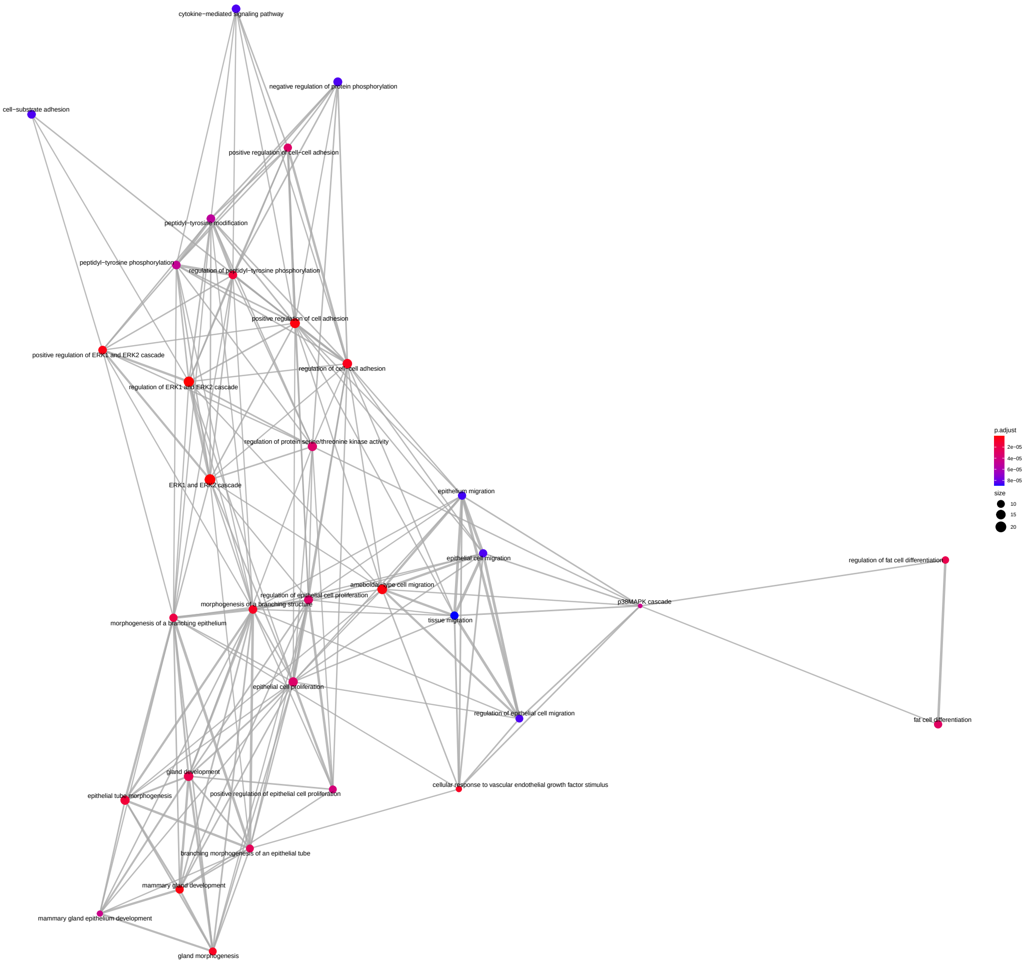


Supplemental Figure 11: **Synthesis rate modulation of the Cluster 9 Gene Ontology analysis.** Biological processes (nodes of the network) associated with the genes belonging to the ninth cluster identified from the analysis of the synthesis rate modulations. The network structure is indicative of the semantic similarity of the terms; i.e. linked and adjacent terms are close to each other in the reference ontology. The size of each dot is proportional to the number of genes identified in the cluster, while the color is a proxy of the significance of the enrichment which takes into account the total number of genes associated with the specific term.

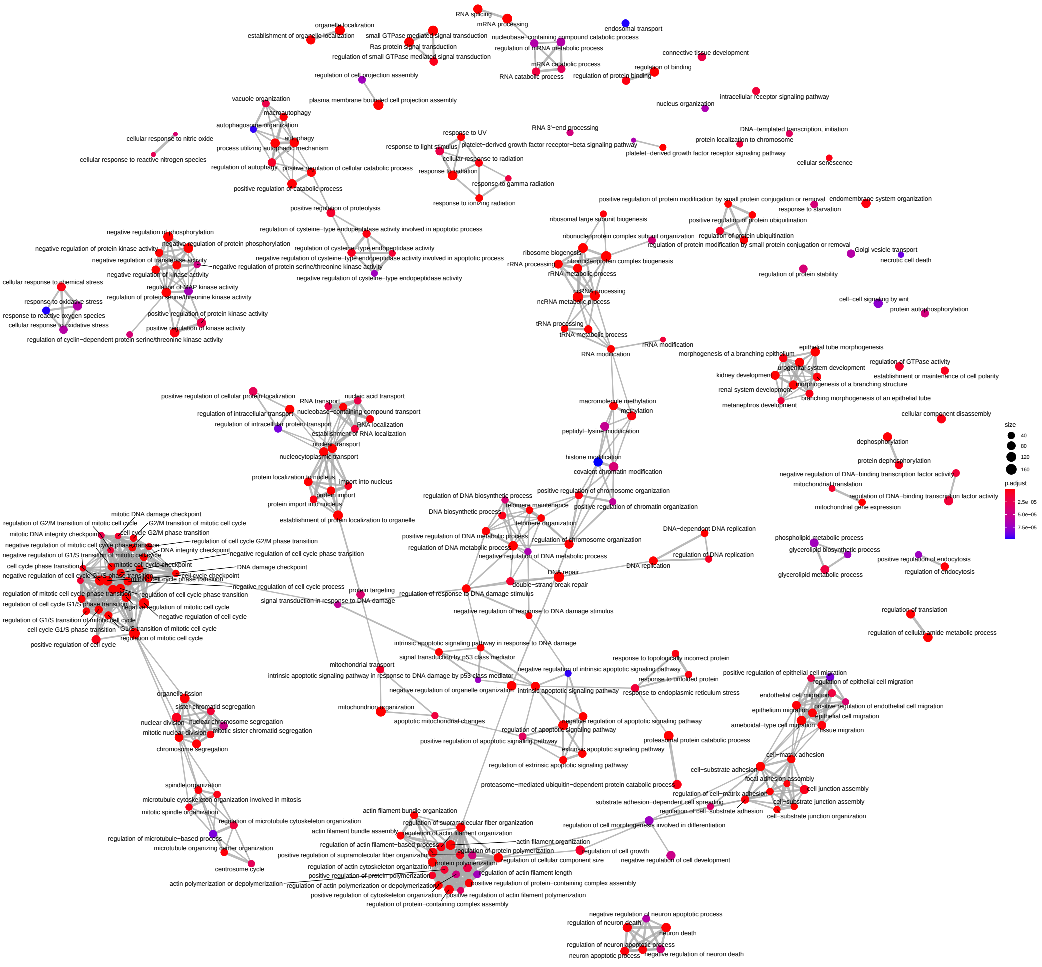


Supplemental Figure 12: **Processing rate modulation of the Cluster 1 Gene Ontology analysis.** Biological processes (nodes of the network) associated with the genes belonging to the first cluster identified from the analysis of the processing rate modulations. The network structure is indicative of the semantic similarity of the terms; i.e. linked and adjacent terms are close to each other in the reference ontology. The size of each dot is proportional to the number of genes identified in the cluster, while the color is a proxy of the significance of the enrichment which takes into account the total number of genes associated with the specific term.

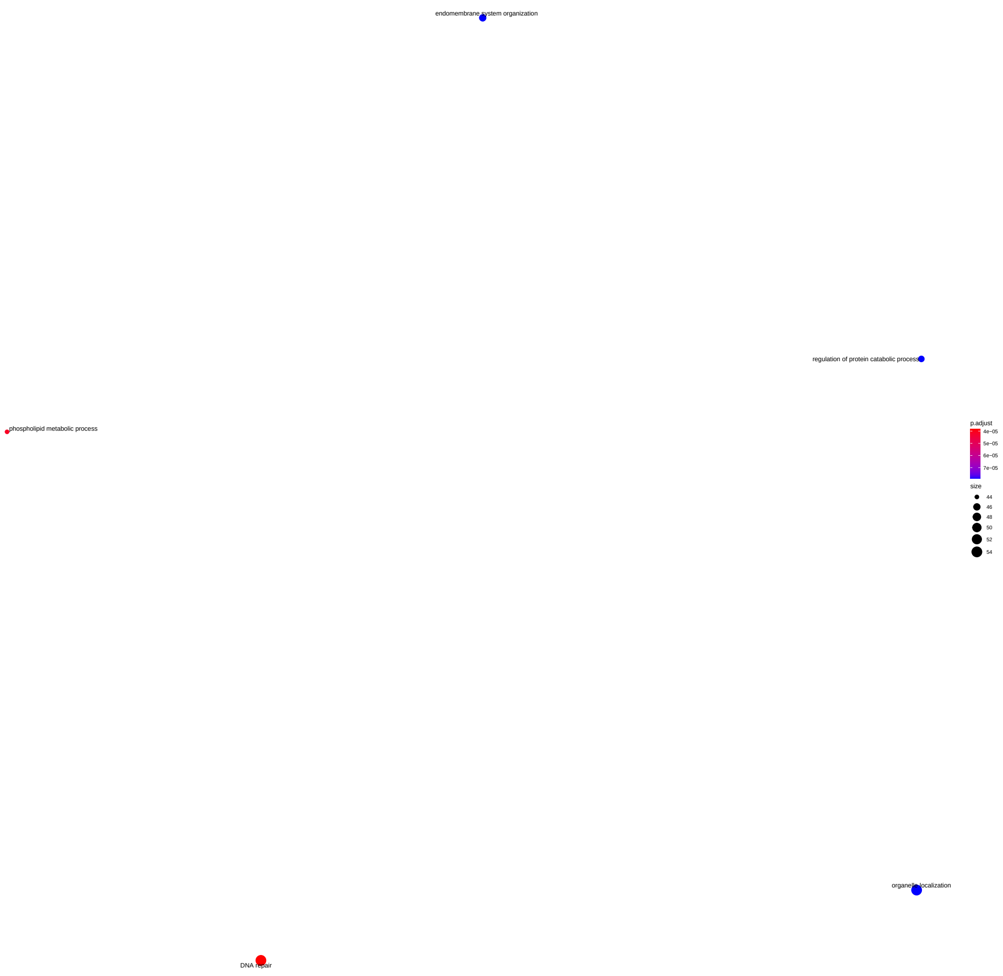


Supplemental Figure 13: **Processing rate modulation of the Cluster 2 Gene Ontology analysis.** Biological processes (nodes of the network) associated with the genes belonging to the second cluster identified from the analysis of the processing rate modulations. The network structure is indicative of the semantic similarity of the terms; i.e. linked and adjacent terms are close to each other in the reference ontology. The size of each dot is proportional to the number of genes identified in the cluster, while the color is a proxy of the significance of the enrichment which takes into account the total number of genes associated with the specific term.

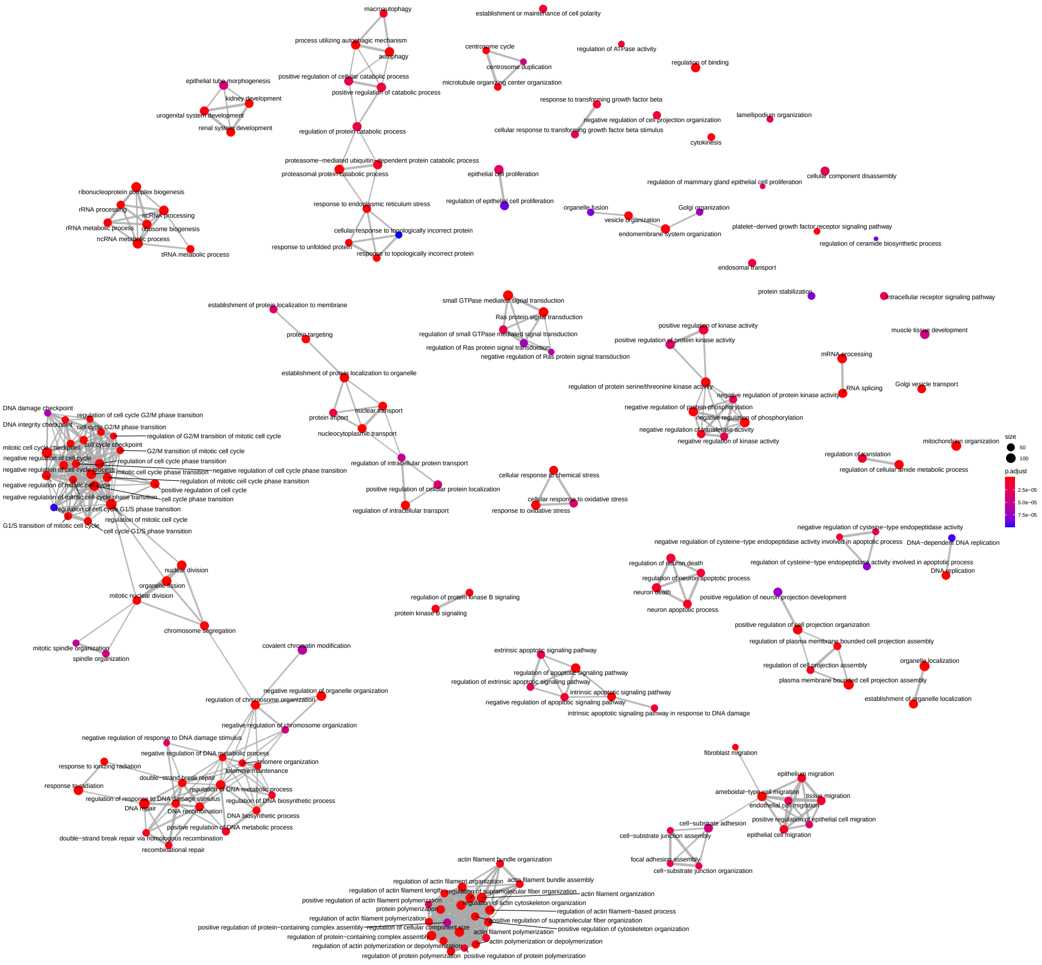


Supplemental Figure 14: **Degradation rate modulation of the Cluster 1 Gene Ontology analysis.** Biological processes (nodes of the network) associated with the genes belonging to the first cluster identified from the analysis of the degradation rate modulations. The network structure is indicative of the semantic similarity of the terms; i.e. linked and adjacent terms are close to each other in the reference ontology. The size of each dot is proportional to the number of genes identified in the cluster, while the color is a proxy of the significance of the enrichment which takes into account the total number of genes associated with the specific term.

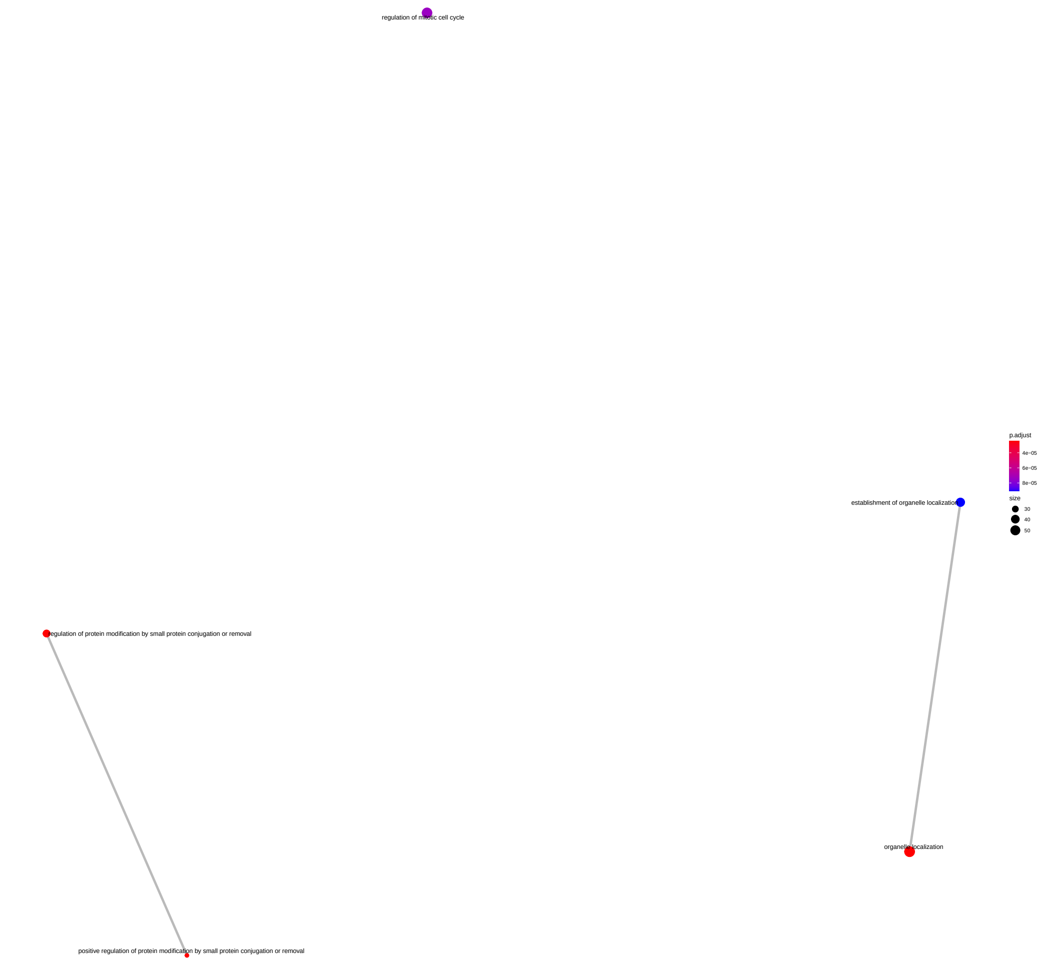


Supplemental Figure 15: **Degradation rate modulation of the Cluster 2 Gene Ontology analysis.** Biological processes (nodes of the network) associated with the genes belonging to the second cluster identified from the analysis of the degradation rate modulations. The network structure is indicative of the semantic similarity of the terms; i.e. linked and adjacent terms are close to each other in the reference ontology. The size of each dot is proportional to the number of genes identified in the cluster, while the color is a proxy of the significance of the enrichment which takes into account the total number of genes associated with the specific term.

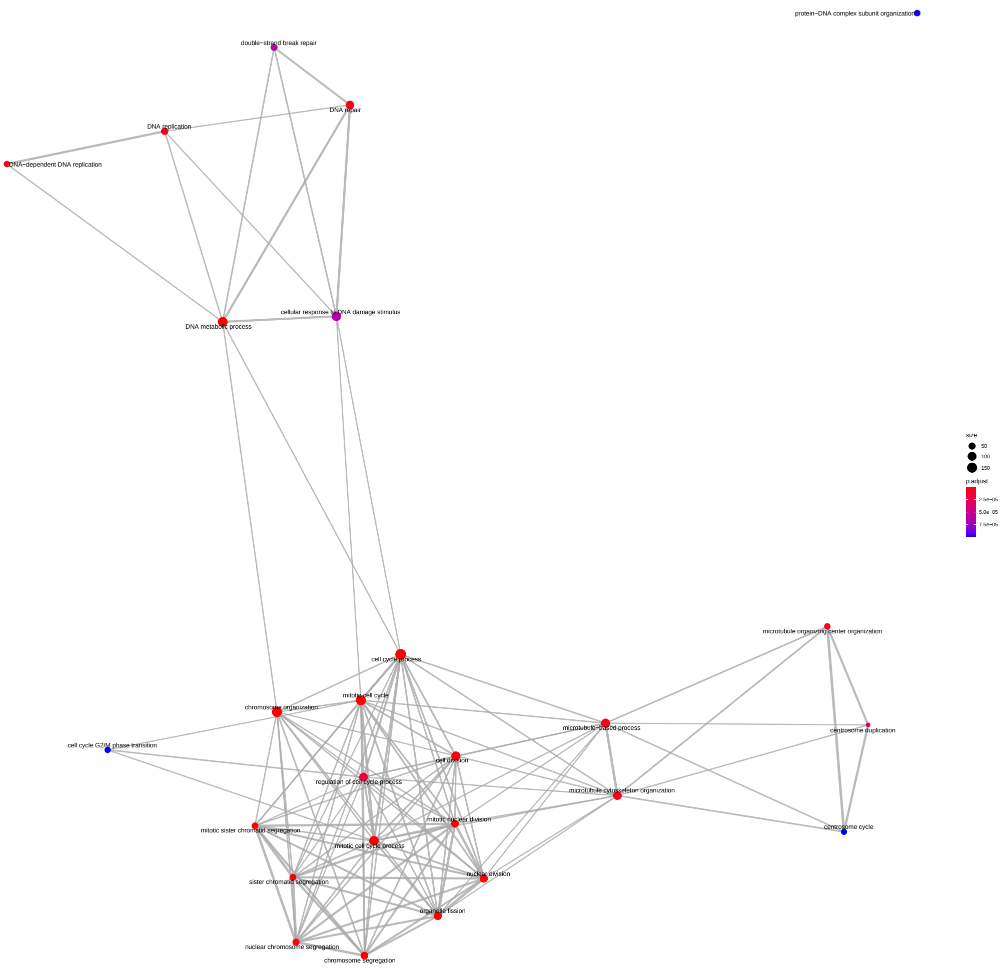


Supplemental Figure 16: **Synthesis rate modulation of the Cluster 1 Gene Ontology analysis on small universe.** Biological processes (nodes of the network) associated with the genes belonging to the first cluster identified from the analysis of the synthesis rate modulations. The network structure is indicative of the semantic similarity of the terms; i.e. linked and adjacent terms are close to each other in the reference ontology. The size of each dot is proportional to the number of genes identified in the cluster, while the color is a proxy of the significance of the enrichment which takes into account the total number of genes associated with the specific term among the 4897 used for the analysis.

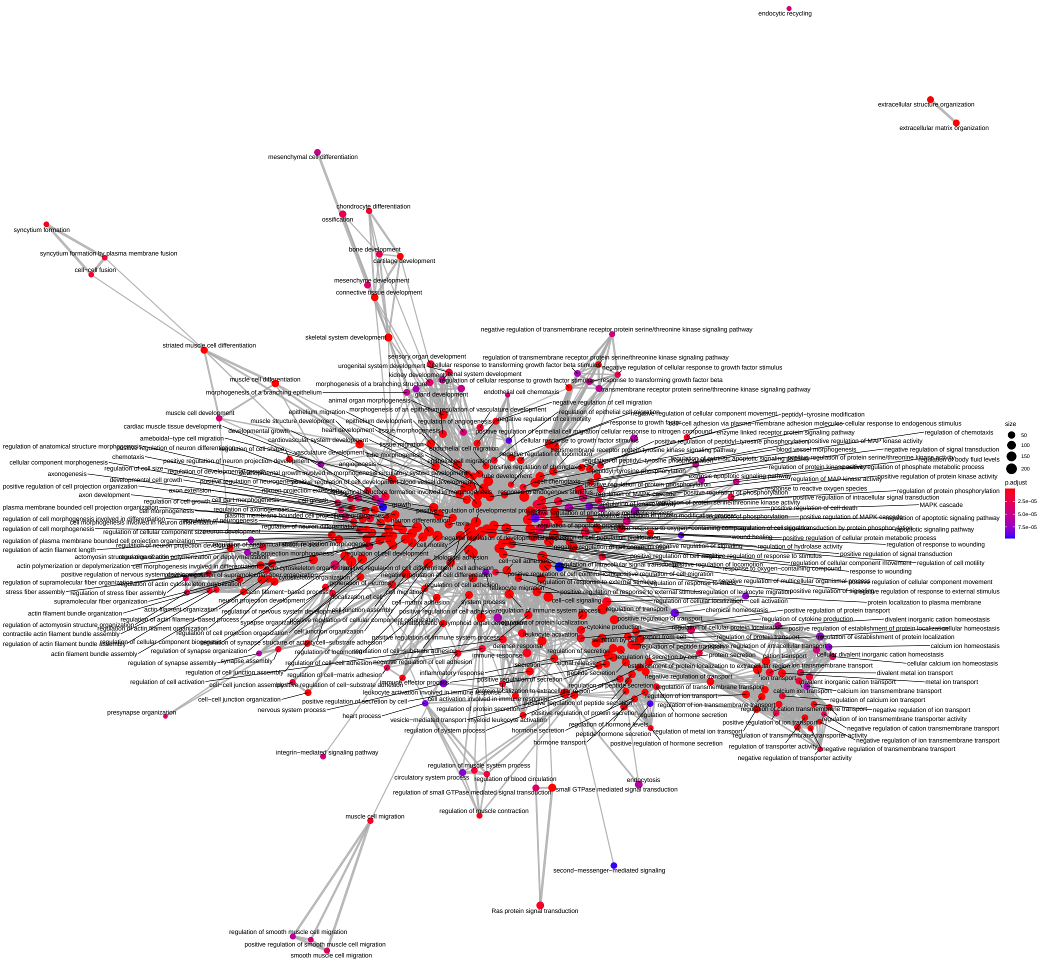


Supplemental Figure 17: **Synthesis rate modulation of the Cluster 2 Gene Ontology analysis on small universe.** Biological processes (nodes of the network) associated with the genes belonging to the second cluster identified from the analysis of the synthesis rate modulations. The network structure is indicative of the semantic similarity of the terms; i.e. linked and adjacent terms are close to each other in the reference ontology. The size of each dot is proportional to the number of genes identified in the cluster, while the color is a proxy of the significance of the enrichment which takes into account the total number of genes associated with the specific term among the 4897 used for the analysis.

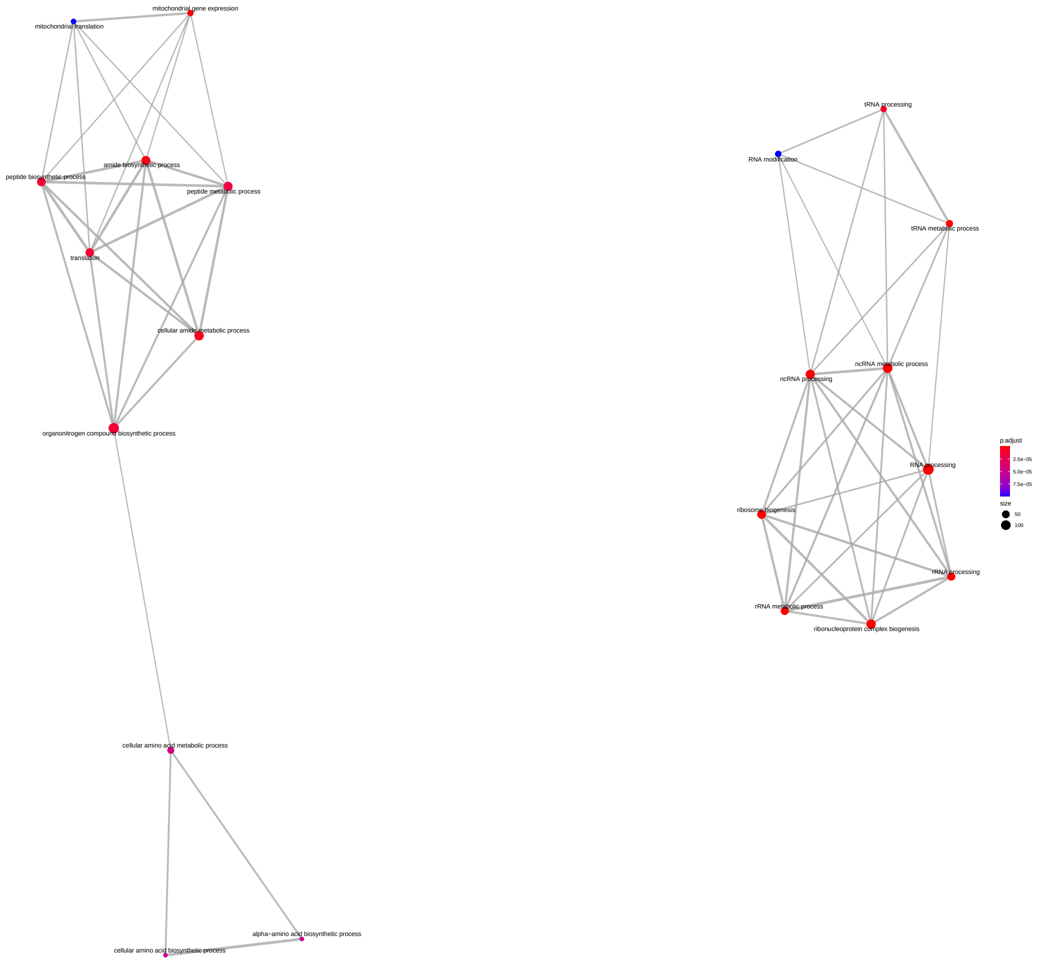


Supplemental Figure 18: **Synthesis rate modulation of the Cluster 3 Gene Ontology analysis on small universe.** Biological processes (nodes of the network) associated with the genes belonging to the third cluster identified from the analysis of the synthesis rate modulations. The network structure is indicative of the semantic similarity of the terms; i.e. linked and adjacent terms are close to each other in the reference ontology. The size of each dot is proportional to the number of genes identified in the cluster, while the color is a proxy of the significance of the enrichment which takes into account the total number of genes associated with the specific term among the 4897 used for the analysis.

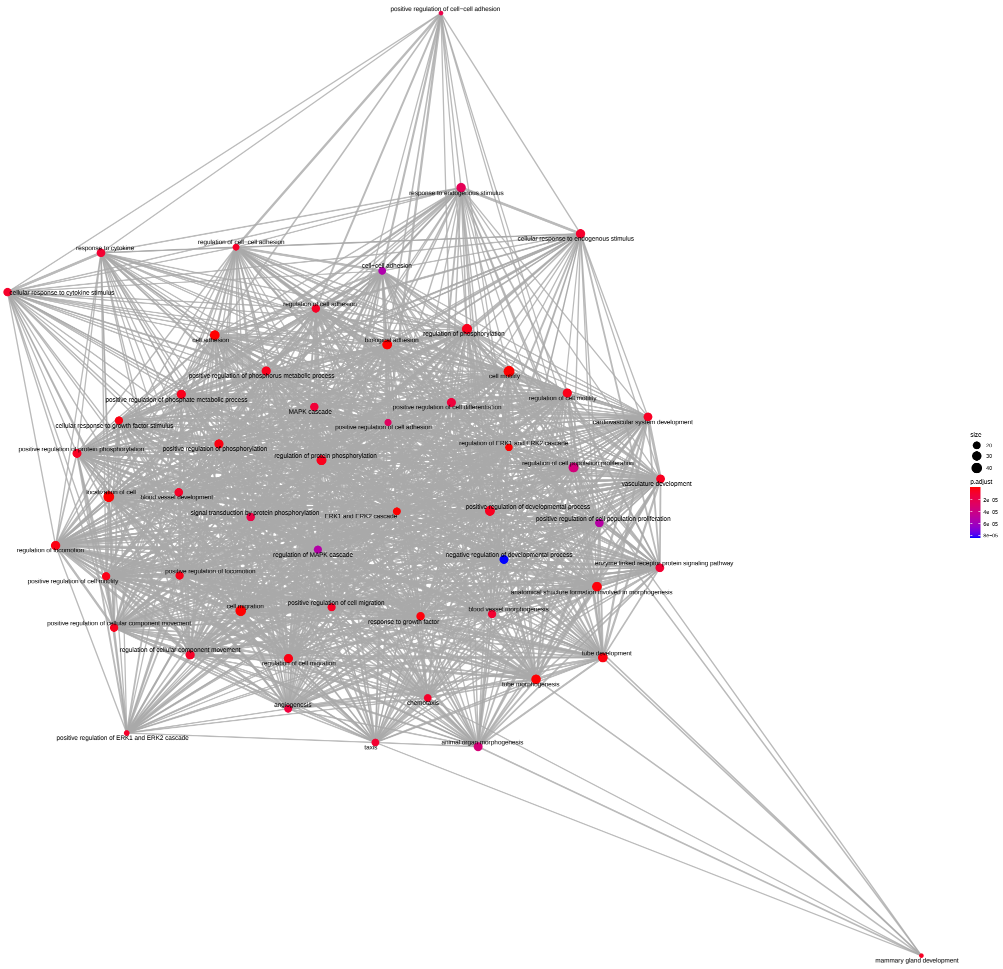


Supplemental Figure 19: **Synthesis rate modulation of the Cluster 9 Gene Ontology analysis on small universe.** Biological processes (nodes of the network) associated with the genes belonging to the ninth cluster identified from the analysis of the synthesis rate modulations. The network structure is indicative of the semantic similarity of the terms; i.e. linked and adjacent terms are close to each other in the reference ontology. The size of each dot is proportional to the number of genes identified in the cluster, while the color is a proxy of the significance of the enrichment which takes into account the total number of genes associated with the specific term among the 4897 used for the analysis.

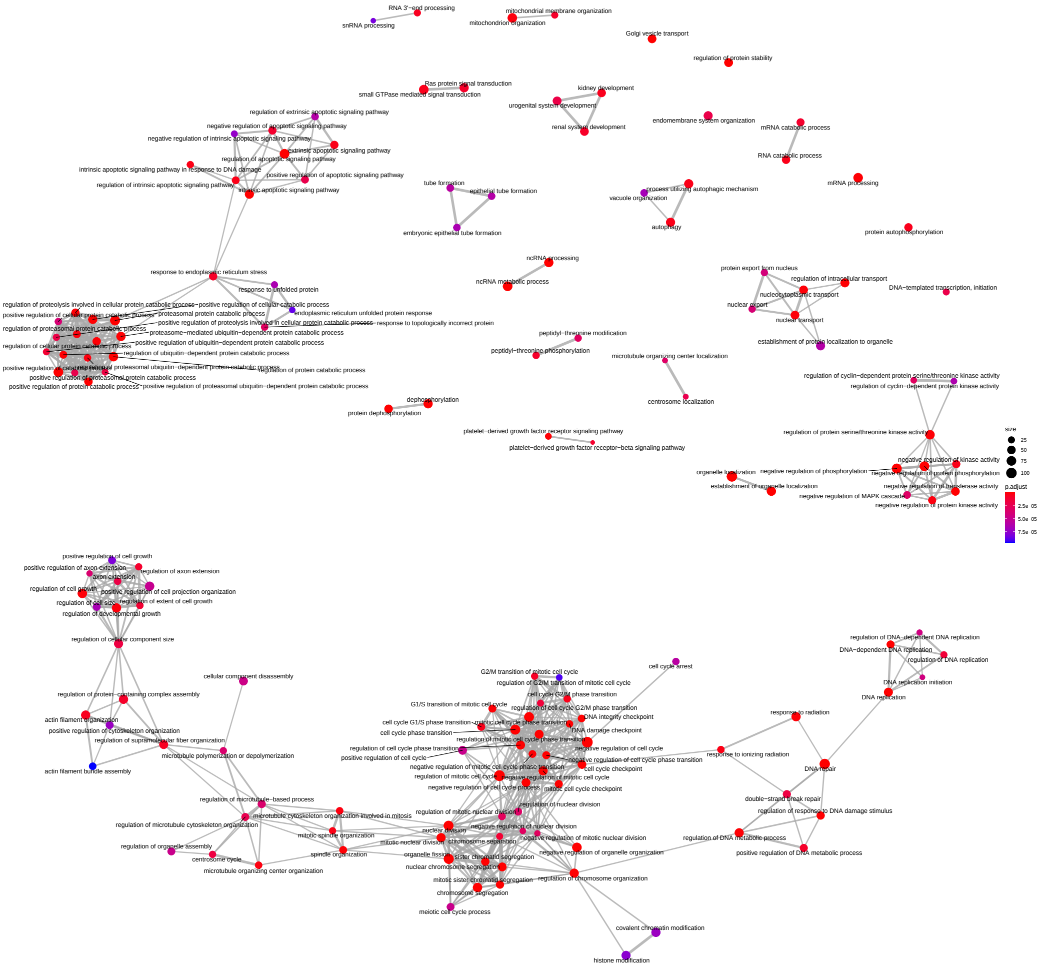


Supplemental Figure 20: **Synthesis rate steady state of the Cluster A Gene Ontology analysis.** Biological processes (nodes of the network) associated with the genes belonging to the first cluster identified from the analysis of the synthesis rate steady state values. The network structure is indicative of the semantic similarity of the terms; i.e. linked and adjacent terms are close to each other in the reference ontology. The size of each dot is proportional to the number of genes identified in the cluster, while the color is a proxy of the significance of the enrichment which takes into account the total number of genes associated with the specific term.

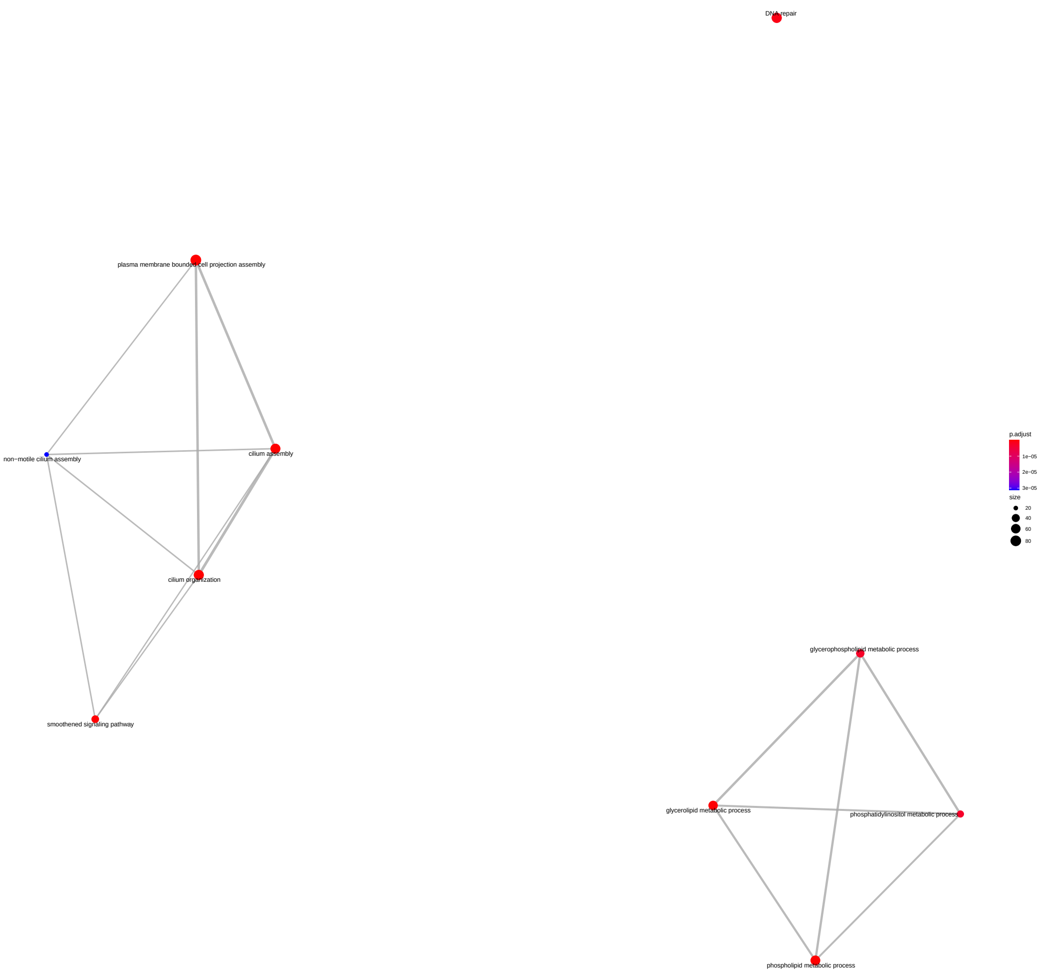


Supplemental Figure 21: **Synthesis rate steady state of the Cluster B Gene Ontology analysis.** Biological processes (nodes of the network) associated with the genes belonging to the second cluster identified from the analysis of the synthesis rate steady state values. The network structure is indicative of the semantic similarity of the terms; i.e. linked and adjacent terms are close to each other in the reference ontology. The size of each dot is proportional to the number of genes identified in the cluster, while the color is a proxy of the significance of the enrichment which takes into account the total number of genes associated with the specific term.

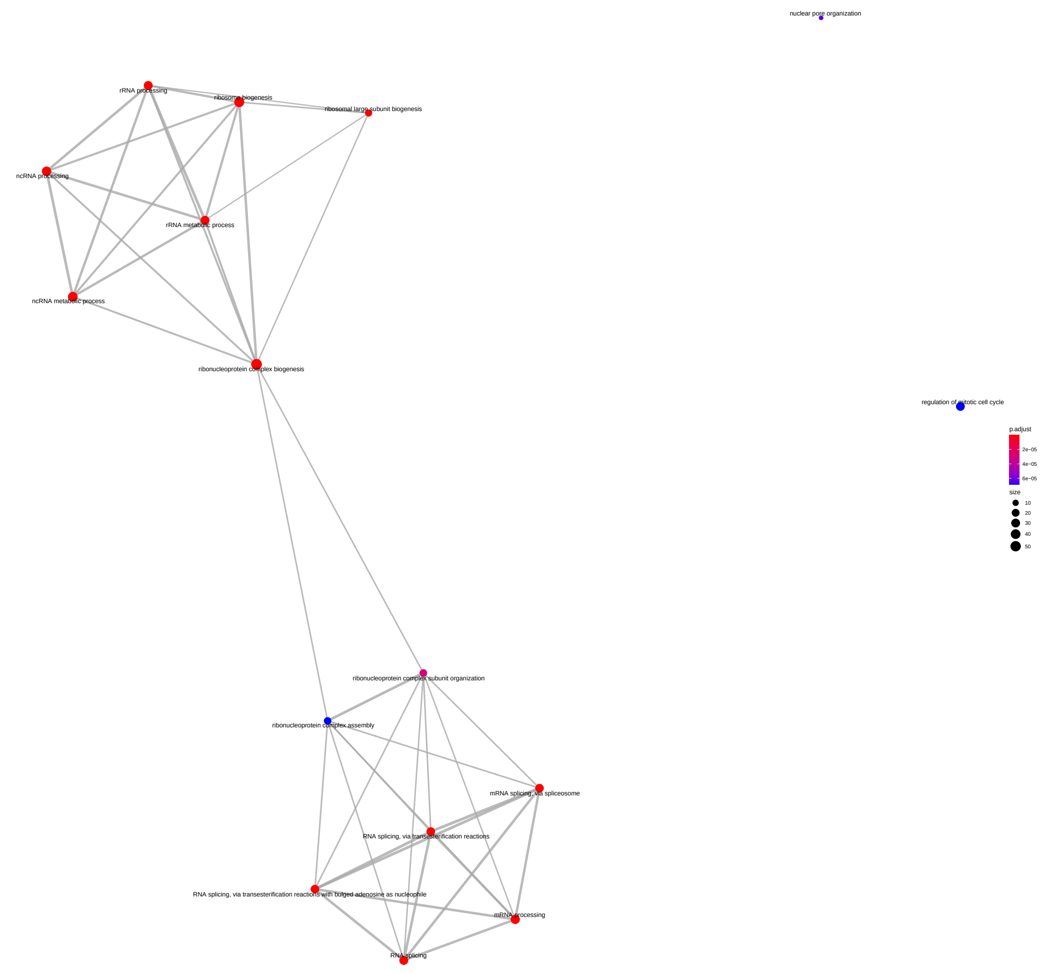


Supplemental Figure 22: **Synthesis rate steady state of the Cluster C Gene Ontology analysis.** Biological processes (nodes of the network) associated with the genes belonging to the third cluster identified from the analysis of the synthesis rate steady state values. The network structure is indicative of the semantic similarity of the terms; i.e. linked and adjacent terms are close to each other in the reference ontology. The size of each dot is proportional to the number of genes identified in the cluster, while the color is a proxy of the significance of the enrichment which takes into account the total number of genes associated with the specific term.

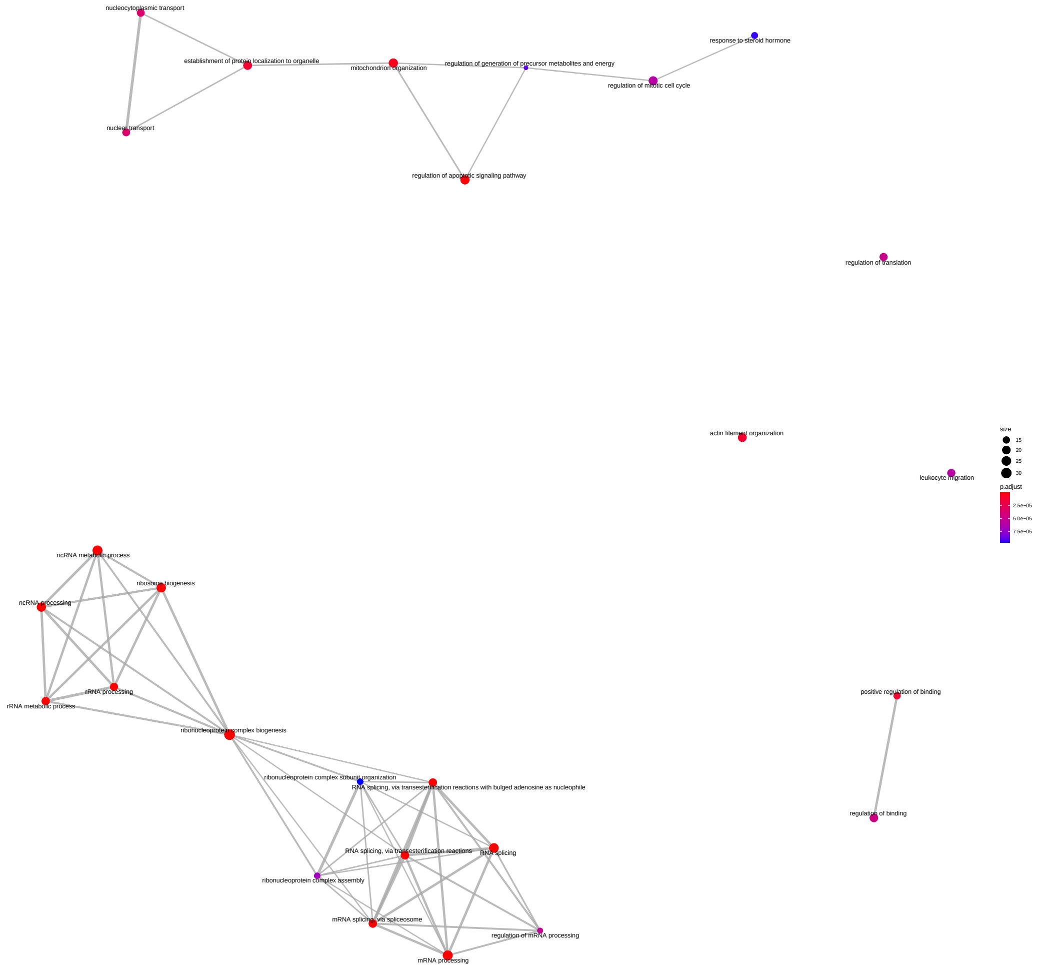


Supplemental Figure 23: **Synthesis rate steady state of the Cluster D Gene Ontology analysis.** Biological processes (nodes of the network) associated with the genes belonging to the fourth cluster identified from the analysis of the synthesis rate steady state values. The network structure is indicative of the semantic similarity of the terms; i.e. linked and adjacent terms are close to each other in the reference ontology. The size of each dot is proportional to the number of genes identified in the cluster, while the color is a proxy of the significance of the enrichment which takes into account the total number of genes associated with the specific term.

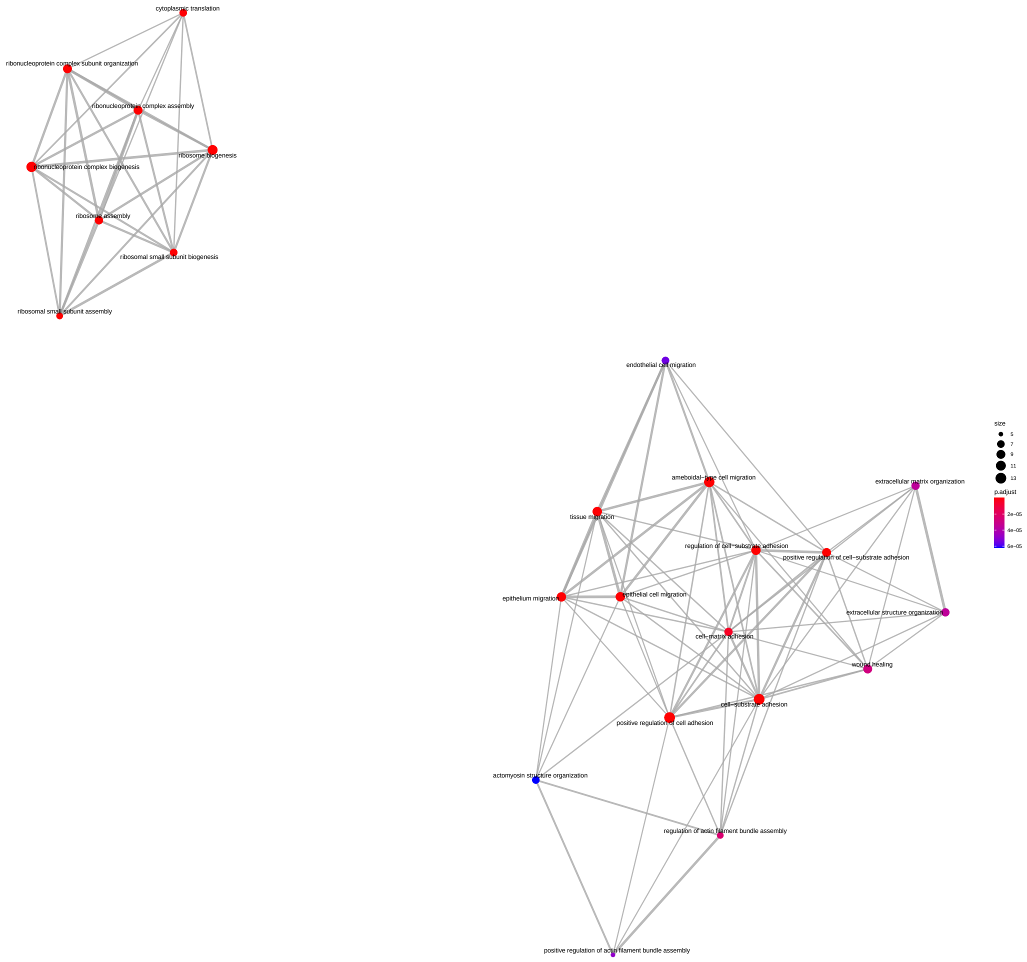


Supplemental Figure 24: **Synthesis rate steady state of the Cluster F Gene Ontology analysis.** Biological processes (nodes of the network) associated with the genes belonging to the sixth cluster identified from the analysis of the synthesis rate steady state values. The network structure is indicative of the semantic similarity of the terms; i.e. linked and adjacent terms are close to each other in the reference ontology. The size of each dot is proportional to the number of genes identified in the cluster, while the color is a proxy of the significance of the enrichment which takes into account the total number of genes associated with the specific term.

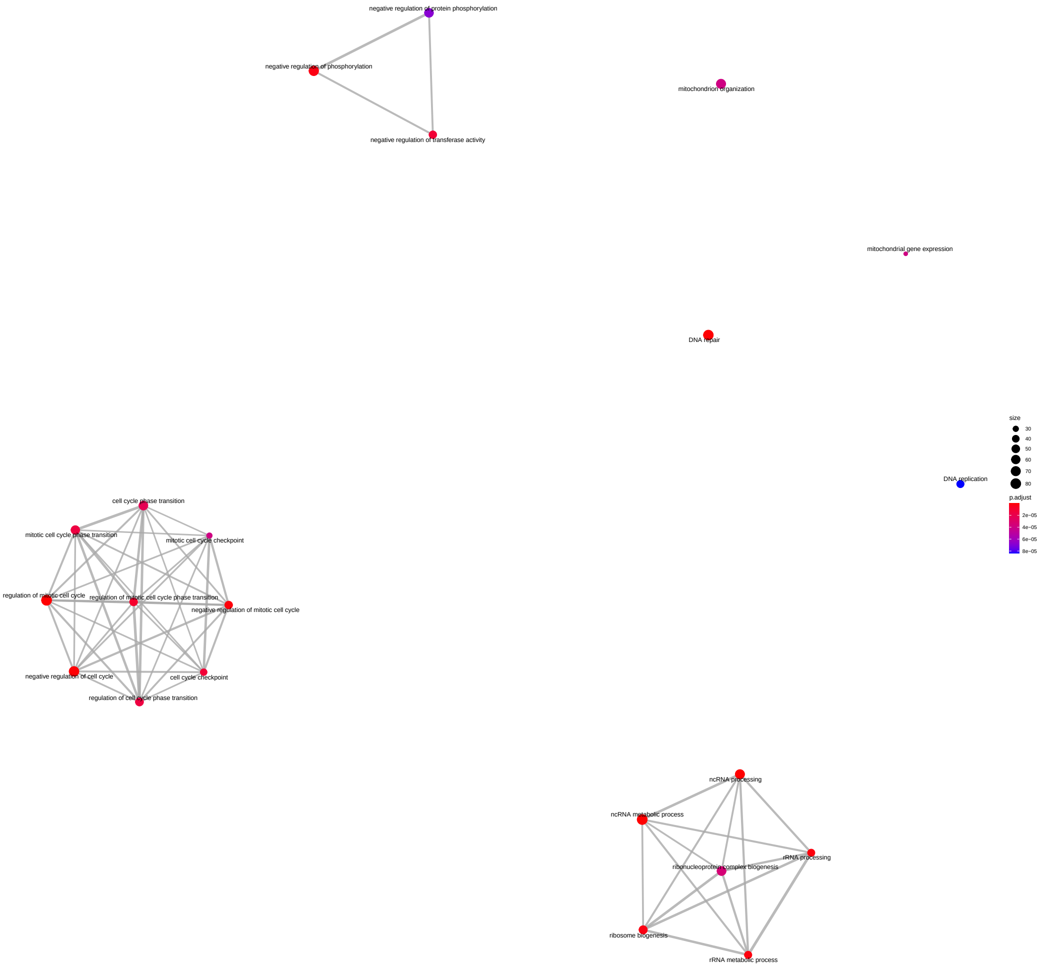


Supplemental Figure 25: **Processing steady-state rate of the Cluster A Gene Ontology analysis.** Biological processes (nodes of the network) associated with the genes belonging to the first cluster identified from the analysis of the processing rate steady state values. The network structure is indicative of the semantic similarity of the terms; i.e. linked and adjacent terms are close to each other in the reference ontology. The size of each dot is proportional to the number of genes identified in the cluster, while the color is a proxy of the significance of the enrichment which takes into account the total number of genes associated with the specific term.

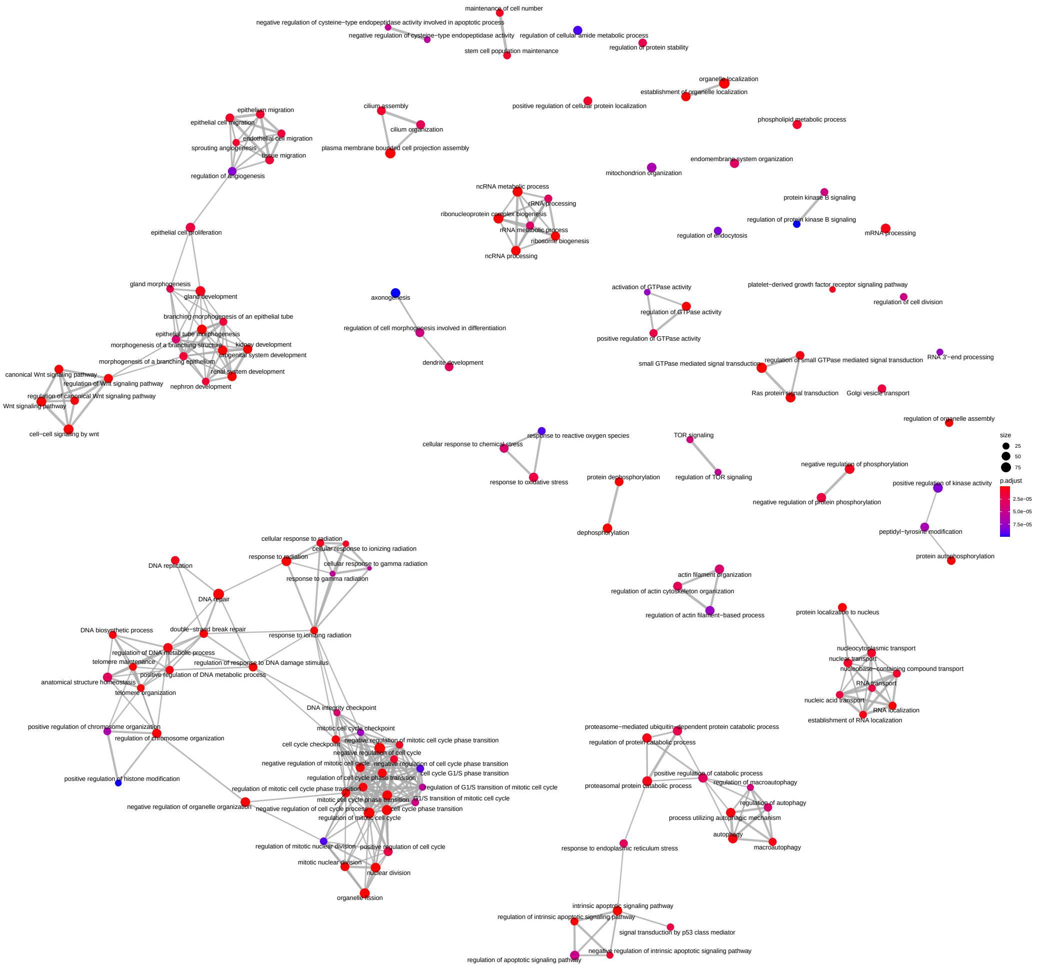


Supplemental Figure 26: **Processing steady-state rate of the Cluster B Gene Ontology analysis.** Biological processes (nodes of the network) associated with the genes belonging to the second cluster identified from the analysis of the processing rate steady state values. The network structure is indicative of the semantic similarity of the terms; i.e. linked and adjacent terms are close to each other in the reference ontology. The size of each dot is proportional to the number of genes identified in the cluster, while the color is a proxy of the significance of the enrichment which takes into account the total number of genes associated with the specific term.

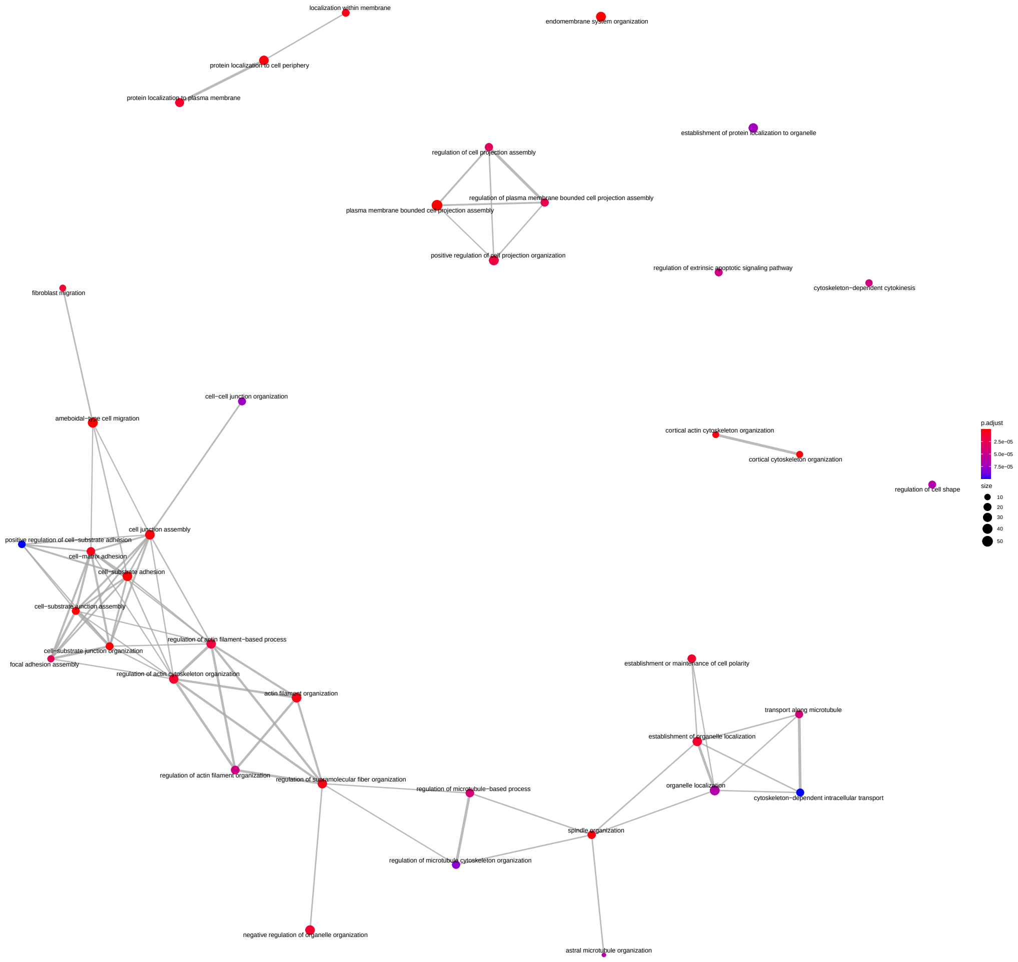


Supplemental Figure 27: **Processing rate steady state of the Cluster C Gene Ontology analysis.** Biological processes (nodes of the network) associated with the genes belonging to the third cluster identified from the analysis of the processing rate steady state values. The network structure is indicative of the semantic similarity of the terms; i.e. linked and adjacent terms are close to each other in the reference ontology. The size of each dot is proportional to the number of genes identified in the cluster, while the color is a proxy of the significance of the enrichment which takes into account the total number of genes associated with the specific term.

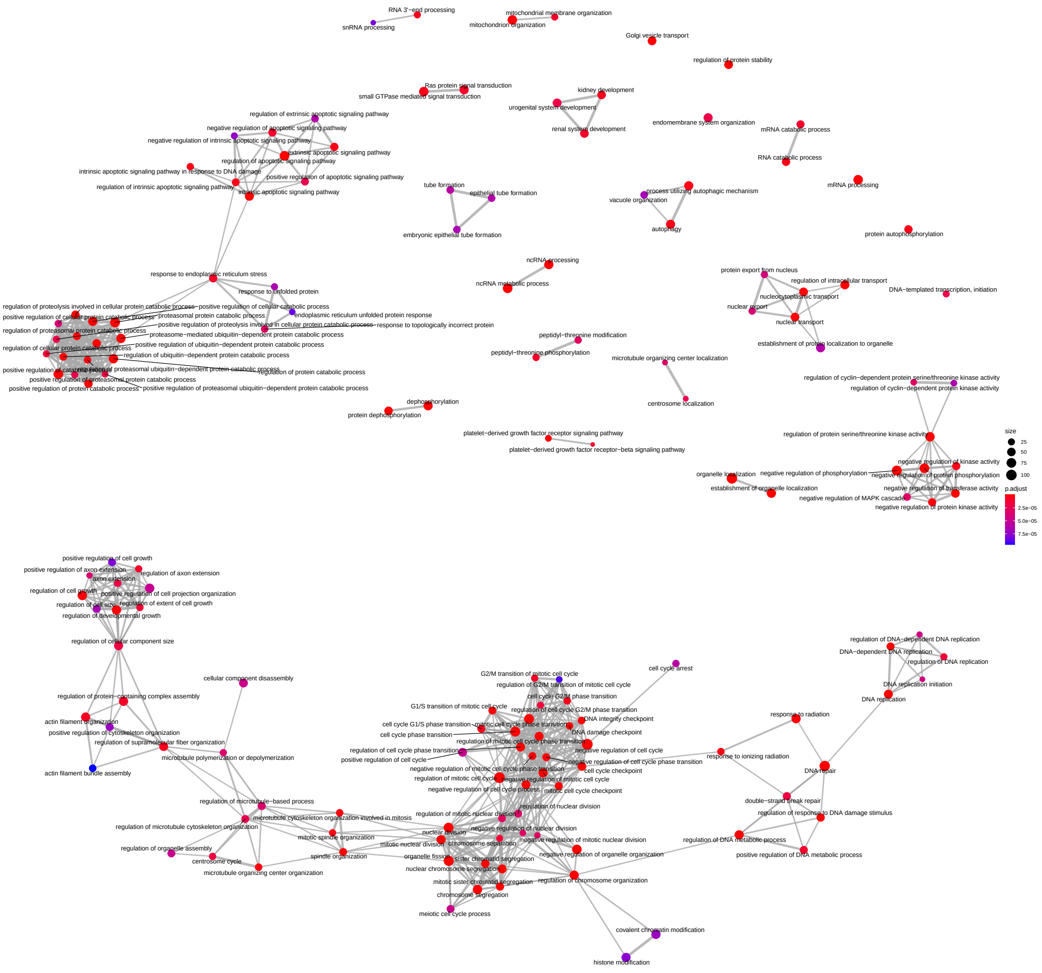


Supplemental Figure 28: **Degradation steady-state rate of the Cluster A Gene Ontology analysis.** Biological processes (nodes of the network) associated with the genes belonging to the first cluster identified from the analysis of the degradation rate steady state values. The network structure is indicative of the semantic similarity of the terms; i.e. linked and adjacent terms are close to each other in the reference ontology. The size of each dot is proportional to the number of genes identified in the cluster, while the color is a proxy of the significance of the enrichment which takes into account the total number of genes associated with the specific term.

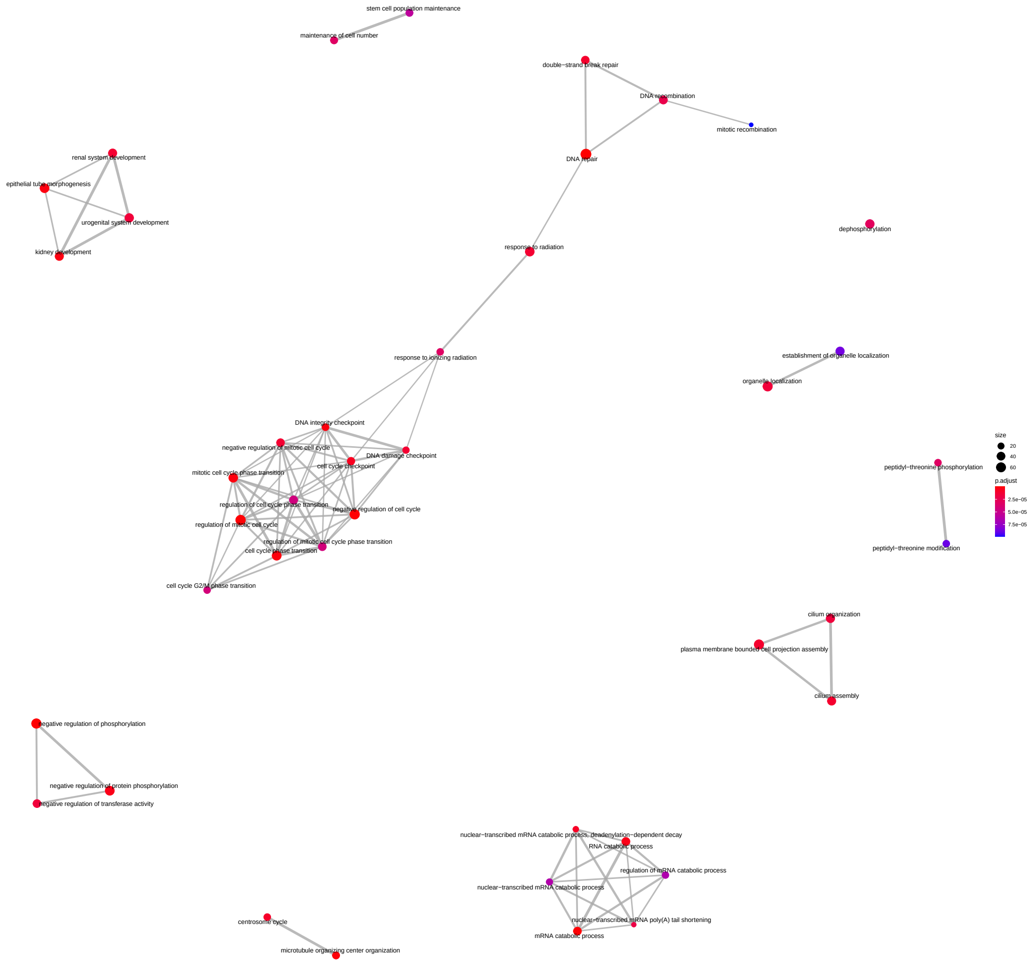


Supplemental Figure 29: **Degradation steady-state rate of the Cluster B Gene Ontology analysis.** Biological processes (nodes of the network) associated with the genes belonging to the second cluster identified from the analysis of the degradation rate steady state values. The network structure is indicative of the semantic similarity of the terms; i.e. linked and adjacent terms are close to each other in the reference ontology. The size of each dot is proportional to the number of genes identified in the cluster, while the color is a proxy of the significance of the enrichment which takes into account the total number of genes associated with the specific term.

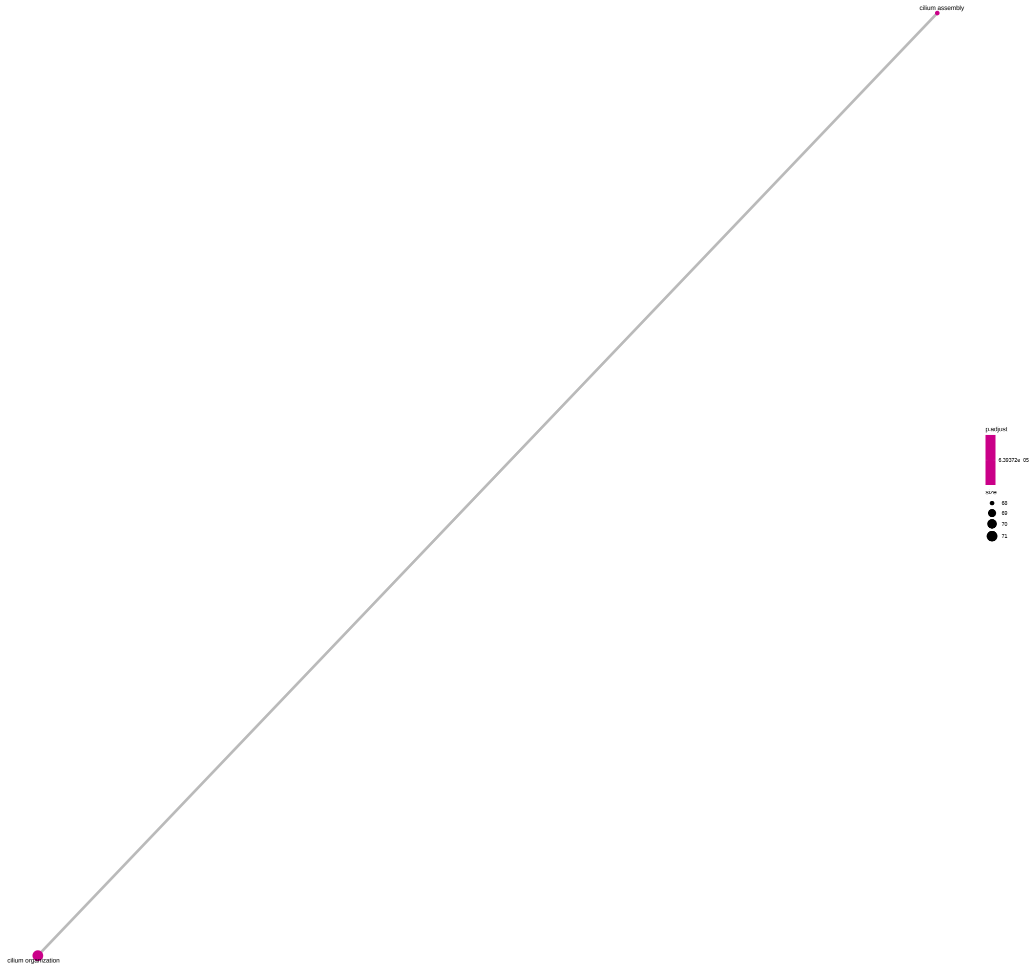


Supplemental Figure 30: **Synthesis rate steady state of the Cluster B Gene Ontology analysis on small universe.** Biological processes (nodes of the network) associated with the genes belonging to the second cluster identified from the analysis of the synthesis rate steady state values. The network structure is indicative of the semantic similarity of the terms; i.e. linked and adjacent terms are close to each other in the reference ontology. The size of each dot is proportional to the number of genes identified in the cluster, while the color is a proxy of the significance of the enrichment which takes into account the total number of genes associated with the specific term among the 4897 used for the analysis.

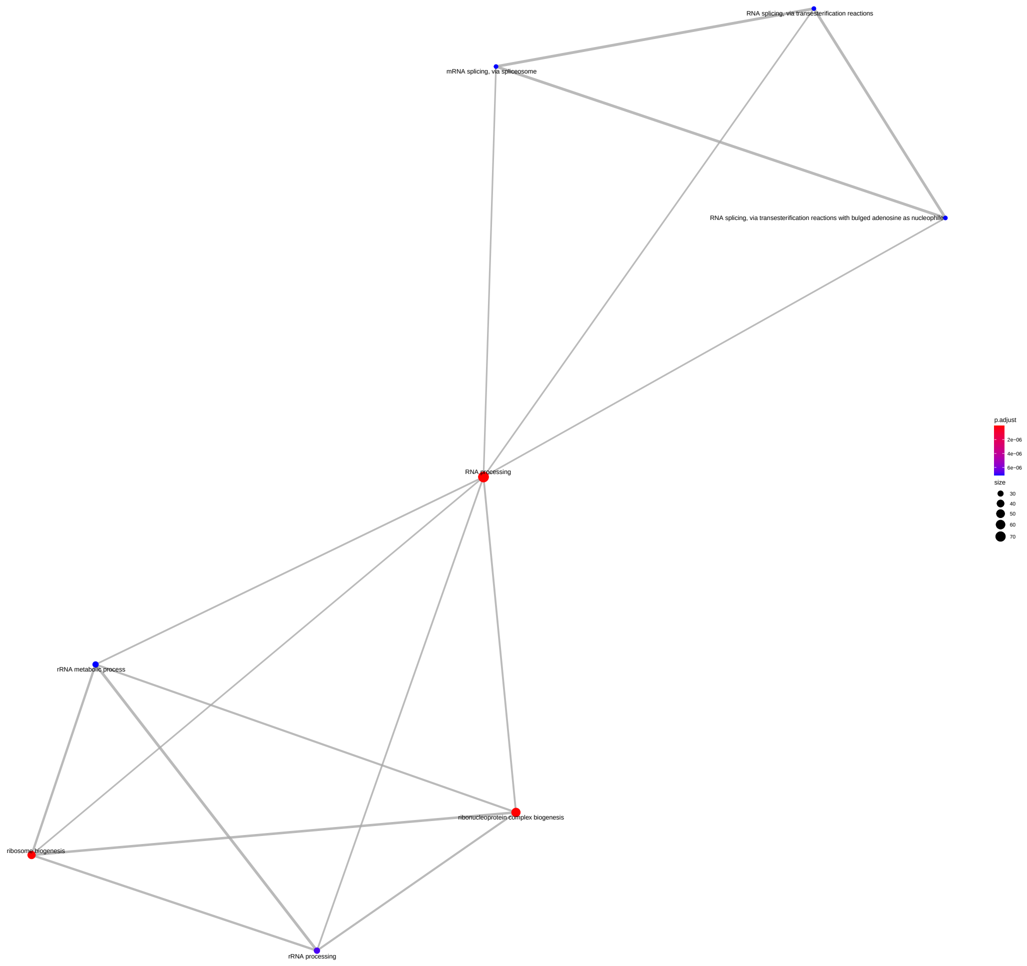


Supplemental Figure 31: **Synthesis rate steady state of the Cluster C Gene Ontology analysis on small universe.** Biological processes (nodes of the network) associated with the genes belonging to the third cluster identified from the analysis of the synthesis rate steady state values. The network structure is indicative of the semantic similarity of the terms; i.e. linked and adjacent terms are close to each other in the reference ontology. The size of each dot is proportional to the number of genes identified in the cluster, while the color is a proxy of the significance of the enrichment which takes into account the total number of genes associated with the specific term among the 4897 used for the analysis.

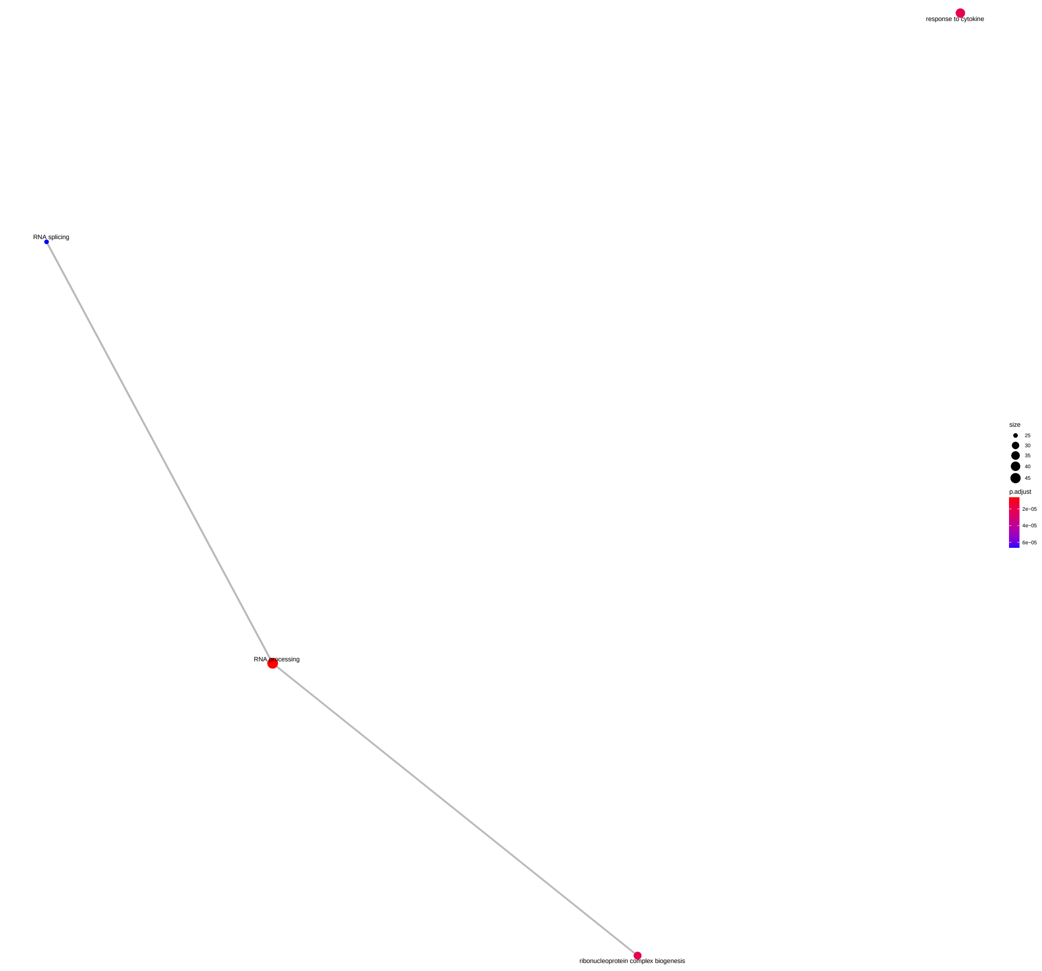


Supplemental Figure 32: **Synthesis rate steady state of the Cluster D Gene Ontology analysis on small universe.** Biological processes (nodes of the network) associated with the genes belonging to the fourth cluster identified from the analysis of the synthesis rate steady state values. The network structure is indicative of the semantic similarity of the terms; i.e. linked and adjacent terms are close to each other in the reference ontology. The size of each dot is proportional to the number of genes identified in the cluster, while the color is a proxy of the significance of the enrichment which takes into account the total number of genes associated with the specific term among the 4897 used for the analysis.

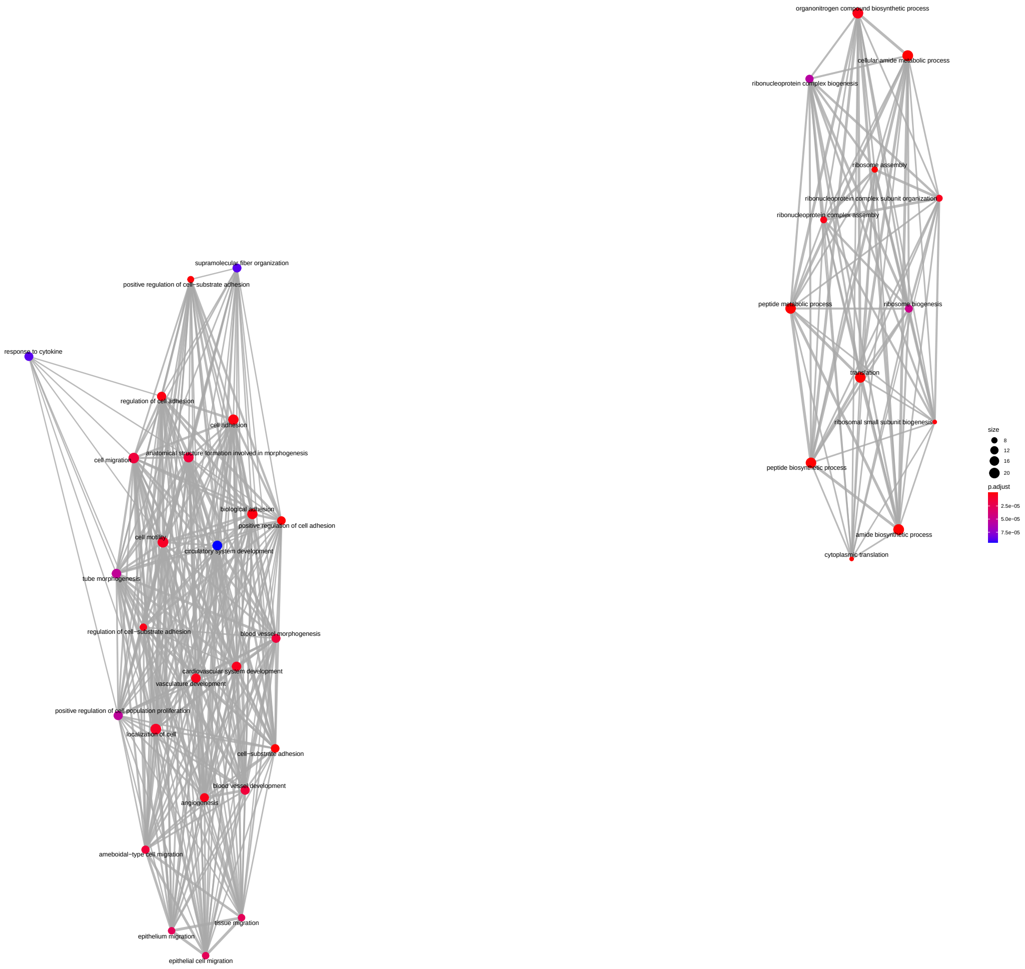


Supplemental Figure 33: **Synthesis rate steady state of the Cluster F Gene Ontology analysis on small universe.** Biological processes (nodes of the network) associated with the genes belonging to the sixth cluster identified from the analysis of the synthesis rate steady state values. The network structure is indicative of the semantic similarity of the terms; i.e. linked and adjacent terms are close to each other in the reference ontology. The size of each dot is proportional to the number of genes identified in the cluster, while the color is a proxy of the significance of the enrichment which takes into account the total number of genes associated with the specific term among the 4897 used for the analysis.

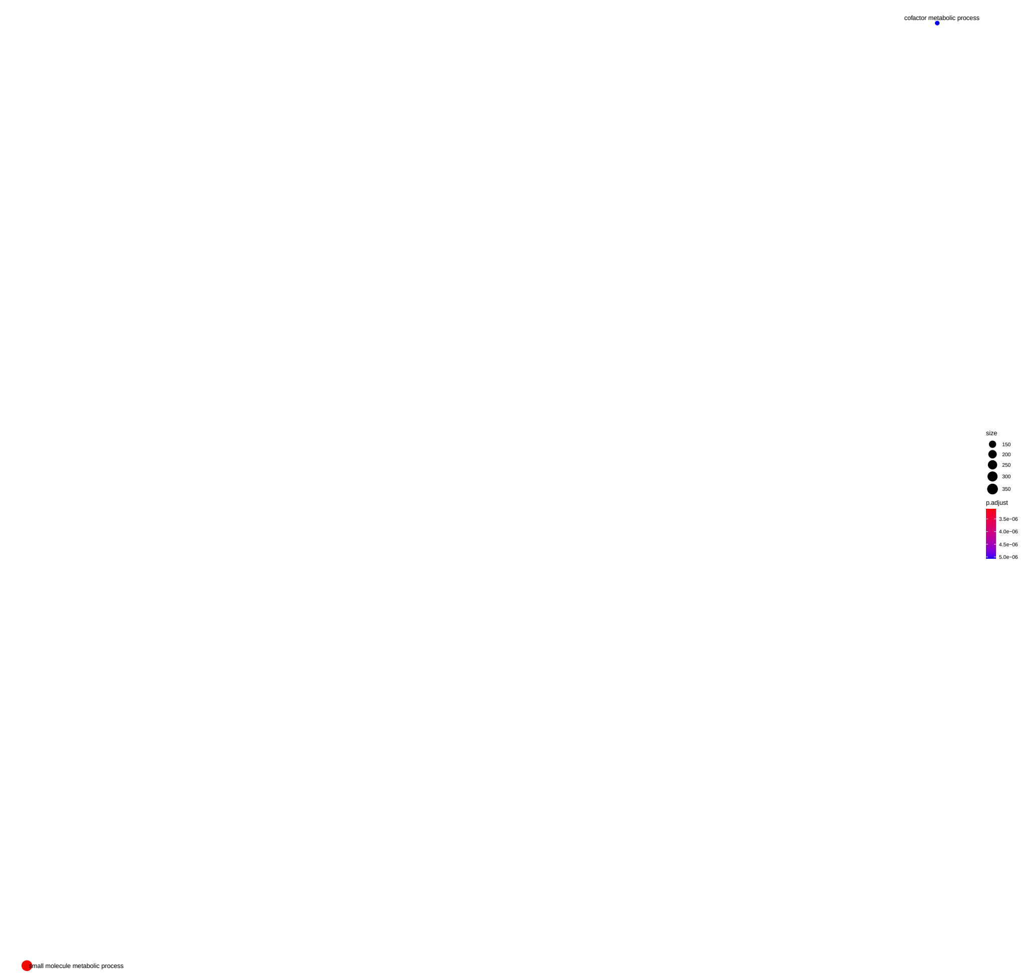


Supplemental Figure 34: **Degradation rate steady state of the Cluster A Gene Ontology analysis on small universe.** Biological processes (nodes of the network) associated with the genes belonging to the first cluster identified from the analysis of the degradation rate steady state values. The network structure is indicative of the semantic similarity of the terms; i.e. linked and adjacent terms are close to each other in the reference ontology. The size of each dot is proportional to the number of genes identified in the cluster, while the color is a proxy of the significance of the enrichment which takes into account the total number of genes associated with the specific term among the 4897 used for the analysis.

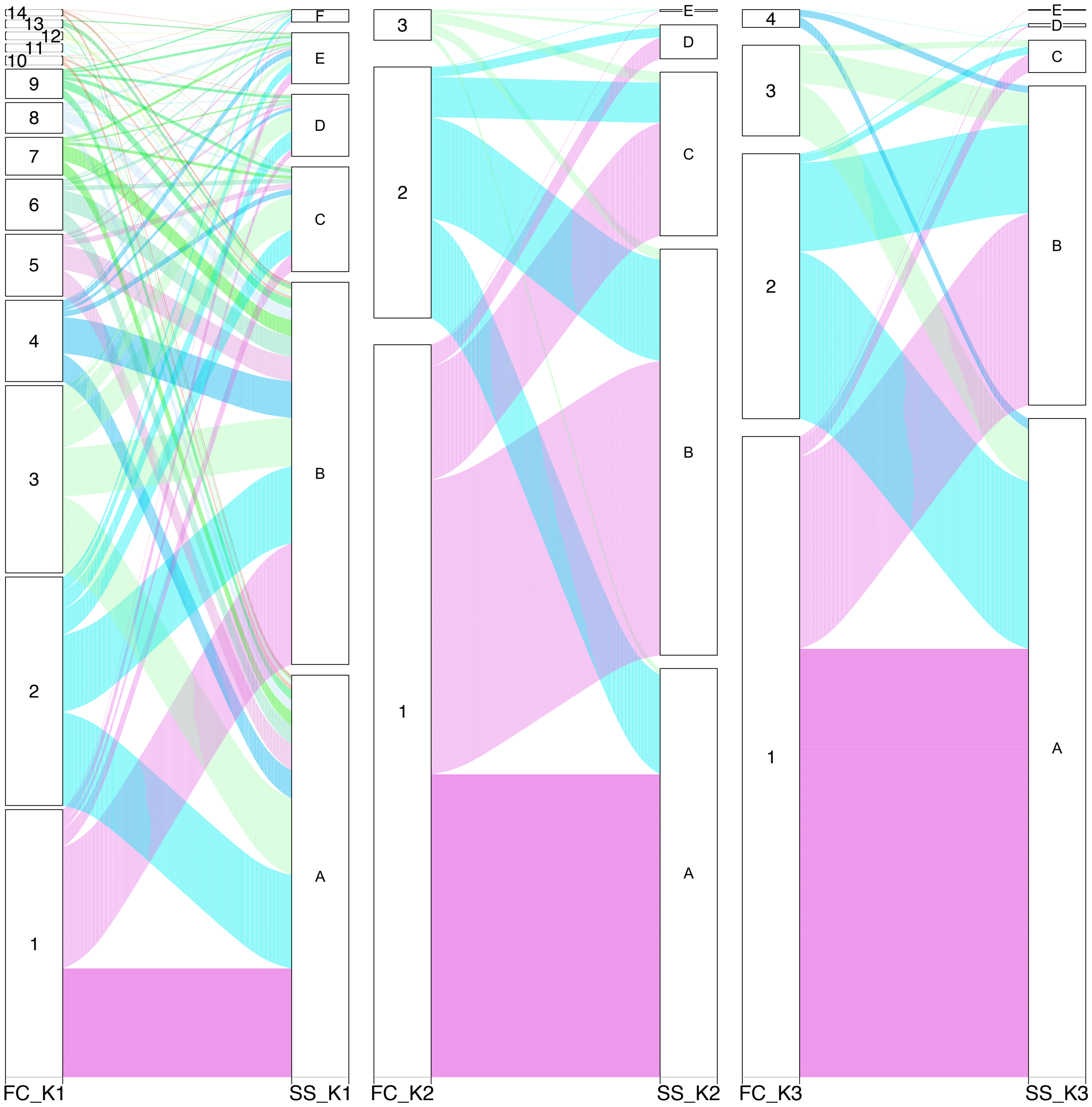


Supplemental Figure 35: **Alluvial plot.** Alluvial plots showing for synthesis, processing and degradation rates the flux of elements from the clusters defined on the rate modulations (FC), to clusters defined on its steady state values (SS).

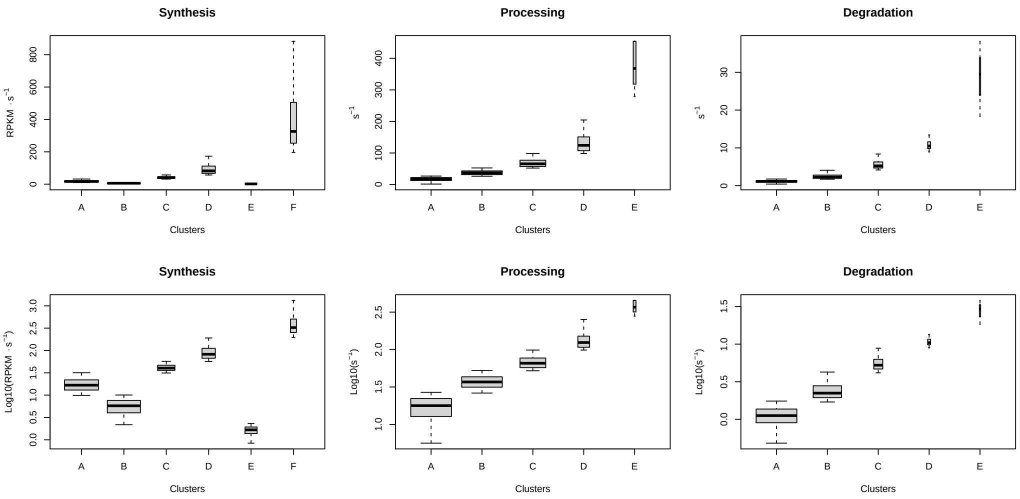


Supplemental Figure 36: **Steady state cluster distributions.** Boxplots showing the linear (top row) and logarithmic (bottom row) distributions of the synthesis, processing and degradation rate steady-state values clustered according to the corresponding mixture models. The sizes of the boxes, for each rate, are proportional to the number of elements in the clusters.

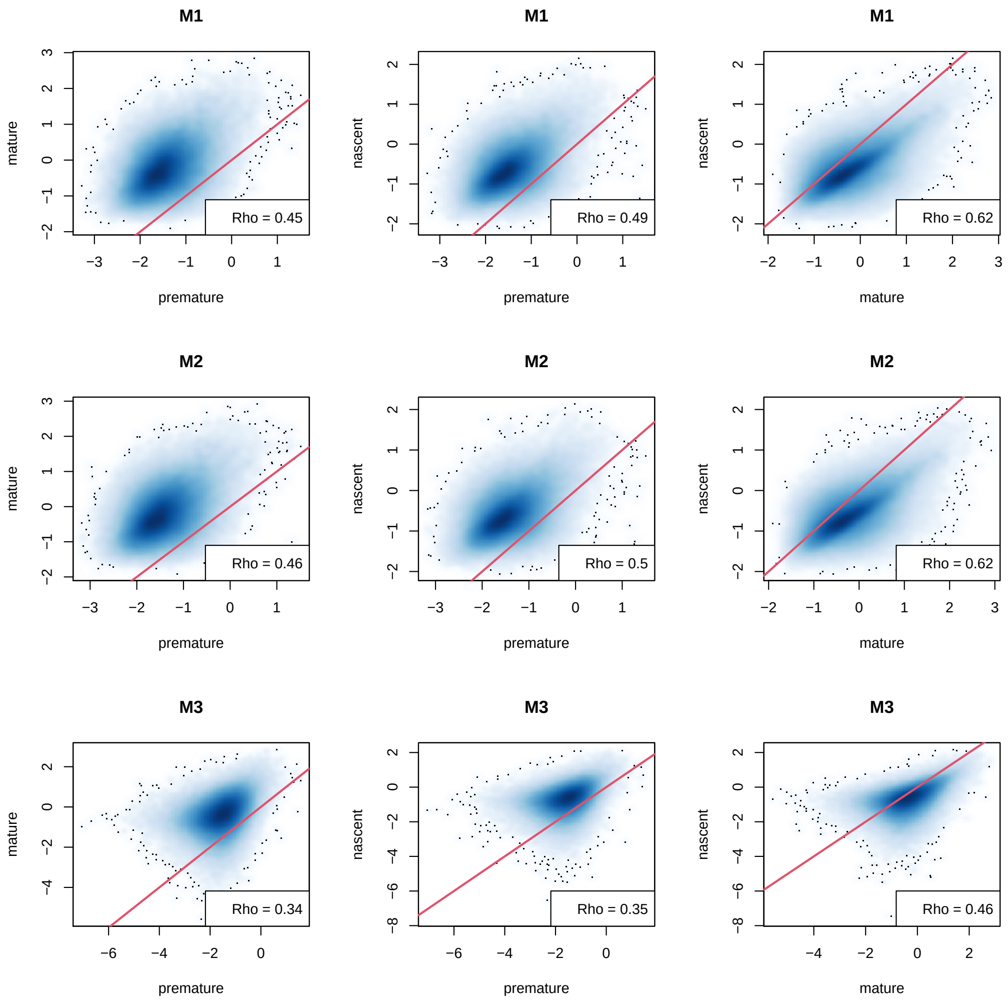


Supplemental Figure 37: **CRPS coefficient analysis.** Smooth-scatter plots comparing CRPS indexes obtained from M1 (top row), M2 (middle row) and M3 (bottom row) for nascent, premature and mature RNA. Red solid lines represent the bisector of the first and third quadrants. Rho is the Pearson's correlation coefficient estimated for each cloud of data.

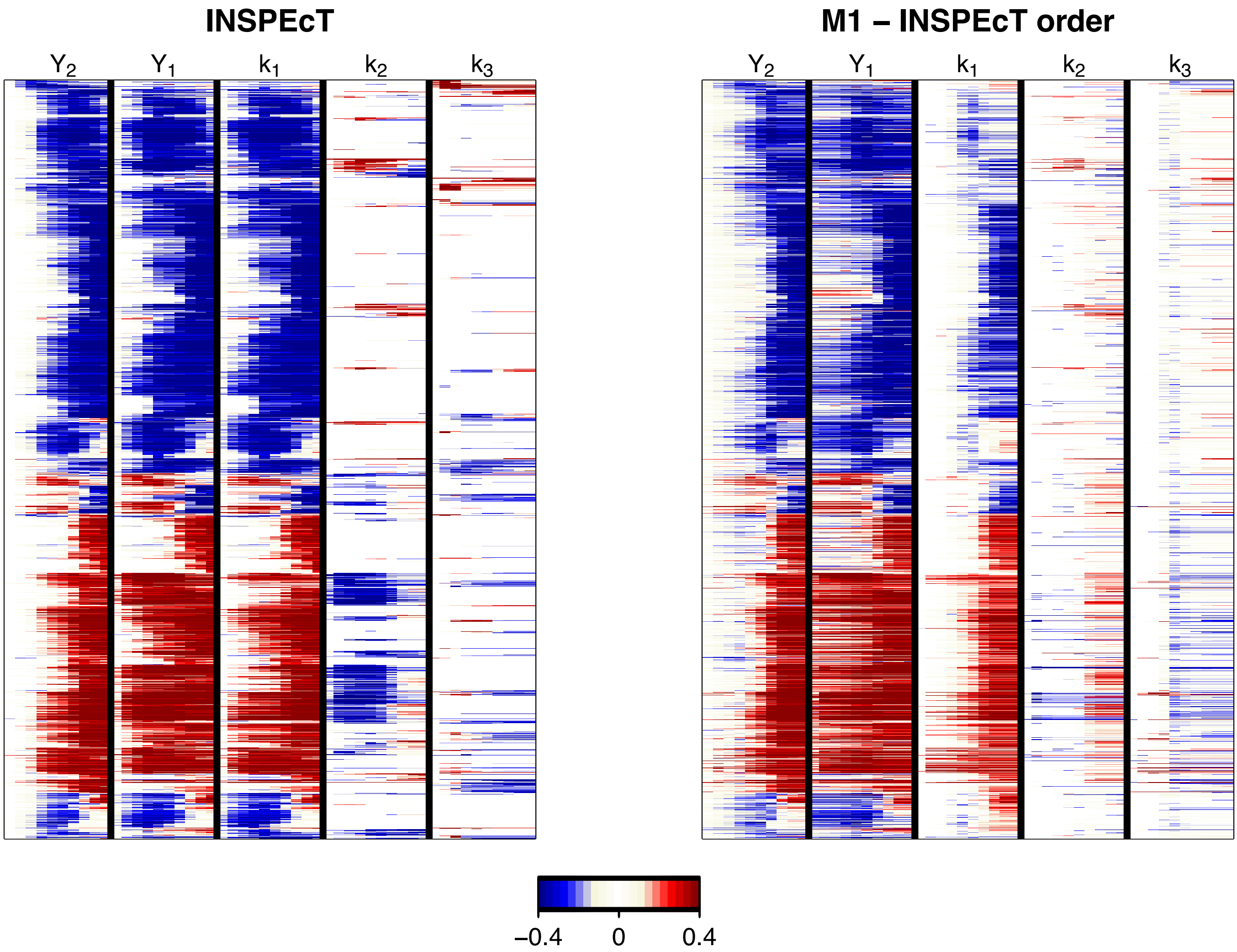


Supplemental Figure 38: **Comparison of the M1 and M3 models.** Heatmaps of M1 and M3 showing the log2-fold-changes, estimated using the initial steady-state values as baselines, of: the modelled mature RNA ($Y_2$), the modelled premature RNA ($Y_1$), synthesis ($k_1$), processing ($k_2$) and degradation ($k_3$) rates. The order of the genes (rows) is obtained by clustering the M3 responses and the same order is preserved in the two heatmaps.
